# Upregulation of a novel lncRNA KLHDC7B-DT in Head and Neck Cancers: Implications for Prognosis and Molecular Mechanisms

**DOI:** 10.64898/2025.11.30.691487

**Authors:** Subhayan Sur, Dimple Davray, Soumya Basu, Samir Gupta, Supriya Kheur, B. M. Rudagi

## Abstract

Head and neck cancer (HNC), the seventh most frequent malignancy globally, is a serious health concern. Despite advancements in treatment, the overall survival rate for HNC remains around 50%, with even poorer outcomes in metastatic cases. This highlights the urgent need to explore the molecular mechanisms underlying HNC for improved diagnostics and therapies. The importance of non-coding RNAs, especially long non-coding RNAs (lncRNAs), in a variety of malignancies, including HNC, has been highlighted in recent researches. Although lncRNAs and mRNA are very similar, lncRNAs do not encode proteins and they are longer than 200 nucleotides. While studies have implicated lncRNAs in several cancers, including HNC, their exact role in HNC remains unclear, and there are currently no clinically established lncRNA biomarkers for diagnosis or therapy. Thus, this study aims to identify novel lncRNAs and investigate their significance in HNC. For this, we looked over recent research and discovered a new lncRNA called KLHDC7B-DT. The KLHDC7B-DT is significantly upregulated in HNC patient samples, and this result is corroborated by TCGA HNC data and cell lines. Further investigation suggests that KLHDC7B-DT may be linked with cancer stage, grade, age, and worse patient survival. We identified potential miRNAs that interact with KLHDC7B-DT and presented mRNA targets of these miRNAs, along with their possible functions. We propose that KLHDC7B-DT may regulate IL-6 expression in HNC, either through miRNAs or alternative mechanisms. Additionally, we found a significant positive correlation between the neighboring protein-coding gene KLHDC7B and the KLHDC7B-DT in HNC, both of which are significantly upregulated in HNC. In conclusion, KLHDC7B-DT is upregulated in HNC and holds promise as a prognostic and therapeutic biomarker. This study marks the first report highlighting the significance of KLHDC7B-DT in HNC.

## I. INTRODUCTION

Head and neck cancer (HNC) is the seventh most frequent cancer worldwide and a major public health concern in India, making it a serious global health challenge. Each year, over 660,000 new cases and approximately 325,000 deaths are attributed to HNC, highlighting its profound impact on public health [1]. In India, the age-standardized incidence rate for HNC is 25.9 per 100,000 for males and 8.0 per 100,000 for females [2]. HNC comprises malignancies that mostly originate from the mucosal epithelium and affect the oral cavity, pharynx, and larynx. According to estimates, oral cavity and pharyngeal cancers will cause 12,230 fatalities and 58,450 new cases in the US in 2024 [3].

A number of risk factors, such as excessive alcohol usage, tobacco use, and chewing betel nut, affect the prevalence of HNC [4, 5]. In addition, human papillomavirus (HPV) infection is strongly linked to oropharyngeal cancers and global vaccination efforts are expected to reduce the incidence of HPV-positive HNC [5]. Despite a 33% decrease in the overall cancer death rate from 1991 to 2021, mortality due to the cancers in the tongue, tonsils, and oropharynx continue to rise by 2% each year in the U.S. [3].

A multimodal strategy, involving radiation therapy, chemotherapy, and surgery, is frequently necessary for the management of HNC, particularly when it is in an advanced stage. A monoclonal antibody called cetuximab, which targets the epidermal growth factor receptor (EGFR), has FDA approval and is used to sensitize patients to radiotherapy for recurrent or metastatic HNC. For the treatment of cisplatin-refractory recurrent or metastatic oral squamous cell carcinoma (OSCC), checkpoint inhibitors of immune system like pembrolizumab and nivolumab have also received approval [5]. However, the overall survival rate for HNC is still about 50%, and the prognosis for metastatic patients is much worse, even with these improvements [6]. The molecular mechanisms of HNC must thus be better understood in order to pinpoint more effective diagnostic and treatment targets.

The importance of non-coding RNAs, especially long non-coding RNAs (lncRNAs), in the development of many malignancies, including HNC, has been brought to light by recent studies [7, 8]. RNA polymerase II transcribes long noncoding RNAs (lncRNAs), which are more than 200 nucleotides long and are essential for controlling gene expression and cellular functions [7, 8]. About 3,000 lncRNAs are linked to different diseases in the “LncRNADisease” database; several of them, including HOTAIR, MALAT-1, and H19, exhibit dysregulation in cancerous settings [9, 10]. The diagnostic and therapeutic potential of these lncRNAs in different malignancies is not being investigated in many clinical trials [7, 8]. These investigations highlight the increasing understanding of lncRNAs as useful targets for cancer treatment and diagnosis. Nevertheless, nothing is known about the functions of lncRNAs in HNC, and no lncRNA biomarkers have been created especially for this kind of cancer.

In this study, we present, for the first time, the involvement of the lncRNA KLHDC7B-DT (CTA-384D8.35 or Ensembl: ENSG00000272666) in HNC. This relatively novel lncRNA has only a few publications detailing its role in human diseases including cancer [11, 12]. Here, we report its significant upregulation in HNC and explore its potential functional and prognostic implications. While these findings are preliminary, they open the door for further mechanistic studies and provide a foundation for future investigations in larger preclinical and clinical settings.

## II. MATERIALS AND METHODS

### A. Collection of patient samples

With the appropriate consent, 20 HNC patient samples and 20 adjacent non-tumor tissues were obtained from Dr. D. Y. Patil Dental College & Hospital and Dr. D. Y. Patil Medical College, Hospital & Research Centre, Dr. D. Y. Patil Vidyapeeth (DPU), Pune (**Table 1**). This study was approved by the Institutional Biosafety Committee (IBSC) and Ethics Committee.

**Table 1:** Demography of Patient Sample.

| Parameters |  | Number |
| --- | --- | --- |
| Total |  | 20 |
| Age | <40 yrs | 2 |
|  | 40-50 yrs | 11 |
|  | >50 yrs | 7 |
| Sex | Male | 19 |
|  | Female | 1 |
| Habit | Tobacco | 17 |
|  | Alcohol | 10 |
| Cancer grade | Well | 8 |
|  | Moderate | 8 |
|  | Poor | 4 |
| HPV status | Positive | 0 |

### B. RNA isolation and lncRNA expression analysis

The tissue’s total ribonucleic acid was extracted by TRIzol reagent (Invitrogen, CA). The iScript cDNA Synthesis Kit (BioRad) was used for making cDNA as per manufacturer’s guidelines. The iTaq Universal SYBR Green Supermix (BioRad) was utilized for real-time PCR (RT-PCR) in accordance with the manufacturer’s instructions for gene expression analysis. FP: 5’ CAGTTTCCCATGAATGATGC 3’ and RP: 5’ CAGGTGTTCCCCAATTCATA 3’ were the primers for expression analysis of KLHDC7B-DT. Endogenous control was achieved using 18S rRNA (FR: 5’ GTCATAAGCTTGCGTTGATT 3’ and RP: 5’ TAGTCAAGTTCGACCGTCTT 3’). The gene’s fold change was computed as 2^(-ΔΔCt), as previously mentioned [13]. A triple of each sample was loaded.

### C. Bioinformatics and statistical analysis

The KLHDC7B-DT (CTA-384D8.35 or Ensembl: ENSG00000272666) gene alteration, gene expression, and patient survival were analyzed from TCGA Head and neck squamous cell carcinoma dataset using cBioPortal (www.cbioportal.org) and GEPIA (Tumor: 519 and normal: 44; http://gepia.cancer-pku.cn) platforms. Gene expression cut-off criteria were |Log2FC|: 1, p-value: 0.01, and median value with levels above it considered high expression and levels below it considered low expression.

To ensure precise comparisons, a total of 42 paired tumors and matched normal samples were selected from the TCGA Head and neck squamous cell carcinoma dataset using a count matrix retrieved from the TCGA database by TCGA-Biolinks R package. DEG analysis was conducted using R-based statistical tools like DESeq2 identifying significantly upregulated and downregulated genes based on thresholds (adjusted p_value less than 0.05 and log2 fold-change > 1 or < −1).

Expressions and copy numbers of the gene in human HNC cell lines were analysed using EMBL-EBI: Expression Atlas (https://www.ebi.ac.uk/) and DepMap portal (https://depmap.org/). Correlation analysis was performed using GEPIA from the TCGA data and Excel from the studied samples [14]. Interaction between the KLHDC7B-DT transcript (ENST00000796180.1) and miRNAs was predicted using miRDB tools (mirdb.org/mirdb/index.html).

The lncRNA expression data (fold change) were analysed by Student’s t-test. Statistical significance was calculated as a p value of less than 0.05. The means ± standard deviations are used to display the representative data. The representative data are shown as means ± standard deviations.

## III. RESULTS AND DISCUSSION

### A. LncRNA KLHDC7B-DT is upregulated in TCGA head and neck cancer (HNC) samples

KLHDC7B divergent transcript (KLHDC7B-DT) is relatively newly discovered lncRNA and is found to be upregulated in pancreatic ductal adenocarcinoma [11]. The KLHDC7B-DT modifies pancreatic ductal adenocarcinoma microenvironment through interaction and transcriptional activation of IL-6 [11]. The modification of tumor microenvironment is also a major event in HNC progression, metastasis, and drug resistance [15, 16]. These observations provoked us to find the status of the new lncRNA KLHDC7B-DT in HNC.

The lncRNA KLHDC7B-DT (Ensembl: ENSG00000272666 or CTA-384D8.35) is located in chr. 22q13.33, positioned near the Kelch Domain Containing 7B (KLDHC7B) on the opposite strand (Fig. 1A). The KLHDC7B-DT is significantly upregulated in the TCGA HNC samples as compared to the normal tissues (Tumor: 519 and normal: 44; GEPIA) (Fig. 1B). To achieve a more precise comparison, we extracted 42 paired HNC samples along with their corresponding normal counterparts from the TCGA dataset. Similarly, KLHDC7B-DT is significantly upregulated in HNC compared to normal tissues (Log2FC: 2.819688579 and P value 3.7×10-15) (Fig. 1C). The lncRNA expression is consistently upregulated in all the cancer stages (Fig. 1D). All these data indicate that the lncRNA KLHDC7B-DT is upregulated HNC.

**Fig. 1.**
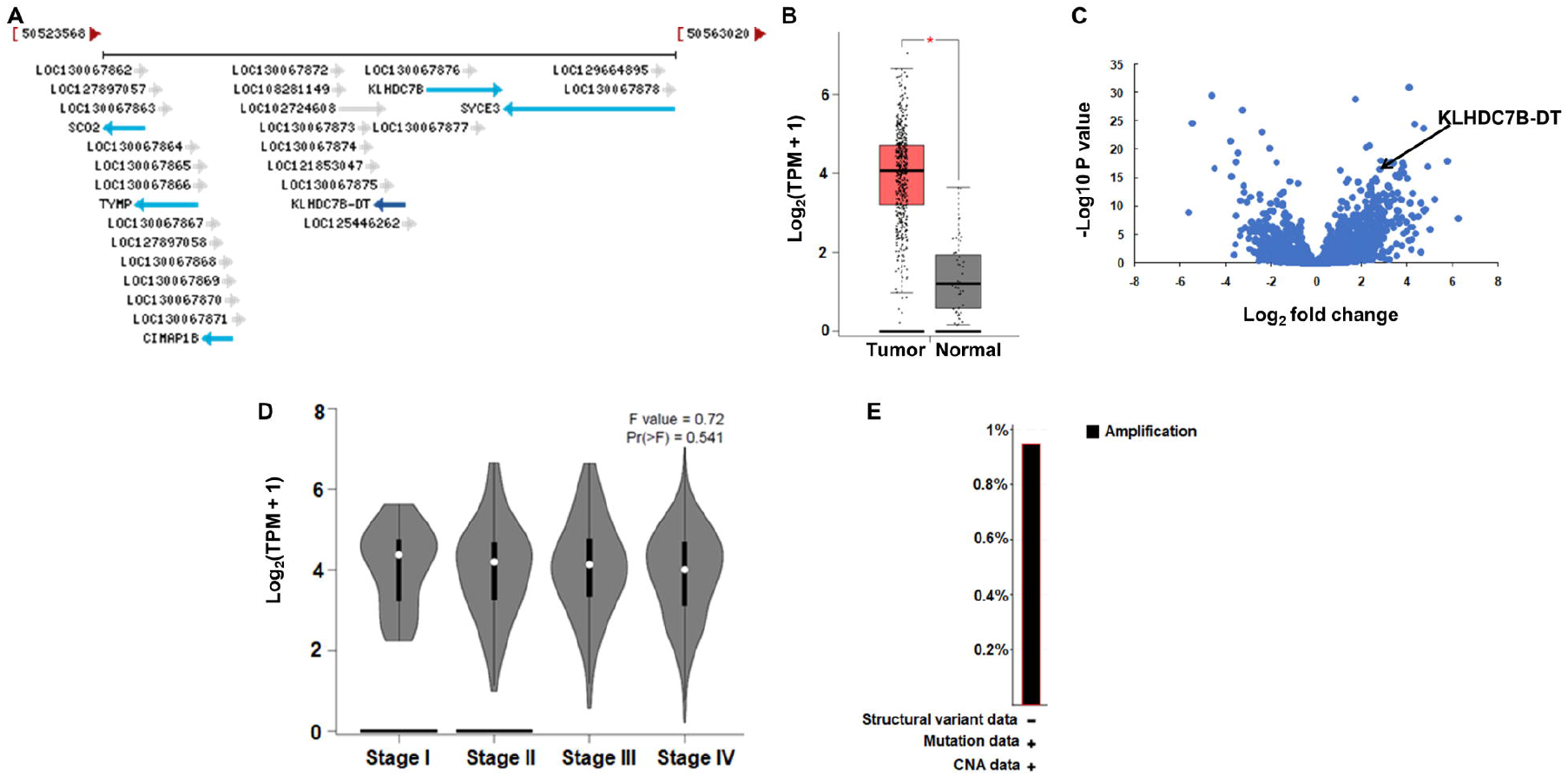
Overexpression of LncRNA KLHDC7B-DT in HNC. **A:** Genomic region of the KLHDC7B-DT gene on chromosome 22q, illustrating neighbouring genes. **B:** KLHDC7B-DT is significantly upregulated in TCGA HNC samples (n=519) compared to normal samples (n=44), as analyzed by GEPIA (*P < 0.01). **C:** Volcano plot showing the expression of lncRNAs (Log2 fold change) against –Log10 P-value, in 42 paired TCGA HNC samples. **D:** Expression levels of KLHDC7B-DT across different stages of HNC were analysed using GEPIA. **E:** Genetic alterations in KLHDC7B-DT detected in 5 out of 528 cases (amplification in 0.95%) in Head and Neck Squamous Cell Carcinoma (TCGA, GDC), analyzed by cBioPortal..

Next, we sought to understand why KLHDC7B-DT is upregulated in HNC samples. However, we found no supporting literature. The Head and Neck Squamous Cell Carcinoma (TCGA, GDC) data showed that 5 out of 528 patients (0.95%) had gene amplification (Fig. 1E). Therefore, more investigation is needed to determine the fundamental reasons behind KLHDC7B-DT overexpression in HNC.

### B. KLHDC7B-DT is involved in poor patient survival

To evaluate the prognostic significance of KLHDC7B-DT in HNC, we analyzed patient survival in relation to its genetic alterations and expression using TCGA data. Although the number of samples with KLHDC7B-DT amplification is limited, the trend suggests poorer overall survival in the altered group compared to the unaltered population (Fig. 2A). Furthermore, in TCGA HNC data, higher KLHDC7B-DT expression is linked to worse_overall and disease_free_survival (Fig. 2B and C) (Fig. 2B and C). Notably, KLHDC7B-DT is also associated with poor patient survival in pancreatic ductal adenocarcinoma, underscoring its potential prognostic role in cancers [11].

**Fig. 2.**
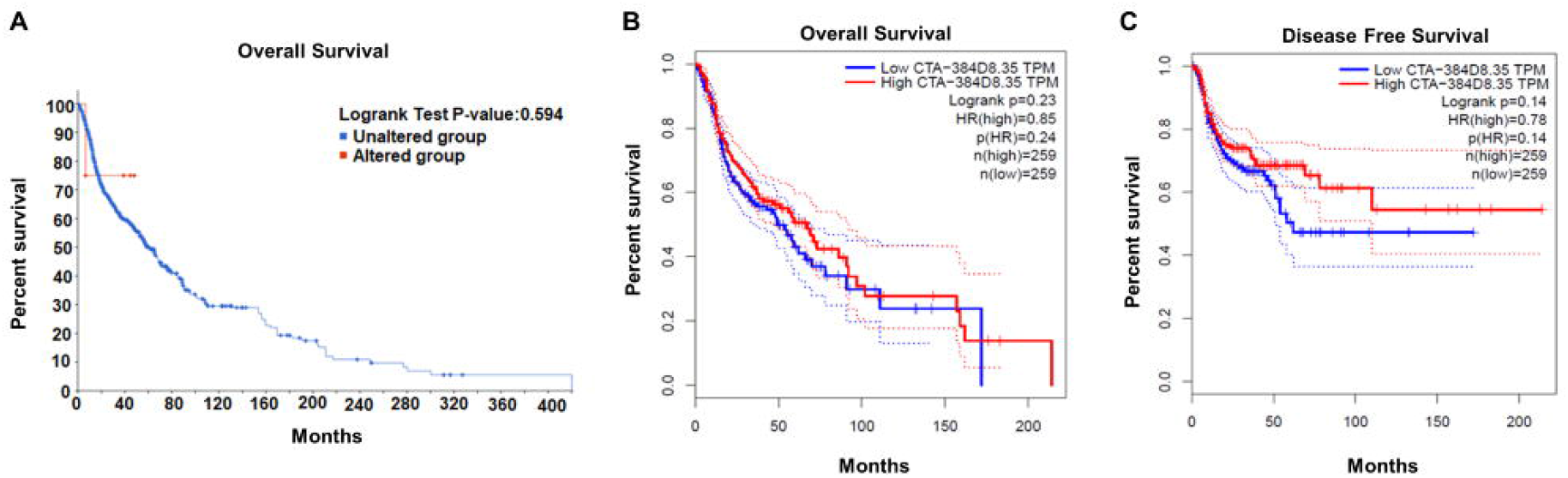
Prognostic Role of KLHDC7B-DT in HNC Survival. **A:** Impact of KLHDC7B-DT gene alteration on overall survival in TCGA HNC patients) analyzed by cBioPortal. **B:** The relationship between KLHDC7B-DT expression levels and overall_survival in patients with TCGA HNC anlyzed by GEPIA. **C:** The relationship between KLHDC7B-DT expression and disease-free survival in TCGA HNC patient samples.

Few studies have investigated the role of lncRNAs in the diagnosis and prognosis of HNC [7, 17-19], and only fewer have been evaluated in clinical trials. But no lncRNAs have yet been established as reliable biomarkers for HNC. Therefore, further research is needed to identify new lncRNAs and evaluate their roles in larger patient cohorts, which could lead to the development of effective lncRNA-based diagnostic and prognostic tools for better disease management.

### C. Expression and copy numbers of KLHDC7B-DT in HNC cell lines

To understand the mechanistic role of lncRNAs, it is essential to investigate their functions in preclinical settings using cancer cell lines. Therefore, assessing the expression and copy number of KLHDC7B-DT in established HNC cell lines is crucial. Comparatively, higher expression levels of KLHDC7B-DT are observed in the YD-10B, SCC-15, Cal27, BICR 56, and BICR 22 HNC cell lines (Fig. 3A). The average Log2 copy number of KLHDC7B-DT across different HNC cell lines is 0.95, with the highest Log2 copy number of 1.89 observed in the FADU cell line (Fig. 3B).

**Fig. 3.**
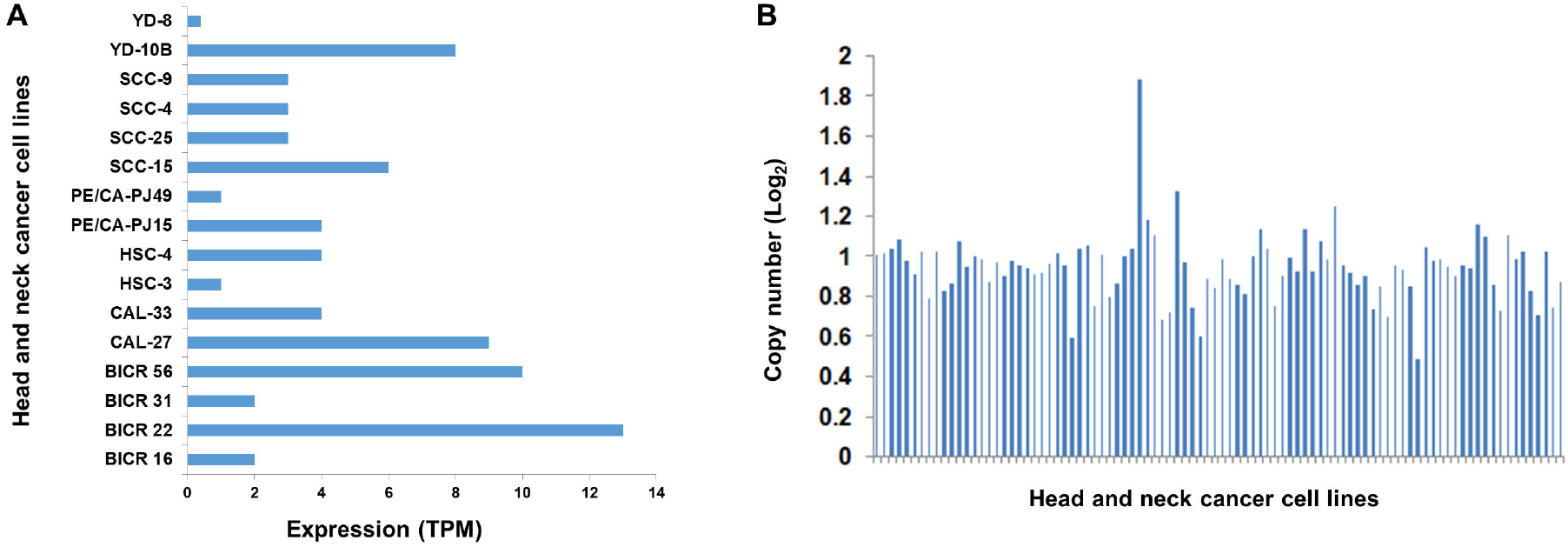
Expression and Copy Number of KLHDC7B-DT in HNC Cell Lines. **A:** Expression levels of KLHDC7B-DT (TPM) across various HNC cell lines, analyzed using the EMBL-EBI Expression Atlas. **B:** Log2 copy number (relative to ploidy +1) of KLHDC7B-DT in HNC cell lines, assessed through the Dependency Map (DepMap) portal (Copy Number Public 24Q4).

### D. Validation of KLHDC7B-DT expression by RT-PCR in HNC patient samples

We next validated the expression of KLHDC7B-DT in HNC patient samples using RT-PCR. A total of 20 tumor samples, along with adjacent non-tumor tissues, were collected. The patient demographics are detailed in Table 1. The cancerous and non-cancerous tissue types, as well as tumor grading, were confirmed through hematoxylin and eosin staining (data not shown). Consistent with TCGA data, our RT-PCR results demonstrated significant upregulation of KLHDC7B-DT in HNC patient samples compared to adjacent non-tumor tissues (Fig. 4A), validating KLHDC7B-DT overexpression in HNC.

**Fig. 4.**
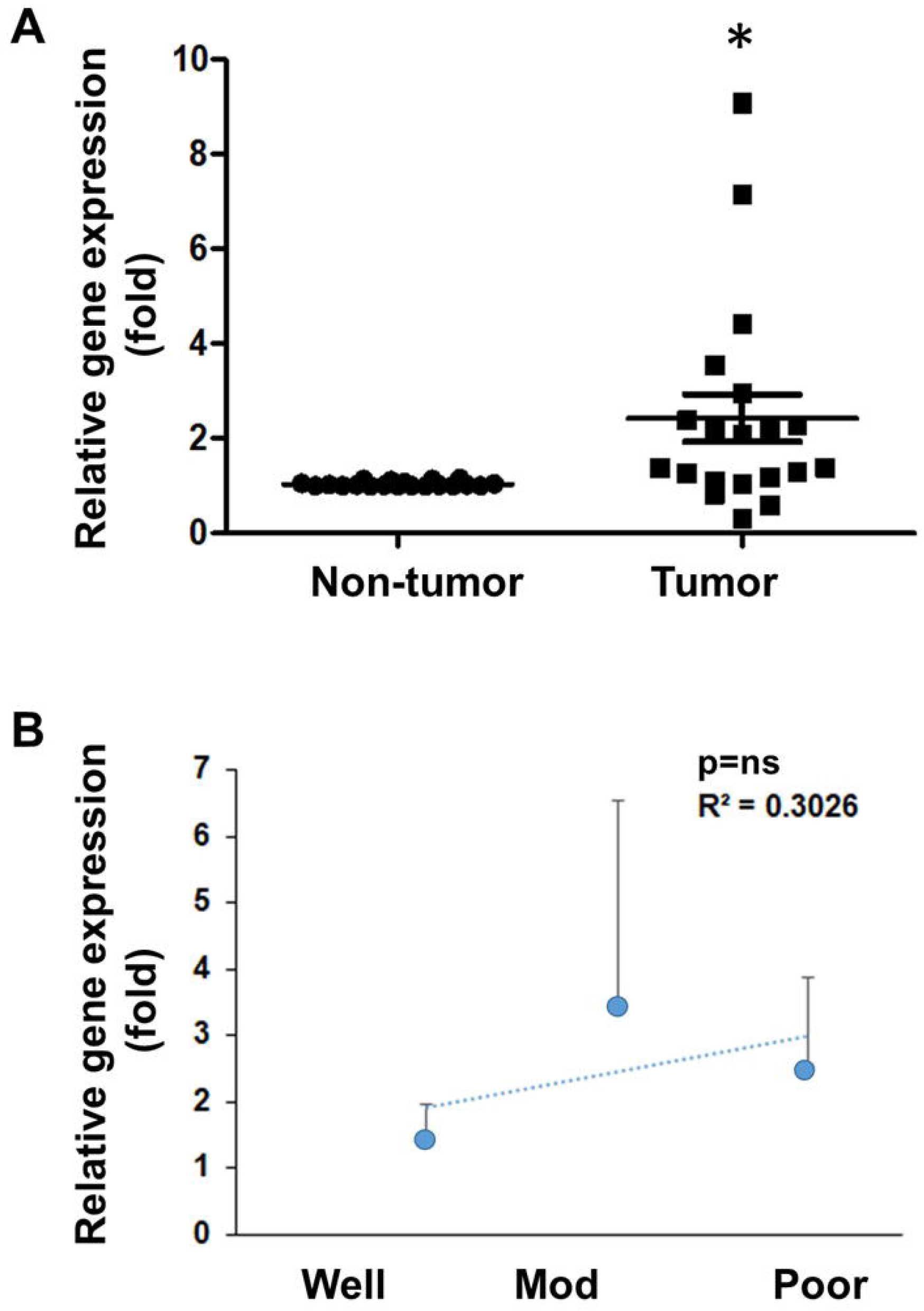
Validation of KLHDC7B-DT Expression by Real-Time PCR. **A:** KLHDC7B-DT expression (fold change) is significantly upregulated in HNC patient samples (n=20) than adjacent non-tumor tissue (*P < 0.05). **B:** Expression levels of KLHDC7B-DT across different grades of HNC: Well_differentiated _carcinoma (sample=8, Well), moderately_differentiated_carcinoma (sample=8; Mod), and poorly_differentiated_carcinoma (sample=4; Poor). The Pearson correlation (R-value) shows expression trends across groups (ns: not significant).

Despite the small sample size, we attempted to correlate KLHDC7B-DT expression with cancer grade and patient age. The analysis revealed a positive association between KLHDC7B-DT expression and cancer grade progression (Fig. 4B), as well as with age groups (data not shown). However, since nearly all subjects were HPV-negative and had a history of tobacco and alcohol use, and only one female subject was obtained, we were unable to correlate KLHDC7B-DT expression with these clinical parameters. Therefore, to gain a clearer understanding, further investigation with a larger patient cohort is necessary for a more robust analysis of this lncRNA.

### E. In-silico evaluation of functional significance of KLHDC7B-DT

Since functional annotations for the KLHDC7B-DT gene, such as Gene Ontology or KEGG pathways, are unavailable, we were unable to predict its associated categories. In pancreatic ductal adenocarcinoma models, KLHDC7B-DT has been shown to contribute to increased cell proliferation, invasion, and migration [11]. Mechanistically, KLHDC7B-DT binds to the IL-6_promoter, inducing IL-6 transactivation, STAT3 signaling, and the activation of M2 macrophage polarization in pancreatic ductal adenocarcinoma [11].

Beyond cancer, KLHDC7B-DT is overexpressed in psoriatic tissues, where it promotes keratinocyte proliferation, inflammation, and the secretion of IL-6 and IL-8 [12]. In this context, KLHDC7B-DT interacts with ILF2, enhancing STAT3 and JNK signalling [12].

Remarkably, HNCs are among the malignancies that activate the IL-6/STAT3 signaling axis [20, 21]. IL-6 has important effect in the processes of survival, proliferation, epithelial_to_mesenchymal transition (EMT), invasion, migration, NLRP3-inflammasome activation, and chemoresistance in HNC [20, 22–24]. In patients with HNC, increased IL-6 expression is an independent predictor of tumor recurrence, metastasis, and poor survival outcomes [20, 22-24]. As a cytokine, IL-6 interacts with its receptor to activate JAK/STAT3 signaling in HNCs [20, 24]. This activation of STAT3 is independent of EGFR, which means that anti-EGFR therapies may not always be effective for treating HNCs [20]. Furthermore, IL-6 is involved in the activation of additional signaling axis, including Wnt, PI3K, AKT, RAS, and MAPK [20].

Therefore, targeting IL-6 signaling either alone or in combination with conventional therapies may represent a promising therapeutic strategy for treating HNCs. Preclinical studies have shown encouraging results with anti-IL-6 therapies in HNCs [20, 24]. Although there are no current clinical trials specifically targeting IL-6 for HNC, a few studies have explored anti-IL-6 therapies, either alone or in combination, in clinical settings, with results yet to be obtained [24].

In addition to binding to RNA-binding proteins, lncRNAs can interact with miRNAs, thereby inhibiting miRNA-mediated degradation of target mRNAs [7, 8, 13, 25]. To explore potential miRNA partners for KLHDC7B-DT, we used the miRDB tool. Our analysis identified 44 miRNAs that may interact with KLHDC7B-DT (Supplementary Table 1), with the top predicted miRNAs shown in Fig. 5A. Notably, 13 of these miRNAs target IL-6 (Fig. 5B). This suggests that KLHDC7B-DT may modulate IL-6 in HNC through miRNAs, warranting validation in pre-clinical systems.

**Fig. 5.**
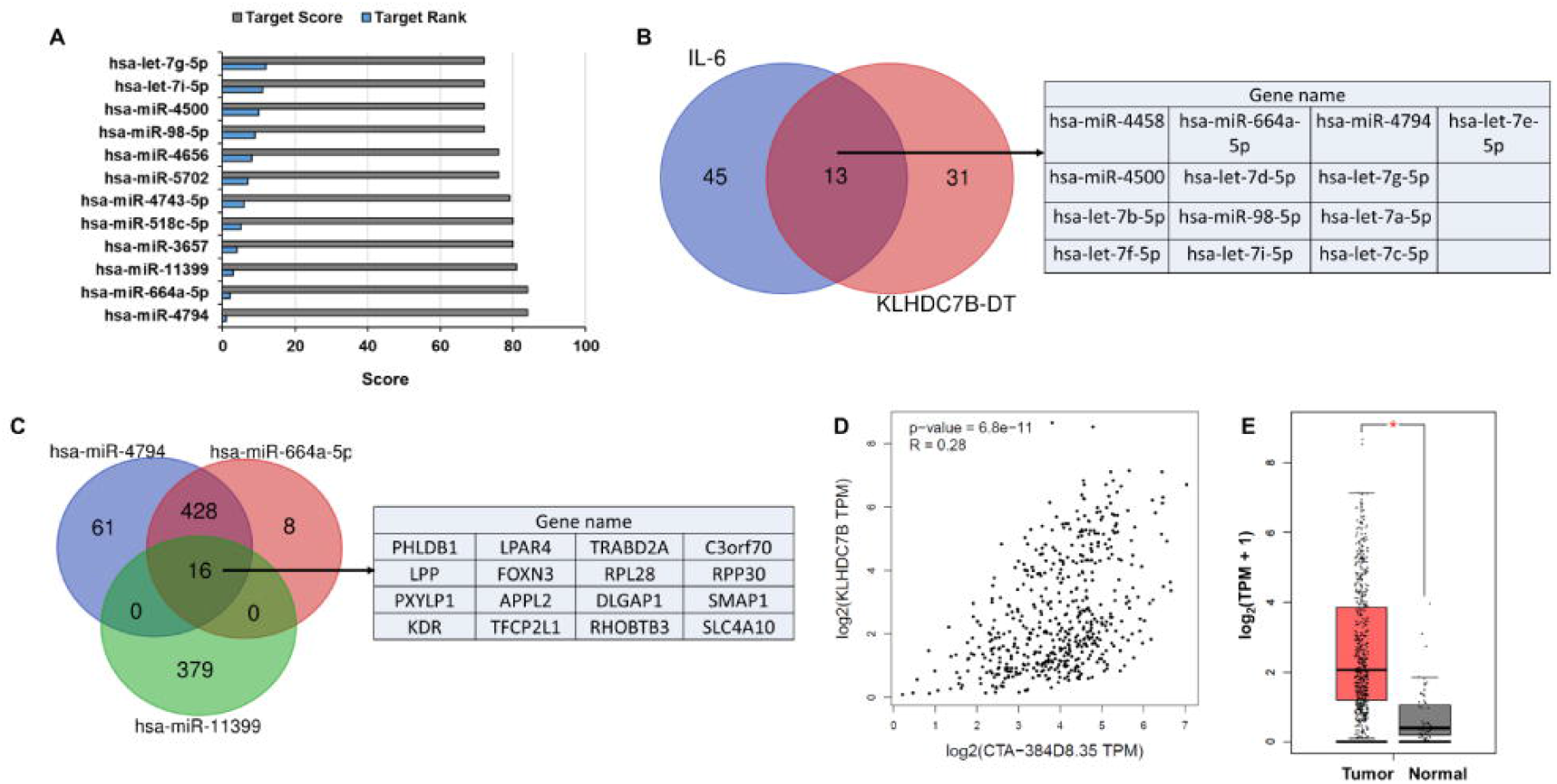
Functional Significance of KLHDC7B-DT. **A:** Top predicted miRNAs interacting with KLHDC7B-DT, identified through miRDB tool. **B:** Venn diagram illustrating the unique and common miRNAs interacting with both IL-6 and KLHDC7B-DT, with a table listing 13 common miRNAs as predicted by miRDB. **C:** Venn diagram showing unique and common target genes of the top three miRNAs interacting with KLHDC7B-DT, with a table displaying 16 common genes, as predicted by miRDB. **D:** Significant positive correlation between the expression of KLHDC7B-DT (CTA-384D8.35) and its adjacent protein-coding gene KLHDC7B in TCGA HNC data, analyzed using GEPIA. **E:** KLHDC7B is significantly upregulated in TCGA HNC samples (n=519) compared to normal samples (n=44), analyzed by GEPIA (*P < 0.01).

Based on target score and rank, has_miR-4794, has_miR-664a-5p, and has_miR-11399 are the top three miRNAs predicted to interact with KLHDC7B-DT. We identified 16 common mRNA targets of these three miRNAs (Fig. 5C) which are associated with many cancers related biological processes including: Angiogenesis, Cell Communication, Cell Cycle, Cell Migration, Developmental Process, Epithelial Cell Differentiation, GTPase Activation, Immune System Process, Increased Inflammatory Response, MAPK Signaling Pathway, Metabolic Process, PI3K_AKT Signaling, RAS_ Signaling, Regulation of Apoptotic Process, VEGF_ Signaling, etc. (Supplementary Table 2). This opens new directions for future mechanistic investigations of KLHDC7B-DT in HNC.

Additionally, cis-regulation of neighbouring protein-coding genes is another characteristic feature of lncRNAs [26]. The KLHDC7B gene is located in close proximity to the KLHDC7B-DT lncRNA but in opposite strand (Fig. 1A). Our analysis of TCGA HNC samples revealed a significantly positive correlation between the expression of KLHDC7B and KLHDC7B-DT (Fig. 5D). Furthermore, KLHDC7B is significantly upregulated in TCGA HNC samples (Fig. 5E), although its precise role in HNC remains unclear. KLHDC7B has been shown to be upregulated in breast cancer, where it regulates cell proliferation, interferon signaling, IFITs, STATs, and IL-29 signaling [27]. Additionally, in bladder cancer, KLHDC7B functions as a urine exosomal signal, promoting cell migration and proliferation while preventing apoptosis [28]. Thus, more mechanistic research is required to investigate the functional connection between KLHDC7B and KLHDC7B-DT in HNCs.

## IV. CONCLUSION

In conclusion, the lncRNA KLHDC7B-DT is upregulated in HNCs, and its elevated expression may be associated with cancer prognosis. This is the first study to report its expression and potential significance in HNC. We suggest a functional link between KLHDC7B-DT, IL-6, miRNAs, and KLHDC7B in this context. However, these findings are preliminary, and further mechanistic studies are needed to validate them in larger preclinical and clinical cohorts. This research paves the way for exploring KLHDC7B-DT’s role, not only as a diagnostic marker but also as a potential therapeutic target.

## Supporting information

Supplementary Table 1: Predicted miRNA interactions with KLHDC7B-DT Supplementary Table 2: Functional annotation of common target genes

## SUPPLEMENTARY MATERIALS

**Supplementary Table 1:** Predicted miRNA interactions with KLHDC7B-DT

**Supplementary Table 2:** Functional annotation of common target genes

## ACKNOWLEDGMENT

We express our gratitude to the Director, Dr. D. Y. Patil Biotechnology and Bioinformatics Institute, Pune for invaluable assistance and motivation.

## AUTHOR CONTRIBUTIONS

Conceptualization: S. Sur; Data Analysis and Interpretation: S. Basu, D. Davray, and S. Sur; Sample Procurement: S. Gupta and B. M. Rudagi; Histopathological Evaluation: S. Kheur; Writing – Original Draft Preparation: S. Sur. All authors have read and approved the final version of the manuscript.

## ETHICS APPROVAL

This study was performed in line with the approval granted by the Institutional Biosafety Committee (IBSC) [DYPBBI/1/2023 dated 7/1/2023] and Ethics Committee [DYPV/EC/910/23 dated 23/ January/ 2023], Dr. D.Y. Patil Biotechnology and Bioinformatics Institute, Dr. D. Y. Patil Vidyapeeth (DPU), Pune, India.

## DATA AVAILABILITY

All data generated in this study are included within the manuscript.

