## Supplementary Table 1: Predicted miRNA interactions with KLHDC7B-DT Supplementary Table 2: Functional annotation of common target genes for "Upregulation of a novel lncRNA KLHDC7B-DT in Head and Neck Cancers: Implications for Prognosis and Molecular Mechanisms"

**Supplementary Table 1: Predicted miRNA interactions with KLHDC7B-DT identified using the miRDB tool**

| miRNA Name | Target Rank | Target Score |
|---|---|---|
| hsa-miR-4794 | 1 | 84 |
| hsa-miR-664a-5p | 2 | 84 |
| hsa-miR-11399 | 3 | 81 |
| hsa-miR-3657 | 4 | 80 |
| hsa-miR-518c-5p | 5 | 80 |
| hsa-miR-4743-5p | 6 | 79 |
| hsa-miR-5702 | 7 | 76 |
| hsa-miR-4656 | 8 | 76 |
| hsa-miR-98-5p | 9 | 72 |
| hsa-miR-4500 | 10 | 72 |
| hsa-let-7i-5p | 11 | 72 |
| hsa-let-7g-5p | 12 | 72 |
| hsa-let-7f-5p | 13 | 72 |
| hsa-let-7e-5p | 14 | 72 |
| hsa-let-7c-5p | 15 | 72 |
| hsa-let-7b-5p | 16 | 72 |
| hsa-let-7a-5p | 17 | 72 |
| hsa-miR-4458 | 18 | 70 |
| hsa-let-7d-5p | 19 | 70 |
| hsa-miR-6857-3p | 20 | 68 |
| hsa-miR-12120 | 21 | 67 |
| hsa-miR-4478 | 22 | 65 |
| hsa-miR-4325 | 23 | 64 |
| hsa-miR-7160-5p | 24 | 62 |
| hsa-miR-4533 | 25 | 61 |
| hsa-miR-218-5p | 26 | 59 |
| hsa-miR-7113-5p | 27 | 58 |
| hsa-miR-4282 | 28 | 58 |
| hsa-miR-3169 | 29 | 57 |
| hsa-miR-3671 | 30 | 57 |
| hsa-miR-665 | 31 | 55 |
| hsa-miR-6837-5p | 32 | 54 |
| hsa-miR-4685-5p | 33 | 54 |
| hsa-miR-4745-3p | 34 | 53 |
| hsa-miR-9900 | 35 | 52 |
| hsa-miR-1538 | 36 | 52 |
| hsa-miR-4483 | 37 | 52 |
| hsa-miR-4747-3p | 38 | 51 |
| hsa-miR-593-3p | 39 | 51 |
| hsa-miR-1587 | 40 | 51 |
| hsa-miR-3160-3p | 41 | 51 |
| hsa-miR-4770 | 42 | 50 |
| hsa-miR-143-3p | 43 | 50 |
| hsa-miR-4471 | 44 | 50 |

**Supplementary Table 2: Functional annotation of common target genes of hsa-miR-4794, hsa-miR-664a-5p, and hsa-miR-11399 done by string-db.org**

| Gene name | identifier | category | term ID | term description |
|---|---|---|---|---|
| APPL2 | 9606.ENSP00000446917 | GO Process | GO:0001817 | Regulation of cytokine production |
| APPL2 | 9606.ENSP00000446917 | GO Process | GO:0001818 | Negative regulation of cytokine production |
| APPL2 | 9606.ENSP00000446917 | GO Process | GO:0002021 | Response to dietary excess |
| APPL2 | 9606.ENSP00000446917 | GO Process | GO:0002024 | Diet induced thermogenesis |
| APPL2 | 9606.ENSP00000446917 | GO Process | GO:0002682 | Regulation of immune system process |
| APPL2 | 9606.ENSP00000446917 | GO Process | GO:0002684 | Positive regulation of immune system process |
| APPL2 | 9606.ENSP00000446917 | GO Process | GO:0002697 | Regulation of immune effector process |
| APPL2 | 9606.ENSP00000446917 | GO Process | GO:0002699 | Positive regulation of immune effector process |
| APPL2 | 9606.ENSP00000446917 | GO Process | GO:0002831 | Regulation of response to biotic stimulus |
| APPL2 | 9606.ENSP00000446917 | GO Process | GO:0006606 | Protein import into nucleus |
| APPL2 | 9606.ENSP00000446917 | GO Process | GO:0006810 | Transport |
| APPL2 | 9606.ENSP00000446917 | GO Process | GO:0006886 | Intracellular protein transport |
| APPL2 | 9606.ENSP00000446917 | GO Process | GO:0006913 | Nucleocytoplasmic transport |
| APPL2 | 9606.ENSP00000446917 | GO Process | GO:0006950 | Response to stress |
| APPL2 | 9606.ENSP00000446917 | GO Process | GO:0007049 | Cell cycle |
| APPL2 | 9606.ENSP00000446917 | GO Process | GO:0007154 | Cell communication |
| APPL2 | 9606.ENSP00000446917 | GO Process | GO:0007165 | Signal transduction |
| APPL2 | 9606.ENSP00000446917 | GO Process | GO:0007166 | Cell surface receptor signaling pathway |
| APPL2 | 9606.ENSP00000446917 | GO Process | GO:0007167 | Enzyme-linked receptor protein signaling pathway |
| APPL2 | 9606.ENSP00000446917 | GO Process | GO:0007178 | Transmembrane receptor protein serine/threonine kinase signaling pathway |
| APPL2 | 9606.ENSP00000446917 | GO Process | GO:0007179 | Transforming growth factor beta receptor signaling pathway |
| APPL2 | 9606.ENSP00000446917 | GO Process | GO:0007346 | Regulation of mitotic cell cycle |
| APPL2 | 9606.ENSP00000446917 | GO Process | GO:0008104 | Protein localization |
| APPL2 | 9606.ENSP00000446917 | GO Process | GO:0008152 | Metabolic process |
| APPL2 | 9606.ENSP00000446917 | GO Process | GO:0008285 | Negative regulation of cell population proliferation |
| APPL2 | 9606.ENSP00000446917 | GO Process | GO:0009266 | Response to temperature stimulus |
| APPL2 | 9606.ENSP00000446917 | GO Process | GO:0009409 | Response to cold |
| APPL2 | 9606.ENSP00000446917 | GO Process | GO:0009605 | Response to external stimulus |
| APPL2 | 9606.ENSP00000446917 | GO Process | GO:0009628 | Response to abiotic stimulus |
| APPL2 | 9606.ENSP00000446917 | GO Process | GO:0009631 | Cold acclimation |

| | | | | |
|---|---|---|---|---|
| APPL2 | 9606.ENSP00000446917 | GO Process | GO:0009719 | Response to endogenous stimulus |
| APPL2 | 9606.ENSP00000446917 | GO Process | GO:0009725 | Response to hormone |
| APPL2 | 9606.ENSP00000446917 | GO Process | GO:0009755 | Hormone-mediated signaling pathway |
| APPL2 | 9606.ENSP00000446917 | GO Process | GO:0009892 | Negative regulation of metabolic process |
| APPL2 | 9606.ENSP00000446917 | GO Process | GO:0009893 | Positive regulation of metabolic process |
| APPL2 | 9606.ENSP00000446917 | GO Process | GO:0009966 | Regulation of signal transduction |
| APPL2 | 9606.ENSP00000446917 | GO Process | GO:0009967 | Positive regulation of signal transduction |
| APPL2 | 9606.ENSP00000446917 | GO Process | GO:0009987 | Cellular process |
| APPL2 | 9606.ENSP00000446917 | GO Process | GO:0009991 | Response to extracellular stimulus |
| APPL2 | 9606.ENSP00000446917 | GO Process | GO:0010033 | Response to organic substance |
| APPL2 | 9606.ENSP00000446917 | GO Process | GO:0010468 | Regulation of gene expression |
| APPL2 | 9606.ENSP00000446917 | GO Process | GO:0010564 | Regulation of cell cycle process |
| APPL2 | 9606.ENSP00000446917 | GO Process | GO:0010565 | Regulation of cellular ketone metabolic process |
| APPL2 | 9606.ENSP00000446917 | GO Process | GO:0010605 | Negative regulation of macromolecule metabolic process |
| APPL2 | 9606.ENSP00000446917 | GO Process | GO:0010629 | Negative regulation of gene expression |
| APPL2 | 9606.ENSP00000446917 | GO Process | GO:0010646 | Regulation of cell communication |
| APPL2 | 9606.ENSP00000446917 | GO Process | GO:0010647 | Positive regulation of cell communication |
| APPL2 | 9606.ENSP00000446917 | GO Process | GO:0010721 | Negative regulation of cell development |
| APPL2 | 9606.ENSP00000446917 | GO Process | GO:0010762 | Regulation of fibroblast migration |
| APPL2 | 9606.ENSP00000446917 | GO Process | GO:0010827 | Regulation of glucose transmembrane transport |
| APPL2 | 9606.ENSP00000446917 | GO Process | GO:0010829 | Negative regulation of glucose transmembrane transport |
| APPL2 | 9606.ENSP00000446917 | GO Process | GO:0015031 | Protein transport |
| APPL2 | 9606.ENSP00000446917 | GO Process | GO:0016043 | Cellular component organization |
| APPL2 | 9606.ENSP00000446917 | GO Process | GO:0019216 | Regulation of lipid metabolic process |
| APPL2 | 9606.ENSP00000446917 | GO Process | GO:0019217 | Regulation of fatty acid metabolic process |
| APPL2 | 9606.ENSP00000446917 | GO Process | GO:0019221 | Cytokine-mediated signaling pathway |
| APPL2 | 9606.ENSP00000446917 | GO Process | GO:0019222 | Regulation of metabolic process |
| APPL2 | 9606.ENSP00000446917 | GO Process | GO:0022607 | Cellular component assembly |
| APPL2 | 9606.ENSP00000446917 | GO Process | GO:0023051 | Regulation of signaling |
| APPL2 | 9606.ENSP00000446917 | GO Process | GO:0023052 | Signaling |
| APPL2 | 9606.ENSP00000446917 | GO Process | GO:0023056 | Positive regulation of signaling |
| APPL2 | 9606.ENSP00000446917 | GO Process | GO:0030100 | Regulation of endocytosis |
| APPL2 | 9606.ENSP00000446917 | GO Process | GO:0030334 | Regulation of cell migration |
| APPL2 | 9606.ENSP00000446917 | GO Process | GO:0031323 | Regulation of cellular metabolic process |
| APPL2 | 9606.ENSP00000446917 | GO Process | GO:0031324 | Negative regulation of cellular metabolic process |

| | | | | |
|---|---|---|---|---|
| APPL2 | 9606.ENSP00000446917 | GO Process | GO:0031347 | Regulation of defense response |
| APPL2 | 9606.ENSP00000446917 | GO Process | GO:0031667 | Response to nutrient levels |
| APPL2 | 9606.ENSP00000446917 | GO Process | GO:0032101 | Regulation of response to external stimulus |
| APPL2 | 9606.ENSP00000446917 | GO Process | GO:0032501 | Multicellular organismal process |
| APPL2 | 9606.ENSP00000446917 | GO Process | GO:0032870 | Cellular response to hormone stimulus |
| APPL2 | 9606.ENSP00000446917 | GO Process | GO:0032879 | Regulation of localization |
| APPL2 | 9606.ENSP00000446917 | GO Process | GO:0033036 | Macromolecule localization |
| APPL2 | 9606.ENSP00000446917 | GO Process | GO:0033211 | Adiponectin-activated signaling pathway |
| APPL2 | 9606.ENSP00000446917 | GO Process | GO:0033365 | Protein localization to organelle |
| APPL2 | 9606.ENSP00000446917 | GO Process | GO:0033500 | Carbohydrate homeostasis |
| APPL2 | 9606.ENSP00000446917 | GO Process | GO:0034097 | Response to cytokine |
| APPL2 | 9606.ENSP00000446917 | GO Process | GO:0034121 | Regulation of toll-like receptor signaling pathway |
| APPL2 | 9606.ENSP00000446917 | GO Process | GO:0034143 | Regulation of toll-like receptor 4 signaling pathway |
| APPL2 | 9606.ENSP00000446917 | GO Process | GO:0034504 | Protein localization to nucleus |
| APPL2 | 9606.ENSP00000446917 | GO Process | GO:0034762 | Regulation of transmembrane transport |
| APPL2 | 9606.ENSP00000446917 | GO Process | GO:0034763 | Negative regulation of transmembrane transport |
| APPL2 | 9606.ENSP00000446917 | GO Process | GO:0035728 | Response to hepatocyte growth factor |
| APPL2 | 9606.ENSP00000446917 | GO Process | GO:0035729 | Cellular response to hepatocyte growth factor stimulus |
| APPL2 | 9606.ENSP00000446917 | GO Process | GO:0040012 | Regulation of locomotion |
| APPL2 | 9606.ENSP00000446917 | GO Process | GO:0042127 | Regulation of cell population proliferation |
| APPL2 | 9606.ENSP00000446917 | GO Process | GO:0042221 | Response to chemical |
| APPL2 | 9606.ENSP00000446917 | GO Process | GO:0042592 | Homeostatic process |
| APPL2 | 9606.ENSP00000446917 | GO Process | GO:0042593 | Glucose homeostasis |
| APPL2 | 9606.ENSP00000446917 | GO Process | GO:0043933 | Protein-containing complex organization |
| APPL2 | 9606.ENSP00000446917 | GO Process | GO:0044085 | Cellular component biogenesis |
| APPL2 | 9606.ENSP00000446917 | GO Process | GO:0045088 | Regulation of innate immune response |
| APPL2 | 9606.ENSP00000446917 | GO Process | GO:0045184 | Establishment of protein localization |
| APPL2 | 9606.ENSP00000446917 | GO Process | GO:0045595 | Regulation of cell differentiation |
| APPL2 | 9606.ENSP00000446917 | GO Process | GO:0045596 | Negative regulation of cell differentiation |
| APPL2 | 9606.ENSP00000446917 | GO Process | GO:0045807 | Positive regulation of endocytosis |
| APPL2 | 9606.ENSP00000446917 | GO Process | GO:0045833 | Negative regulation of lipid metabolic process |
| APPL2 | 9606.ENSP00000446917 | GO Process | GO:0045922 | Negative regulation of fatty acid metabolic process |
| APPL2 | 9606.ENSP00000446917 | GO Process | GO:0046320 | Regulation of fatty acid oxidation |
| APPL2 | 9606.ENSP00000446917 | GO Process | GO:0046322 | Negative regulation of fatty acid oxidation |
| APPL2 | 9606.ENSP00000446917 | GO Process | GO:0046324 | Regulation of glucose import |

| | | | | |
|---|---|---|---|---|
| APPL2 | 9606.ENSP00000446917 | GO Process | GO:0046325 | Negative regulation of glucose import |
| APPL2 | 9606.ENSP00000446917 | GO Process | GO:0046907 | Intracellular transport |
| APPL2 | 9606.ENSP00000446917 | GO Process | GO:0048518 | Positive regulation of biological process |
| APPL2 | 9606.ENSP00000446917 | GO Process | GO:0048519 | Negative regulation of biological process |
| APPL2 | 9606.ENSP00000446917 | GO Process | GO:0048522 | Positive regulation of cellular process |
| APPL2 | 9606.ENSP00000446917 | GO Process | GO:0048523 | Negative regulation of cellular process |
| APPL2 | 9606.ENSP00000446917 | GO Process | GO:0048548 | Regulation of pinocytosis |
| APPL2 | 9606.ENSP00000446917 | GO Process | GO:0048549 | Positive regulation of pinocytosis |
| APPL2 | 9606.ENSP00000446917 | GO Process | GO:0048583 | Regulation of response to stimulus |
| APPL2 | 9606.ENSP00000446917 | GO Process | GO:0048584 | Positive regulation of response to stimulus |
| APPL2 | 9606.ENSP00000446917 | GO Process | GO:0048585 | Negative regulation of response to stimulus |
| APPL2 | 9606.ENSP00000446917 | GO Process | GO:0048878 | Chemical homeostasis |
| APPL2 | 9606.ENSP00000446917 | GO Process | GO:0050764 | Regulation of phagocytosis |
| APPL2 | 9606.ENSP00000446917 | GO Process | GO:0050766 | Positive regulation of phagocytosis |
| APPL2 | 9606.ENSP00000446917 | GO Process | GO:0050767 | Regulation of neurogenesis |
| APPL2 | 9606.ENSP00000446917 | GO Process | GO:0050768 | Negative regulation of neurogenesis |
| APPL2 | 9606.ENSP00000446917 | GO Process | GO:0050776 | Regulation of immune response |
| APPL2 | 9606.ENSP00000446917 | GO Process | GO:0050778 | Positive regulation of immune response |
| APPL2 | 9606.ENSP00000446917 | GO Process | GO:0050789 | Regulation of biological process |
| APPL2 | 9606.ENSP00000446917 | GO Process | GO:0050793 | Regulation of developmental process |
| APPL2 | 9606.ENSP00000446917 | GO Process | GO:0050794 | Regulation of cellular process |
| APPL2 | 9606.ENSP00000446917 | GO Process | GO:0050896 | Response to stimulus |
| APPL2 | 9606.ENSP00000446917 | GO Process | GO:0051049 | Regulation of transport |
| APPL2 | 9606.ENSP00000446917 | GO Process | GO:0051050 | Positive regulation of transport |
| APPL2 | 9606.ENSP00000446917 | GO Process | GO:0051051 | Negative regulation of transport |
| APPL2 | 9606.ENSP00000446917 | GO Process | GO:0051093 | Negative regulation of developmental process |
| APPL2 | 9606.ENSP00000446917 | GO Process | GO:0051128 | Regulation of cellular component organization |
| APPL2 | 9606.ENSP00000446917 | GO Process | GO:0051130 | Positive regulation of cellular component organization |
| APPL2 | 9606.ENSP00000446917 | GO Process | GO:0051169 | Nuclear transport |
| APPL2 | 9606.ENSP00000446917 | GO Process | GO:0051170 | Import into nucleus |
| APPL2 | 9606.ENSP00000446917 | GO Process | GO:0051179 | Localization |
| APPL2 | 9606.ENSP00000446917 | GO Process | GO:0051234 | Establishment of localization |
| APPL2 | 9606.ENSP00000446917 | GO Process | GO:0051239 | Regulation of multicellular organismal process |
| APPL2 | 9606.ENSP00000446917 | GO Process | GO:0051240 | Positive regulation of multicellular organismal process |
| APPL2 | 9606.ENSP00000446917 | GO Process | GO:0051241 | Negative regulation of multicellular organismal process |

| | | | | |
|---|---|---|---|---|
| APPL2 | 9606.ENSP00000446917 | GO Process | GO:0051259 | Protein complex oligomerization |
| APPL2 | 9606.ENSP00000446917 | GO Process | GO:0051260 | Protein homooligomerization |
| APPL2 | 9606.ENSP00000446917 | GO Process | GO:0051262 | Protein tetramerization |
| APPL2 | 9606.ENSP00000446917 | GO Process | GO:0051289 | Protein homotetramerization |
| APPL2 | 9606.ENSP00000446917 | GO Process | GO:0051641 | Cellular localization |
| APPL2 | 9606.ENSP00000446917 | GO Process | GO:0051649 | Establishment of localization in cell |
| APPL2 | 9606.ENSP00000446917 | GO Process | GO:0051716 | Cellular response to stimulus |
| APPL2 | 9606.ENSP00000446917 | GO Process | GO:0051726 | Regulation of cell cycle |
| APPL2 | 9606.ENSP00000446917 | GO Process | GO:0051960 | Regulation of nervous system development |
| APPL2 | 9606.ENSP00000446917 | GO Process | GO:0051961 | Negative regulation of nervous system development |
| APPL2 | 9606.ENSP00000446917 | GO Process | GO:0060099 | Regulation of phagocytosis, engulfment |
| APPL2 | 9606.ENSP00000446917 | GO Process | GO:0060100 | Positive regulation of phagocytosis, engulfment |
| APPL2 | 9606.ENSP00000446917 | GO Process | GO:0060255 | Regulation of macromolecule metabolic process |
| APPL2 | 9606.ENSP00000446917 | GO Process | GO:0060284 | Regulation of cell development |
| APPL2 | 9606.ENSP00000446917 | GO Process | GO:0060368 | Regulation of Fc receptor mediated stimulatory signaling pathway |
| APPL2 | 9606.ENSP00000446917 | GO Process | GO:0060369 | Positive regulation of Fc receptor mediated stimulatory signaling pathway |
| APPL2 | 9606.ENSP00000446917 | GO Process | GO:0060627 | Regulation of vesicle-mediated transport |
| APPL2 | 9606.ENSP00000446917 | GO Process | GO:0062012 | Regulation of small molecule metabolic process |
| APPL2 | 9606.ENSP00000446917 | GO Process | GO:0062014 | Negative regulation of small molecule metabolic process |
| APPL2 | 9606.ENSP00000446917 | GO Process | GO:0062207 | Regulation of pattern recognition receptor signaling pathway |
| APPL2 | 9606.ENSP00000446917 | GO Process | GO:0065003 | Protein-containing complex assembly |
| APPL2 | 9606.ENSP00000446917 | GO Process | GO:0065007 | Biological regulation |
| APPL2 | 9606.ENSP00000446917 | GO Process | GO:0065008 | Regulation of biological quality |
| APPL2 | 9606.ENSP00000446917 | GO Process | GO:0070727 | Cellular macromolecule localization |
| APPL2 | 9606.ENSP00000446917 | GO Process | GO:0070848 | Response to growth factor |
| APPL2 | 9606.ENSP00000446917 | GO Process | GO:0070887 | Cellular response to chemical stimulus |
| APPL2 | 9606.ENSP00000446917 | GO Process | GO:0071310 | Cellular response to organic substance |
| APPL2 | 9606.ENSP00000446917 | GO Process | GO:0071345 | Cellular response to cytokine stimulus |
| APPL2 | 9606.ENSP00000446917 | GO Process | GO:0071363 | Cellular response to growth factor stimulus |
| APPL2 | 9606.ENSP00000446917 | GO Process | GO:0071495 | Cellular response to endogenous stimulus |
| APPL2 | 9606.ENSP00000446917 | GO Process | GO:0071559 | Response to transforming growth factor beta |
| APPL2 | 9606.ENSP00000446917 | GO Process | GO:0071560 | Cellular response to transforming growth factor beta stimulus |
| APPL2 | 9606.ENSP00000446917 | GO Process | GO:0071702 | Organic substance transport |
| APPL2 | 9606.ENSP00000446917 | GO Process | GO:0071705 | Nitrogen compound transport |
| APPL2 | 9606.ENSP00000446917 | GO Process | GO:0071840 | Cellular component organization or biogenesis |

| | | | | |
|---|---|---|---|---|
| APPL2 | 9606.ENSP00000446917 | GO Process | GO:0072594 | Establishment of protein localization to organelle |
| APPL2 | 9606.ENSP00000446917 | GO Process | GO:0080090 | Regulation of primary metabolic process |
| APPL2 | 9606.ENSP00000446917 | GO Process | GO:0080134 | Regulation of response to stress |
| APPL2 | 9606.ENSP00000446917 | GO Process | GO:0120161 | Regulation of cold-induced thermogenesis |
| APPL2 | 9606.ENSP00000446917 | GO Process | GO:0120162 | Positive regulation of cold-induced thermogenesis |
| APPL2 | 9606.ENSP00000446917 | GO Process | GO:1900015 | Regulation of cytokine production involved in inflammatory response |
| APPL2 | 9606.ENSP00000446917 | GO Process | GO:1900016 | Negative regulation of cytokine production involved in inflammatory response |
| APPL2 | 9606.ENSP00000446917 | GO Process | GO:1900076 | Regulation of cellular response to insulin stimulus |
| APPL2 | 9606.ENSP00000446917 | GO Process | GO:1900077 | Negative regulation of cellular response to insulin stimulus |
| APPL2 | 9606.ENSP00000446917 | GO Process | GO:1901987 | Regulation of cell cycle phase transition |
| APPL2 | 9606.ENSP00000446917 | GO Process | GO:1901990 | Regulation of mitotic cell cycle phase transition |
| APPL2 | 9606.ENSP00000446917 | GO Process | GO:1902806 | Regulation of cell cycle G1/S phase transition |
| APPL2 | 9606.ENSP00000446917 | GO Process | GO:1905153 | Regulation of membrane invagination |
| APPL2 | 9606.ENSP00000446917 | GO Process | GO:1905155 | Positive regulation of membrane invagination |
| APPL2 | 9606.ENSP00000446917 | GO Process | GO:1905301 | Regulation of macropinocytosis |
| APPL2 | 9606.ENSP00000446917 | GO Process | GO:1905303 | Positive regulation of macropinocytosis |
| APPL2 | 9606.ENSP00000446917 | GO Process | GO:1905449 | Regulation of Fc-gamma receptor signaling pathway involved in phagocytosis |
| APPL2 | 9606.ENSP00000446917 | GO Process | GO:1905451 | Positive regulation of Fc-gamma receptor signaling pathway involved in phagocytosis |
| APPL2 | 9606.ENSP00000446917 | GO Process | GO:1990845 | Adaptive thermogenesis |
| APPL2 | 9606.ENSP00000446917 | GO Process | GO:2000026 | Regulation of multicellular organismal development |
| APPL2 | 9606.ENSP00000446917 | GO Process | GO:2000045 | Regulation of G1/S transition of mitotic cell cycle |
| APPL2 | 9606.ENSP00000446917 | GO Process | GO:2000145 | Regulation of cell motility |
| APPL2 | 9606.ENSP00000446917 | GO Process | GO:2000177 | Regulation of neural precursor cell proliferation |
| APPL2 | 9606.ENSP00000446917 | GO Process | GO:2000178 | Negative regulation of neural precursor cell proliferation |
| APPL2 | 9606.ENSP00000446917 | GO Function | GO:0001786 | Phosphatidylserine binding |
| APPL2 | 9606.ENSP00000446917 | GO Function | GO:0005488 | Binding |
| APPL2 | 9606.ENSP00000446917 | GO Function | GO:0005515 | Protein binding |
| APPL2 | 9606.ENSP00000446917 | GO Function | GO:0005543 | Phospholipid binding |
| APPL2 | 9606.ENSP00000446917 | GO Function | GO:0008289 | Lipid binding |
| APPL2 | 9606.ENSP00000446917 | GO Function | GO:0035091 | Phosphatidylinositol binding |
| APPL2 | 9606.ENSP00000446917 | GO Function | GO:0042802 | Identical protein binding |
| APPL2 | 9606.ENSP00000446917 | GO Function | GO:0042803 | Protein homodimerization activity |

| | | | | |
|---|---|---|---|---|
| APPL2 | 9606.ENSP00000446917 | GO Function | GO:0044877 | Protein-containing complex binding |
| APPL2 | 9606.ENSP00000446917 | GO Function | GO:0046983 | Protein dimerization activity |
| APPL2 | 9606.ENSP00000446917 | GO Function | GO:0072341 | Modified amino acid binding |
| APPL2 | 9606.ENSP00000446917 | GO Component | GO:0001726 | Ruffle |
| APPL2 | 9606.ENSP00000446917 | GO Component | GO:0005576 | Extracellular region |
| APPL2 | 9606.ENSP00000446917 | GO Component | GO:0005615 | Extracellular space |
| APPL2 | 9606.ENSP00000446917 | GO Component | GO:0005622 | Intracellular anatomical structure |
| APPL2 | 9606.ENSP00000446917 | GO Component | GO:0005634 | Nucleus |
| APPL2 | 9606.ENSP00000446917 | GO Component | GO:0005737 | Cytoplasm |
| APPL2 | 9606.ENSP00000446917 | GO Component | GO:0005768 | Endosome |
| APPL2 | 9606.ENSP00000446917 | GO Component | GO:0005769 | Early endosome |
| APPL2 | 9606.ENSP00000446917 | GO Component | GO:0005886 | Plasma membrane |
| APPL2 | 9606.ENSP00000446917 | GO Component | GO:0010008 | Endosome membrane |
| APPL2 | 9606.ENSP00000446917 | GO Component | GO:0012505 | Endomembrane system |
| APPL2 | 9606.ENSP00000446917 | GO Component | GO:0012506 | Vesicle membrane |
| APPL2 | 9606.ENSP00000446917 | GO Component | GO:0016020 | Membrane |
| APPL2 | 9606.ENSP00000446917 | GO Component | GO:0030139 | Endocytic vesicle |
| APPL2 | 9606.ENSP00000446917 | GO Component | GO:0030659 | Cytoplasmic vesicle membrane |
| APPL2 | 9606.ENSP00000446917 | GO Component | GO:0030666 | Endocytic vesicle membrane |
| APPL2 | 9606.ENSP00000446917 | GO Component | GO:0030670 | Phagocytic vesicle membrane |
| APPL2 | 9606.ENSP00000446917 | GO Component | GO:0031090 | Organelle membrane |
| APPL2 | 9606.ENSP00000446917 | GO Component | GO:0031252 | Cell leading edge |
| APPL2 | 9606.ENSP00000446917 | GO Component | GO:0031253 | Cell projection membrane |
| APPL2 | 9606.ENSP00000446917 | GO Component | GO:0031256 | Leading edge membrane |
| APPL2 | 9606.ENSP00000446917 | GO Component | GO:0031410 | Cytoplasmic vesicle |
| APPL2 | 9606.ENSP00000446917 | GO Component | GO:0031901 | Early endosome membrane |
| APPL2 | 9606.ENSP00000446917 | GO Component | GO:0031982 | Vesicle |
| APPL2 | 9606.ENSP00000446917 | GO Component | GO:0032009 | Early phagosome |
| APPL2 | 9606.ENSP00000446917 | GO Component | GO:0032587 | Ruffle membrane |
| APPL2 | 9606.ENSP00000446917 | GO Component | GO:0036186 | Early phagosome membrane |
| APPL2 | 9606.ENSP00000446917 | GO Component | GO:0042995 | Cell projection |
| APPL2 | 9606.ENSP00000446917 | GO Component | GO:0043226 | Organelle |
| APPL2 | 9606.ENSP00000446917 | GO Component | GO:0043227 | Membrane-bounded organelle |
| APPL2 | 9606.ENSP00000446917 | GO Component | GO:0043229 | Intracellular organelle |
| APPL2 | 9606.ENSP00000446917 | GO Component | GO:0043230 | Extracellular organelle |

| | | | | |
|---|---|---|---|---|
| APPL2 | 9606.ENSP00000446917 | GO Component | GO:0043231 | Intracellular membrane-bounded organelle |
| APPL2 | 9606.ENSP00000446917 | GO Component | GO:0044352 | Pinosome |
| APPL2 | 9606.ENSP00000446917 | GO Component | GO:0044354 | Macropinosome |
| APPL2 | 9606.ENSP00000446917 | GO Component | GO:0045335 | Phagocytic vesicle |
| APPL2 | 9606.ENSP00000446917 | GO Component | GO:0065010 | Extracellular membrane-bounded organelle |
| APPL2 | 9606.ENSP00000446917 | GO Component | GO:0070062 | Extracellular exosome |
| APPL2 | 9606.ENSP00000446917 | GO Component | GO:0071944 | Cell periphery |
| APPL2 | 9606.ENSP00000446917 | GO Component | GO:0097708 | Intracellular vesicle |
| APPL2 | 9606.ENSP00000446917 | GO Component | GO:0098588 | Bounding membrane of organelle |
| APPL2 | 9606.ENSP00000446917 | GO Component | GO:0098590 | Plasma membrane region |
| APPL2 | 9606.ENSP00000446917 | GO Component | GO:0110165 | Cellular anatomical entity |
| APPL2 | 9606.ENSP00000446917 | GO Component | GO:0120025 | Plasma membrane bounded cell projection |
| APPL2 | 9606.ENSP00000446917 | GO Component | GO:1903561 | Extracellular vesicle |
| APPL2 | 9606.ENSP00000446917 | STRING clusters | CL:25990 | Centrosome maturation, and Centriole |
| APPL2 | 9606.ENSP00000446917 | STRING clusters | CL:26162 | Mixed, incl. Rab-like protein 2, and Immunodeficiency 46 |
| APPL2 | 9606.ENSP00000446917 | TISSUES | BTO:0000000 | Tissues, cell types and enzyme sources |
| APPL2 | 9606.ENSP00000446917 | TISSUES | BTO:0000042 | Animal |
| APPL2 | 9606.ENSP00000446917 | COMPARTMENTS | GOCC:0001726 | Ruffle |
| APPL2 | 9606.ENSP00000446917 | COMPARTMENTS | GOCC:0005622 | Intracellular |
| APPL2 | 9606.ENSP00000446917 | COMPARTMENTS | GOCC:0005634 | Nucleus |
| APPL2 | 9606.ENSP00000446917 | COMPARTMENTS | GOCC:0005737 | Cytoplasm |
| APPL2 | 9606.ENSP00000446917 | COMPARTMENTS | GOCC:0005768 | Endosome |
| APPL2 | 9606.ENSP00000446917 | COMPARTMENTS | GOCC:0005886 | Plasma membrane |
| APPL2 | 9606.ENSP00000446917 | COMPARTMENTS | GOCC:0010008 | Endosome membrane |
| APPL2 | 9606.ENSP00000446917 | COMPARTMENTS | GOCC:0012505 | Endomembrane system |
| APPL2 | 9606.ENSP00000446917 | COMPARTMENTS | GOCC:0012506 | Vesicle membrane |
| APPL2 | 9606.ENSP00000446917 | COMPARTMENTS | GOCC:0016020 | Membrane |
| APPL2 | 9606.ENSP00000446917 | COMPARTMENTS | GOCC:0030139 | Endocytic vesicle |
| APPL2 | 9606.ENSP00000446917 | COMPARTMENTS | GOCC:0030659 | Cytoplasmic vesicle membrane |
| APPL2 | 9606.ENSP00000446917 | COMPARTMENTS | GOCC:0030666 | Endocytic vesicle membrane |
| APPL2 | 9606.ENSP00000446917 | COMPARTMENTS | GOCC:0030670 | Phagocytic vesicle membrane |
| APPL2 | 9606.ENSP00000446917 | COMPARTMENTS | GOCC:0031090 | Organelle membrane |
| APPL2 | 9606.ENSP00000446917 | COMPARTMENTS | GOCC:0031252 | Cell leading edge |
| APPL2 | 9606.ENSP00000446917 | COMPARTMENTS | GOCC:0031253 | Cell projection membrane |
| APPL2 | 9606.ENSP00000446917 | COMPARTMENTS | GOCC:0031256 | Leading edge membrane |

| | | | | |
|---|---|---|---|---|
| APPL2 | 9606.ENSP00000446917 | COMPARTMENTS | GOCC:0031410 | Cytoplasmic vesicle |
| APPL2 | 9606.ENSP00000446917 | COMPARTMENTS | GOCC:0031982 | Vesicle |
| APPL2 | 9606.ENSP00000446917 | COMPARTMENTS | GOCC:0032009 | Early phagosome |
| APPL2 | 9606.ENSP00000446917 | COMPARTMENTS | GOCC:0032587 | Ruffle membrane |
| APPL2 | 9606.ENSP00000446917 | COMPARTMENTS | GOCC:0036186 | Early phagosome membrane |
| APPL2 | 9606.ENSP00000446917 | COMPARTMENTS | GOCC:0042995 | Cell projection |
| APPL2 | 9606.ENSP00000446917 | COMPARTMENTS | GOCC:0043226 | Organelle |
| APPL2 | 9606.ENSP00000446917 | COMPARTMENTS | GOCC:0043227 | Membrane-bounded organelle |
| APPL2 | 9606.ENSP00000446917 | COMPARTMENTS | GOCC:0043229 | Intracellular organelle |
| APPL2 | 9606.ENSP00000446917 | COMPARTMENTS | GOCC:0043231 | Intracellular membrane-bounded organelle |
| APPL2 | 9606.ENSP00000446917 | COMPARTMENTS | GOCC:0044352 | Pinosome |
| APPL2 | 9606.ENSP00000446917 | COMPARTMENTS | GOCC:0044354 | Macropinosome |
| APPL2 | 9606.ENSP00000446917 | COMPARTMENTS | GOCC:0045335 | Phagocytic vesicle |
| APPL2 | 9606.ENSP00000446917 | COMPARTMENTS | GOCC:0071944 | Cell periphery |
| APPL2 | 9606.ENSP00000446917 | COMPARTMENTS | GOCC:0097708 | Intracellular vesicle |
| APPL2 | 9606.ENSP00000446917 | COMPARTMENTS | GOCC:0098588 | Bounding membrane of organelle |
| APPL2 | 9606.ENSP00000446917 | COMPARTMENTS | GOCC:0098590 | Plasma membrane region |
| APPL2 | 9606.ENSP00000446917 | COMPARTMENTS | GOCC:0110165 | Cellular anatomical entity |
| APPL2 | 9606.ENSP00000446917 | COMPARTMENTS | GOCC:0120025 | Plasma membrane bounded cell projection |
| APPL2 | 9606.ENSP00000446917 | Monarch | EFO:0004784 | Self reported educational attainment |
| APPL2 | 9606.ENSP00000446917 | Monarch | EFO:0008376 | Mosquito bite measurement |
| APPL2 | 9606.ENSP00000446917 | Monarch | EFO:0008378 | Mosquito bite reaction size measurement |
| APPL2 | 9606.ENSP00000446917 | Monarch | EFO:0011015 | Educational attainment |
| APPL2 | 9606.ENSP00000446917 | UniProt Keywords | KW-0025 | Alternative splicing |
| APPL2 | 9606.ENSP00000446917 | UniProt Keywords | KW-0131 | Cell cycle |
| APPL2 | 9606.ENSP00000446917 | UniProt Keywords | KW-0160 | Chromosomal rearrangement |
| APPL2 | 9606.ENSP00000446917 | UniProt Keywords | KW-0472 | Membrane |
| APPL2 | 9606.ENSP00000446917 | UniProt Keywords | KW-0539 | Nucleus |
| APPL2 | 9606.ENSP00000446917 | UniProt Keywords | KW-0963 | Cytoplasm |
| APPL2 | 9606.ENSP00000446917 | UniProt Keywords | KW-0966 | Cell projection |
| APPL2 | 9606.ENSP00000446917 | UniProt Keywords | KW-0967 | Endosome |
| APPL2 | 9606.ENSP00000446917 | UniProt Keywords | KW-0968 | Cytoplasmic vesicle |
| APPL2 | 9606.ENSP00000446917 | UniProt Keywords | KW-1003 | Cell membrane |
| APPL2 | 9606.ENSP00000446917 | Pfam | PF00640 | Phosphotyrosine interaction domain (PTB/PID) |
| APPL2 | 9606.ENSP00000446917 | Pfam | PF16746 | BAR domain of APPL family |

| | | | | |
|---|---|---|---|---|
| APPL2 | 9606.ENSP00000446917 | InterPro | IPR001849 | Pleckstrin homology domain |
| APPL2 | 9606.ENSP00000446917 | InterPro | IPR004148 | BAR domain |
| APPL2 | 9606.ENSP00000446917 | InterPro | IPR006020 | PTB/PI domain |
| APPL2 | 9606.ENSP00000446917 | InterPro | IPR011993 | PH-like domain superfamily |
| APPL2 | 9606.ENSP00000446917 | InterPro | IPR027267 | AH/BAR domain superfamily |
| APPL2 | 9606.ENSP00000446917 | InterPro | IPR047181 | DCC-interacting protein 13-alpha/beta |
| APPL2 | 9606.ENSP00000446917 | InterPro | IPR047236 | DCC-interacting protein 13-alpha/beta, PH domain |
| APPL2 | 9606.ENSP00000446917 | InterPro | IPR047237 | DCC-interacting protein 13-alpha/beta, PTB domain |
| APPL2 | 9606.ENSP00000446917 | InterPro | IPR047239 | DCC-interacting protein 13-beta, BAR domain |
| APPL2 | 9606.ENSP00000446917 | SMART | SM00233 | Pleckstrin homology domain. |
| C3orf70 | 9606.ENSP00000334974 | GO Process | GO:0007275 | Multicellular organism development |
| C3orf70 | 9606.ENSP00000334974 | GO Process | GO:0007399 | Nervous system development |
| C3orf70 | 9606.ENSP00000334974 | GO Process | GO:0007610 | Behavior |
| C3orf70 | 9606.ENSP00000334974 | GO Process | GO:0007622 | Rhythmic behavior |
| C3orf70 | 9606.ENSP00000334974 | GO Process | GO:0007623 | Circadian rhythm |
| C3orf70 | 9606.ENSP00000334974 | GO Process | GO:0032501 | Multicellular organismal process |
| C3orf70 | 9606.ENSP00000334974 | GO Process | GO:0032502 | Developmental process |
| C3orf70 | 9606.ENSP00000334974 | GO Process | GO:0048511 | Rhythmic process |
| C3orf70 | 9606.ENSP00000334974 | GO Process | GO:0048512 | Circadian behavior |
| C3orf70 | 9606.ENSP00000334974 | GO Process | GO:0048731 | System development |
| C3orf70 | 9606.ENSP00000334974 | GO Process | GO:0048856 | Anatomical structure development |
| C3orf70 | 9606.ENSP00000334974 | STRING clusters | CL:32614 | Mostly uncharacterized, incl. Interferon-induced transmembrane protein, and Potassium channel tetramerisation-type BTB domain |
| C3orf70 | 9606.ENSP00000334974 | STRING clusters | CL:32712 | Mixed, incl. Potassium channel tetramerisation-type BTB domain, and BTBD10/KCTD20, BTB/POZ domain |
| C3orf70 | 9606.ENSP00000334974 | STRING clusters | CL:32754 | Mixed, incl. Potassium channel tetramerisation-type BTB domain, and BTBD10/KCTD20, BTB/POZ domain |
| C3orf70 | 9606.ENSP00000334974 | STRING clusters | CL:32784 | Mixed, incl. SCAN domain, and KN motif |
| C3orf70 | 9606.ENSP00000334974 | Monarch | EFO:0004784 | Self reported educational attainment |
| C3orf70 | 9606.ENSP00000334974 | Monarch | EFO:0011015 | Educational attainment |
| C3orf70 | 9606.ENSP00000334974 | UniProt Keywords | KW-0524 | Neurogenesis |
| C3orf70 | 9606.ENSP00000334974 | Pfam | PF15823 | UPF0524 of C3orf70 |
| C3orf70 | 9606.ENSP00000334974 | InterPro | IPR029670 | UPF0524 family |
| DLGAP1 | 9606.ENSP00000316377 | GO Process | GO:0007154 | Cell communication |
| DLGAP1 | 9606.ENSP00000316377 | GO Process | GO:0007267 | Cell-cell signaling |

| | | | | |
|---|---|---|---|---|
| DLGAP1 | 9606.ENSP00000316377 | GO Process | GO:0007268 | Chemical synaptic transmission |
| DLGAP1 | 9606.ENSP00000316377 | GO Process | GO:0009966 | Regulation of signal transduction |
| DLGAP1 | 9606.ENSP00000316377 | GO Process | GO:0009987 | Cellular process |
| DLGAP1 | 9606.ENSP00000316377 | GO Process | GO:0010469 | Regulation of signaling receptor activity |
| DLGAP1 | 9606.ENSP00000316377 | GO Process | GO:0010646 | Regulation of cell communication |
| DLGAP1 | 9606.ENSP00000316377 | GO Process | GO:0023051 | Regulation of signaling |
| DLGAP1 | 9606.ENSP00000316377 | GO Process | GO:0023052 | Signaling |
| DLGAP1 | 9606.ENSP00000316377 | GO Process | GO:0031644 | Regulation of nervous system process |
| DLGAP1 | 9606.ENSP00000316377 | GO Process | GO:0044057 | Regulation of system process |
| DLGAP1 | 9606.ENSP00000316377 | GO Process | GO:0048583 | Regulation of response to stimulus |
| DLGAP1 | 9606.ENSP00000316377 | GO Process | GO:0050789 | Regulation of biological process |
| DLGAP1 | 9606.ENSP00000316377 | GO Process | GO:0050794 | Regulation of cellular process |
| DLGAP1 | 9606.ENSP00000316377 | GO Process | GO:0050804 | Modulation of chemical synaptic transmission |
| DLGAP1 | 9606.ENSP00000316377 | GO Process | GO:0051239 | Regulation of multicellular organismal process |
| DLGAP1 | 9606.ENSP00000316377 | GO Process | GO:0065007 | Biological regulation |
| DLGAP1 | 9606.ENSP00000316377 | GO Process | GO:0065009 | Regulation of molecular function |
| DLGAP1 | 9606.ENSP00000316377 | GO Process | GO:0098916 | Anterograde trans-synaptic signaling |
| DLGAP1 | 9606.ENSP00000316377 | GO Process | GO:0098962 | Regulation of postsynaptic neurotransmitter receptor activity |
| DLGAP1 | 9606.ENSP00000316377 | GO Process | GO:0099177 | Regulation of trans-synaptic signaling |
| DLGAP1 | 9606.ENSP00000316377 | GO Process | GO:0099536 | Synaptic signaling |
| DLGAP1 | 9606.ENSP00000316377 | GO Process | GO:0099537 | Trans-synaptic signaling |
| DLGAP1 | 9606.ENSP00000316377 | GO Process | GO:0099601 | Regulation of neurotransmitter receptor activity |
| DLGAP1 | 9606.ENSP00000316377 | GO Function | GO:0005488 | Binding |
| DLGAP1 | 9606.ENSP00000316377 | GO Function | GO:0044877 | Protein-containing complex binding |
| DLGAP1 | 9606.ENSP00000316377 | GO Function | GO:0060090 | Molecular adaptor activity |
| DLGAP1 | 9606.ENSP00000316377 | GO Component | GO:0005886 | Plasma membrane |
| DLGAP1 | 9606.ENSP00000316377 | GO Component | GO:0014069 | Postsynaptic density |
| DLGAP1 | 9606.ENSP00000316377 | GO Component | GO:0016020 | Membrane |
| DLGAP1 | 9606.ENSP00000316377 | GO Component | GO:0030054 | Cell junction |
| DLGAP1 | 9606.ENSP00000316377 | GO Component | GO:0032279 | Asymmetric synapse |
| DLGAP1 | 9606.ENSP00000316377 | GO Component | GO:0043226 | Organelle |
| DLGAP1 | 9606.ENSP00000316377 | GO Component | GO:0045202 | Synapse |
| DLGAP1 | 9606.ENSP00000316377 | GO Component | GO:0070161 | Anchoring junction |
| DLGAP1 | 9606.ENSP00000316377 | GO Component | GO:0071944 | Cell periphery |
| DLGAP1 | 9606.ENSP00000316377 | GO Component | GO:0098794 | Postsynapse |

| | | | | |
|---|---|---|---|---|
| DLGAP1 | 9606.ENSP00000316377 | GO Component | GO:0098978 | Glutamatergic synapse |
| DLGAP1 | 9606.ENSP00000316377 | GO Component | GO:0098984 | Neuron to neuron synapse |
| DLGAP1 | 9606.ENSP00000316377 | GO Component | GO:0099572 | Postsynaptic specialization |
| DLGAP1 | 9606.ENSP00000316377 | GO Component | GO:0110165 | Cellular anatomical entity |
| DLGAP1 | 9606.ENSP00000316377 | STRING clusters | CL:22977 | Postsynaptic cell membrane, and Protein-protein interactions at synapses |
| DLGAP1 | 9606.ENSP00000316377 | STRING clusters | CL:22978 | Transmitter-gated channel activity, and Neurexins and neuroligins |
| DLGAP1 | 9606.ENSP00000316377 | STRING clusters | CL:22979 | Neurotransmitter receptor complex, and Neurexins and neuroligins |
| DLGAP1 | 9606.ENSP00000316377 | STRING clusters | CL:22980 | Neurexins and neuroligins, and Regulation of presynapse assembly |
| DLGAP1 | 9606.ENSP00000316377 | STRING clusters | CL:22982 | Neurexins and neuroligins |
| DLGAP1 | 9606.ENSP00000316377 | STRING clusters | CL:22984 | Neurexins and neuroligins |
| DLGAP1 | 9606.ENSP00000316377 | STRING clusters | CL:22986 | Vocalization behavior, and Homer family |
| DLGAP1 | 9606.ENSP00000316377 | STRING clusters | CL:22988 | Vocalization behavior |
| DLGAP1 | 9606.ENSP00000316377 | STRING clusters | CL:22991 | Phelan-McDermid syndrome, and Positive regulation of AMPA glutamate receptor clustering |
| DLGAP1 | 9606.ENSP00000316377 | KEGG | hsa04724 | Glutamatergic synapse |
| DLGAP1 | 9606.ENSP00000316377 | Reactome | HSA-112316 | Neuronal System |
| DLGAP1 | 9606.ENSP00000316377 | Reactome | HSA-6794361 | Neurexins and neuroligins |
| DLGAP1 | 9606.ENSP00000316377 | Reactome | HSA-6794362 | Protein-protein interactions at synapses |
| DLGAP1 | 9606.ENSP00000316377 | WikiPathways | WP4875 | Disruption of postsynaptic signaling by CNV |
| DLGAP1 | 9606.ENSP00000316377 | TISSUES | BTO:0000000 | Tissues, cell types and enzyme sources |
| DLGAP1 | 9606.ENSP00000316377 | TISSUES | BTO:0000042 | Animal |
| DLGAP1 | 9606.ENSP00000316377 | TISSUES | BTO:0000081 | Reproductive system |
| DLGAP1 | 9606.ENSP00000316377 | TISSUES | BTO:0000083 | Female reproductive system |
| DLGAP1 | 9606.ENSP00000316377 | TISSUES | BTO:0000142 | Brain |
| DLGAP1 | 9606.ENSP00000316377 | TISSUES | BTO:0000227 | Central nervous system |
| DLGAP1 | 9606.ENSP00000316377 | TISSUES | BTO:0000282 | Head |
| DLGAP1 | 9606.ENSP00000316377 | TISSUES | BTO:0000342 | Diencephalon |
| DLGAP1 | 9606.ENSP00000316377 | TISSUES | BTO:0000478 | Forebrain |
| DLGAP1 | 9606.ENSP00000316377 | TISSUES | BTO:0001365 | Thalamus |
| DLGAP1 | 9606.ENSP00000316377 | TISSUES | BTO:0001484 | Nervous system |
| DLGAP1 | 9606.ENSP00000316377 | TISSUES | BTO:0001489 | Whole body |
| DLGAP1 | 9606.ENSP00000316377 | TISSUES | BTO:0003091 | Urogenital system |
| DLGAP1 | 9606.ENSP00000316377 | COMPARTMENTS | GOCC:0014069 | Postsynaptic density |
| DLGAP1 | 9606.ENSP00000316377 | COMPARTMENTS | GOCC:0030054 | Cell junction |
| DLGAP1 | 9606.ENSP00000316377 | COMPARTMENTS | GOCC:0032279 | Asymmetric synapse |

| | | | | |
|---|---|---|---|---|
| DLGAP1 | 9606.ENSP00000316377 | COMPARTMENTS | GOCC:0043226 | Organelle |
| DLGAP1 | 9606.ENSP00000316377 | COMPARTMENTS | GOCC:0045202 | Synapse |
| DLGAP1 | 9606.ENSP00000316377 | COMPARTMENTS | GOCC:0098794 | Postsynapse |
| DLGAP1 | 9606.ENSP00000316377 | COMPARTMENTS | GOCC:0098978 | Glutamatergic synapse |
| DLGAP1 | 9606.ENSP00000316377 | COMPARTMENTS | GOCC:0098984 | Neuron to neuron synapse |
| DLGAP1 | 9606.ENSP00000316377 | COMPARTMENTS | GOCC:0099572 | Postsynaptic specialization |
| DLGAP1 | 9606.ENSP00000316377 | COMPARTMENTS | GOCC:0110165 | Cellular anatomical entity |
| DLGAP1 | 9606.ENSP00000316377 | Monarch | EFO:0000246 | Age |
| DLGAP1 | 9606.ENSP00000316377 | Monarch | EFO:0000719 | Temporal measurement |
| DLGAP1 | 9606.ENSP00000316377 | Monarch | EFO:0003917 | Premature birth |
| DLGAP1 | 9606.ENSP00000316377 | Monarch | EFO:0004298 | Cardiovascular measurement |
| DLGAP1 | 9606.ENSP00000316377 | Monarch | EFO:0004302 | Anthropometric measurement |
| DLGAP1 | 9606.ENSP00000316377 | Monarch | EFO:0004306 | Erythrocyte indices |
| DLGAP1 | 9606.ENSP00000316377 | Monarch | EFO:0004311 | Heart function measurement |
| DLGAP1 | 9606.ENSP00000316377 | Monarch | EFO:0004324 | Body weights and measures |
| DLGAP1 | 9606.ENSP00000316377 | Monarch | EFO:0004339 | Body height |
| DLGAP1 | 9606.ENSP00000316377 | Monarch | EFO:0004503 | Hematological measurement |
| DLGAP1 | 9606.ENSP00000316377 | Monarch | EFO:0004512 | Bone measurement |
| DLGAP1 | 9606.ENSP00000316377 | Monarch | EFO:0004526 | Mean corpuscular volume |
| DLGAP1 | 9606.ENSP00000316377 | Monarch | EFO:0004529 | Lipid measurement |
| DLGAP1 | 9606.ENSP00000316377 | Monarch | EFO:0004557 | Population measurement |
| DLGAP1 | 9606.ENSP00000316377 | Monarch | EFO:0004582 | Liver enzyme measurement |
| DLGAP1 | 9606.ENSP00000316377 | Monarch | EFO:0004626 | IGFBP-3 measurement |
| DLGAP1 | 9606.ENSP00000316377 | Monarch | EFO:0004703 | Age at menarche |
| DLGAP1 | 9606.ENSP00000316377 | Monarch | EFO:0004730 | Hormone measurement |
| DLGAP1 | 9606.ENSP00000316377 | Monarch | EFO:0004735 | Serum alanine aminotransferase measurement |
| DLGAP1 | 9606.ENSP00000316377 | Monarch | EFO:0004742 | Renal system measurement |
| DLGAP1 | 9606.ENSP00000316377 | Monarch | EFO:0004747 | Protein measurement |
| DLGAP1 | 9606.ENSP00000316377 | Monarch | EFO:0004827 | Economic and social preference |
| DLGAP1 | 9606.ENSP00000316377 | Monarch | EFO:0004908 | Testosterone measurement |
| DLGAP1 | 9606.ENSP00000316377 | Monarch | EFO:0005043 | Cardiac troponin T measurement |
| DLGAP1 | 9606.ENSP00000316377 | Monarch | EFO:0005047 | Erythrocyte measurement |
| DLGAP1 | 9606.ENSP00000316377 | Monarch | EFO:0005093 | Hip circumference |
| DLGAP1 | 9606.ENSP00000316377 | Monarch | EFO:0005105 | Lipid or lipoprotein measurement |
| DLGAP1 | 9606.ENSP00000316377 | Monarch | EFO:0005127 | Cancer biomarker measurement |

| | | | | |
|---|---|---|---|---|
| DLGAP1 | 9606.ENSP00000316377 | Monarch | EFO:0005134 | Amino acid measurement |
| DLGAP1 | 9606.ENSP00000316377 | Monarch | EFO:0005278 | Cardiovascular disease biomarker measurement |
| DLGAP1 | 9606.ENSP00000316377 | Monarch | EFO:0005409 | Fat body mass |
| DLGAP1 | 9606.ENSP00000316377 | Monarch | EFO:0005526 | Response to alcohol |
| DLGAP1 | 9606.ENSP00000316377 | Monarch | EFO:0005527 | Ejection fraction measurement |
| DLGAP1 | 9606.ENSP00000316377 | Monarch | EFO:0005671 | Smoking behaviour measurement |
| DLGAP1 | 9606.ENSP00000316377 | Monarch | EFO:0005677 | Puberty onset measurement |
| DLGAP1 | 9606.ENSP00000316377 | Monarch | EFO:0006527 | Smoking status measurement |
| DLGAP1 | 9606.ENSP00000316377 | Monarch | EFO:0006917 | Spontaneous preterm birth |
| DLGAP1 | 9606.ENSP00000316377 | Monarch | EFO:0007042 | Polychlorinated biphenyls measurement |
| DLGAP1 | 9606.ENSP00000316377 | Monarch | EFO:0007807 | Erythrocyte cadmium measurement |
| DLGAP1 | 9606.ENSP00000316377 | Monarch | EFO:0007841 | Facial morphology measurement |
| DLGAP1 | 9606.ENSP00000316377 | Monarch | EFO:0007845 | Lip morphology measurement |
| DLGAP1 | 9606.ENSP00000316377 | Monarch | EFO:0007874 | Gut microbiome measurement |
| DLGAP1 | 9606.ENSP00000316377 | Monarch | EFO:0007882 | Microbiome measurement |
| DLGAP1 | 9606.ENSP00000316377 | Monarch | EFO:0007961 | Polybrominated biphenyl measurement |
| DLGAP1 | 9606.ENSP00000316377 | Monarch | EFO:0007962 | Polybrominated diphenyl ether measurement |
| DLGAP1 | 9606.ENSP00000316377 | Monarch | EFO:0007964 | Gestational serum measurement |
| DLGAP1 | 9606.ENSP00000316377 | Monarch | EFO:0008039 | BMI-adjusted hip circumference |
| DLGAP1 | 9606.ENSP00000316377 | Monarch | EFO:0009360 | Eyelid sagging measurement |
| DLGAP1 | 9606.ENSP00000316377 | Monarch | EFO:0009765 | Alanine measurement |
| DLGAP1 | 9606.ENSP00000316377 | Monarch | EFO:0009795 | Serum urea measurement |
| DLGAP1 | 9606.ENSP00000316377 | Monarch | EFO:0010224 | Lysophosphatidylcholine measurement |
| DLGAP1 | 9606.ENSP00000316377 | Monarch | EFO:0010362 | Lysophosphatidylcholine 20:3 measurement |
| DLGAP1 | 9606.ENSP00000316377 | Monarch | EFO:0010700 | Reticulocyte measurement |
| DLGAP1 | 9606.ENSP00000316377 | Monarch | EFO:0010701 | Mean reticulocyte volume |
| DLGAP1 | 9606.ENSP00000316377 | Monarch | EFO:0010948 | Lower face morphology measurement |
| DLGAP1 | 9606.ENSP00000316377 | Monarch | EFO:0010949 | Upper face morphology measurement |
| DLGAP1 | 9606.ENSP00000316377 | Monarch | EFO:0010968 | Phosphate measurement |
| DLGAP1 | 9606.ENSP00000316377 | Monarch | EFO:0011008 | Sex hormone measurement |
| DLGAP1 | 9606.ENSP00000316377 | Monarch | HP:0000118 | Phenotypic abnormality |
| DLGAP1 | 9606.ENSP00000316377 | Monarch | HP:0000951 | Abnormality of the skin |
| DLGAP1 | 9606.ENSP00000316377 | Monarch | HP:0000964 | Eczema |
| DLGAP1 | 9606.ENSP00000316377 | Monarch | HP:0001574 | Abnormality of the integument |
| DLGAP1 | 9606.ENSP00000316377 | Monarch | HP:0002715 | Abnormality of the immune system |

| | | | | |
|---|---|---|---|---|
| DLGAP1 | 9606.ENSP00000316377 | Monarch | HP:0010978 | Abnormality of immune system physiology |
| DLGAP1 | 9606.ENSP00000316377 | Monarch | HP:0011122 | Abnormality of skin physiology |
| DLGAP1 | 9606.ENSP00000316377 | Monarch | HP:0011123 | Inflammatory abnormality of the skin |
| DLGAP1 | 9606.ENSP00000316377 | Monarch | HP:0012647 | Abnormal inflammatory response |
| DLGAP1 | 9606.ENSP00000316377 | Monarch | HP:0012649 | Increased inflammatory response |
| DLGAP1 | 9606.ENSP00000316377 | UniProt Keywords | KW-0025 | Alternative splicing |
| DLGAP1 | 9606.ENSP00000316377 | UniProt Keywords | KW-0472 | Membrane |
| DLGAP1 | 9606.ENSP00000316377 | UniProt Keywords | KW-0597 | Phosphoprotein |
| DLGAP1 | 9606.ENSP00000316377 | UniProt Keywords | KW-0770 | Synapse |
| DLGAP1 | 9606.ENSP00000316377 | UniProt Keywords | KW-0965 | Cell junction |
| DLGAP1 | 9606.ENSP00000316377 | UniProt Keywords | KW-1003 | Cell membrane |
| DLGAP1 | 9606.ENSP00000316377 | Pfam | PF03359 | Guanylate-kinase-associated protein (GKAP) protein |
| DLGAP1 | 9606.ENSP00000316377 | InterPro | IPR005026 | SAPAP family |
| FOXN3 | 9606.ENSP00000343288 | GO Process | GO:0000075 | Cell cycle checkpoint signaling |
| FOXN3 | 9606.ENSP00000343288 | GO Process | GO:0000077 | DNA damage checkpoint signaling |
| FOXN3 | 9606.ENSP00000343288 | GO Process | GO:0000278 | Mitotic cell cycle |
| FOXN3 | 9606.ENSP00000343288 | GO Process | GO:0001501 | Skeletal system development |
| FOXN3 | 9606.ENSP00000343288 | GO Process | GO:0006355 | Regulation of transcription, DNA-templated |
| FOXN3 | 9606.ENSP00000343288 | GO Process | GO:0006357 | Regulation of transcription by RNA polymerase II |
| FOXN3 | 9606.ENSP00000343288 | GO Process | GO:0006950 | Response to stress |
| FOXN3 | 9606.ENSP00000343288 | GO Process | GO:0006974 | Cellular response to DNA damage stimulus |
| FOXN3 | 9606.ENSP00000343288 | GO Process | GO:0007049 | Cell cycle |
| FOXN3 | 9606.ENSP00000343288 | GO Process | GO:0007093 | Mitotic cell cycle checkpoint signaling |
| FOXN3 | 9606.ENSP00000343288 | GO Process | GO:0007095 | Mitotic G2 DNA damage checkpoint signaling |
| FOXN3 | 9606.ENSP00000343288 | GO Process | GO:0007154 | Cell communication |
| FOXN3 | 9606.ENSP00000343288 | GO Process | GO:0007165 | Signal transduction |
| FOXN3 | 9606.ENSP00000343288 | GO Process | GO:0007275 | Multicellular organism development |
| FOXN3 | 9606.ENSP00000343288 | GO Process | GO:0007346 | Regulation of mitotic cell cycle |
| FOXN3 | 9606.ENSP00000343288 | GO Process | GO:0009653 | Anatomical structure morphogenesis |
| FOXN3 | 9606.ENSP00000343288 | GO Process | GO:0009887 | Animal organ morphogenesis |
| FOXN3 | 9606.ENSP00000343288 | GO Process | GO:0009889 | Regulation of biosynthetic process |
| FOXN3 | 9606.ENSP00000343288 | GO Process | GO:0009890 | Negative regulation of biosynthetic process |
| FOXN3 | 9606.ENSP00000343288 | GO Process | GO:0009892 | Negative regulation of metabolic process |
| FOXN3 | 9606.ENSP00000343288 | GO Process | GO:0009987 | Cellular process |
| FOXN3 | 9606.ENSP00000343288 | GO Process | GO:0010389 | Regulation of G2/M transition of mitotic cell cycle |

| | | | | |
|---|---|---|---|---|
| FOXN3 | 9606.ENSP00000343288 | GO Process | GO:0010468 | Regulation of gene expression |
| FOXN3 | 9606.ENSP00000343288 | GO Process | GO:0010556 | Regulation of macromolecule biosynthetic process |
| FOXN3 | 9606.ENSP00000343288 | GO Process | GO:0010558 | Negative regulation of macromolecule biosynthetic process |
| FOXN3 | 9606.ENSP00000343288 | GO Process | GO:0010564 | Regulation of cell cycle process |
| FOXN3 | 9606.ENSP00000343288 | GO Process | GO:0010605 | Negative regulation of macromolecule metabolic process |
| FOXN3 | 9606.ENSP00000343288 | GO Process | GO:0010948 | Negative regulation of cell cycle process |
| FOXN3 | 9606.ENSP00000343288 | GO Process | GO:0010972 | Negative regulation of G2/M transition of mitotic cell cycle |
| FOXN3 | 9606.ENSP00000343288 | GO Process | GO:0019219 | Regulation of nucleobase-containing compound metabolic process |
| FOXN3 | 9606.ENSP00000343288 | GO Process | GO:0019222 | Regulation of metabolic process |
| FOXN3 | 9606.ENSP00000343288 | GO Process | GO:0022402 | Cell cycle process |
| FOXN3 | 9606.ENSP00000343288 | GO Process | GO:0023052 | Signaling |
| FOXN3 | 9606.ENSP00000343288 | GO Process | GO:0031323 | Regulation of cellular metabolic process |
| FOXN3 | 9606.ENSP00000343288 | GO Process | GO:0031324 | Negative regulation of cellular metabolic process |
| FOXN3 | 9606.ENSP00000343288 | GO Process | GO:0031326 | Regulation of cellular biosynthetic process |
| FOXN3 | 9606.ENSP00000343288 | GO Process | GO:0031327 | Negative regulation of cellular biosynthetic process |
| FOXN3 | 9606.ENSP00000343288 | GO Process | GO:0031570 | DNA integrity checkpoint signaling |
| FOXN3 | 9606.ENSP00000343288 | GO Process | GO:0032501 | Multicellular organismal process |
| FOXN3 | 9606.ENSP00000343288 | GO Process | GO:0032502 | Developmental process |
| FOXN3 | 9606.ENSP00000343288 | GO Process | GO:0033554 | Cellular response to stress |
| FOXN3 | 9606.ENSP00000343288 | GO Process | GO:0035556 | Intracellular signal transduction |
| FOXN3 | 9606.ENSP00000343288 | GO Process | GO:0042770 | Signal transduction in response to DNA damage |
| FOXN3 | 9606.ENSP00000343288 | GO Process | GO:0044773 | Mitotic DNA damage checkpoint signaling |
| FOXN3 | 9606.ENSP00000343288 | GO Process | GO:0044774 | Mitotic DNA integrity checkpoint signaling |
| FOXN3 | 9606.ENSP00000343288 | GO Process | GO:0044818 | Mitotic G2/M transition checkpoint |
| FOXN3 | 9606.ENSP00000343288 | GO Process | GO:0045786 | Negative regulation of cell cycle |
| FOXN3 | 9606.ENSP00000343288 | GO Process | GO:0045892 | Negative regulation of transcription, DNA-templated |
| FOXN3 | 9606.ENSP00000343288 | GO Process | GO:0045930 | Negative regulation of mitotic cell cycle |
| FOXN3 | 9606.ENSP00000343288 | GO Process | GO:0045934 | Negative regulation of nucleobase-containing compound metabolic process |
| FOXN3 | 9606.ENSP00000343288 | GO Process | GO:0048513 | Animal organ development |
| FOXN3 | 9606.ENSP00000343288 | GO Process | GO:0048519 | Negative regulation of biological process |
| FOXN3 | 9606.ENSP00000343288 | GO Process | GO:0048523 | Negative regulation of cellular process |
| FOXN3 | 9606.ENSP00000343288 | GO Process | GO:0048705 | Skeletal system morphogenesis |
| FOXN3 | 9606.ENSP00000343288 | GO Process | GO:0048731 | System development |
| FOXN3 | 9606.ENSP00000343288 | GO Process | GO:0048856 | Anatomical structure development |

| | | | | |
|---|---|---|---|---|
| FOXN3 | 9606.ENSP00000343288 | GO Process | GO:0050789 | Regulation of biological process |
| FOXN3 | 9606.ENSP00000343288 | GO Process | GO:0050794 | Regulation of cellular process |
| FOXN3 | 9606.ENSP00000343288 | GO Process | GO:0050896 | Response to stimulus |
| FOXN3 | 9606.ENSP00000343288 | GO Process | GO:0051171 | Regulation of nitrogen compound metabolic process |
| FOXN3 | 9606.ENSP00000343288 | GO Process | GO:0051172 | Negative regulation of nitrogen compound metabolic process |
| FOXN3 | 9606.ENSP00000343288 | GO Process | GO:0051252 | Regulation of RNA metabolic process |
| FOXN3 | 9606.ENSP00000343288 | GO Process | GO:0051253 | Negative regulation of RNA metabolic process |
| FOXN3 | 9606.ENSP00000343288 | GO Process | GO:0051716 | Cellular response to stimulus |
| FOXN3 | 9606.ENSP00000343288 | GO Process | GO:0051726 | Regulation of cell cycle |
| FOXN3 | 9606.ENSP00000343288 | GO Process | GO:0060255 | Regulation of macromolecule metabolic process |
| FOXN3 | 9606.ENSP00000343288 | GO Process | GO:0060348 | Bone development |
| FOXN3 | 9606.ENSP00000343288 | GO Process | GO:0060349 | Bone morphogenesis |
| FOXN3 | 9606.ENSP00000343288 | GO Process | GO:0065007 | Biological regulation |
| FOXN3 | 9606.ENSP00000343288 | GO Process | GO:0080090 | Regulation of primary metabolic process |
| FOXN3 | 9606.ENSP00000343288 | GO Process | GO:0097094 | Craniofacial suture morphogenesis |
| FOXN3 | 9606.ENSP00000343288 | GO Process | GO:1901987 | Regulation of cell cycle phase transition |
| FOXN3 | 9606.ENSP00000343288 | GO Process | GO:1901988 | Negative regulation of cell cycle phase transition |
| FOXN3 | 9606.ENSP00000343288 | GO Process | GO:1901990 | Regulation of mitotic cell cycle phase transition |
| FOXN3 | 9606.ENSP00000343288 | GO Process | GO:1901991 | Negative regulation of mitotic cell cycle phase transition |
| FOXN3 | 9606.ENSP00000343288 | GO Process | GO:1902679 | Negative regulation of RNA biosynthetic process |
| FOXN3 | 9606.ENSP00000343288 | GO Process | GO:1902749 | Regulation of cell cycle G2/M phase transition |
| FOXN3 | 9606.ENSP00000343288 | GO Process | GO:1902750 | Negative regulation of cell cycle G2/M phase transition |
| FOXN3 | 9606.ENSP00000343288 | GO Process | GO:1903047 | Mitotic cell cycle process |
| FOXN3 | 9606.ENSP00000343288 | GO Process | GO:1903506 | Regulation of nucleic acid-templated transcription |
| FOXN3 | 9606.ENSP00000343288 | GO Process | GO:1903507 | Negative regulation of nucleic acid-templated transcription |
| FOXN3 | 9606.ENSP00000343288 | GO Process | GO:1904888 | Cranial skeletal system development |
| FOXN3 | 9606.ENSP00000343288 | GO Process | GO:2001141 | Regulation of RNA biosynthetic process |
| FOXN3 | 9606.ENSP00000343288 | GO Function | GO:0000976 | Transcription cis-regulatory region binding |
| FOXN3 | 9606.ENSP00000343288 | GO Function | GO:0000981 | DNA-binding transcription factor activity, RNA polymerase II-specific |
| FOXN3 | 9606.ENSP00000343288 | GO Function | GO:0000987 | Cis-regulatory region sequence-specific DNA binding |
| FOXN3 | 9606.ENSP00000343288 | GO Function | GO:0001067 | Transcription regulatory region nucleic acid binding |
| FOXN3 | 9606.ENSP00000343288 | GO Function | GO:0003676 | Nucleic acid binding |
| FOXN3 | 9606.ENSP00000343288 | GO Function | GO:0003677 | DNA binding |
| FOXN3 | 9606.ENSP00000343288 | GO Function | GO:0003690 | Double-stranded DNA binding |
| FOXN3 | 9606.ENSP00000343288 | GO Function | GO:0003700 | DNA-binding transcription factor activity |

| | | | | |
|---|---|---|---|---|
| FOXN3 | 9606.ENSP00000343288 | GO Function | GO:0005488 | Binding |
| FOXN3 | 9606.ENSP00000343288 | GO Function | GO:0005515 | Protein binding |
| FOXN3 | 9606.ENSP00000343288 | GO Function | GO:0008022 | Protein C-terminus binding |
| FOXN3 | 9606.ENSP00000343288 | GO Function | GO:0043565 | Sequence-specific DNA binding |
| FOXN3 | 9606.ENSP00000343288 | GO Function | GO:0097159 | Organic cyclic compound binding |
| FOXN3 | 9606.ENSP00000343288 | GO Function | GO:0140110 | Transcription regulator activity |
| FOXN3 | 9606.ENSP00000343288 | GO Function | GO:1901363 | Heterocyclic compound binding |
| FOXN3 | 9606.ENSP00000343288 | GO Function | GO:1990837 | Sequence-specific double-stranded DNA binding |
| FOXN3 | 9606.ENSP00000343288 | GO Component | GO:0000785 | Chromatin |
| FOXN3 | 9606.ENSP00000343288 | GO Component | GO:0005622 | Intracellular anatomical structure |
| FOXN3 | 9606.ENSP00000343288 | GO Component | GO:0005634 | Nucleus |
| FOXN3 | 9606.ENSP00000343288 | GO Component | GO:0005694 | Chromosome |
| FOXN3 | 9606.ENSP00000343288 | GO Component | GO:0043226 | Organelle |
| FOXN3 | 9606.ENSP00000343288 | GO Component | GO:0043227 | Membrane-bounded organelle |
| FOXN3 | 9606.ENSP00000343288 | GO Component | GO:0043228 | Non-membrane-bounded organelle |
| FOXN3 | 9606.ENSP00000343288 | GO Component | GO:0043229 | Intracellular organelle |
| FOXN3 | 9606.ENSP00000343288 | GO Component | GO:0043231 | Intracellular membrane-bounded organelle |
| FOXN3 | 9606.ENSP00000343288 | GO Component | GO:0043232 | Intracellular non-membrane-bounded organelle |
| FOXN3 | 9606.ENSP00000343288 | GO Component | GO:0110165 | Cellular anatomical entity |
| FOXN3 | 9606.ENSP00000343288 | STRING clusters | CL:21575 | Mixed, incl. FORKHEAD, and GCM motif protein |
| FOXN3 | 9606.ENSP00000343288 | STRING clusters | CL:21577 | Mixed, incl. FORKHEAD, and Trichodontoosseous syndrome |
| FOXN3 | 9606.ENSP00000343288 | STRING clusters | CL:21603 | Mixed, incl. Forkhead box protein B1/B2, forkhead domain, and Forkhead box protein N2-4-like |
| FOXN3 | 9606.ENSP00000343288 | WikiPathways | WP2853 | Endoderm differentiation |
| FOXN3 | 9606.ENSP00000343288 | WikiPathways | WP4754 | IL-18 signaling pathway |
| FOXN3 | 9606.ENSP00000343288 | TISSUES | BTO:0000000 | Tissues, cell types and enzyme sources |
| FOXN3 | 9606.ENSP00000343288 | TISSUES | BTO:0000042 | Animal |
| FOXN3 | 9606.ENSP00000343288 | TISSUES | BTO:0000580 | Blood cancer cell |
| FOXN3 | 9606.ENSP00000343288 | TISSUES | BTO:0000744 | Lymphocytic leukemia cell |
| FOXN3 | 9606.ENSP00000343288 | TISSUES | BTO:0000887 | Muscle |
| FOXN3 | 9606.ENSP00000343288 | TISSUES | BTO:0001271 | Leukemia cell |
| FOXN3 | 9606.ENSP00000343288 | TISSUES | BTO:0001485 | Muscular system |
| FOXN3 | 9606.ENSP00000343288 | TISSUES | BTO:0001489 | Whole body |
| FOXN3 | 9606.ENSP00000343288 | TISSUES | BTO:0001546 | Chronic lymphocytic leukemia cell |
| FOXN3 | 9606.ENSP00000343288 | TISSUES | BTO:0003505 | T-cell chronic lymphocytic leukemia cell |

| | | | | |
|---|---|---|---|---|
| FOXN3 | 9606.ENSP00000343288 | Monarch | EFO:0003923 | Bone density |
| FOXN3 | 9606.ENSP00000343288 | Monarch | EFO:0003925 | Cognition |
| FOXN3 | 9606.ENSP00000343288 | Monarch | EFO:0004298 | Cardiovascular measurement |
| FOXN3 | 9606.ENSP00000343288 | Monarch | EFO:0004302 | Anthropometric measurement |
| FOXN3 | 9606.ENSP00000343288 | Monarch | EFO:0004311 | Heart function measurement |
| FOXN3 | 9606.ENSP00000343288 | Monarch | EFO:0004323 | Mental process |
| FOXN3 | 9606.ENSP00000343288 | Monarch | EFO:0004324 | Body weights and measures |
| FOXN3 | 9606.ENSP00000343288 | Monarch | EFO:0004327 | Electrocardiography |
| FOXN3 | 9606.ENSP00000343288 | Monarch | EFO:0004337 | Intelligence |
| FOXN3 | 9606.ENSP00000343288 | Monarch | EFO:0004340 | Body mass index |
| FOXN3 | 9606.ENSP00000343288 | Monarch | EFO:0004464 | Brain measurement |
| FOXN3 | 9606.ENSP00000343288 | Monarch | EFO:0004465 | Fasting blood glucose measurement |
| FOXN3 | 9606.ENSP00000343288 | Monarch | EFO:0004468 | Glucose measurement |
| FOXN3 | 9606.ENSP00000343288 | Monarch | EFO:0004503 | Hematological measurement |
| FOXN3 | 9606.ENSP00000343288 | Monarch | EFO:0004509 | Hemoglobin measurement |
| FOXN3 | 9606.ENSP00000343288 | Monarch | EFO:0004512 | Bone measurement |
| FOXN3 | 9606.ENSP00000343288 | Monarch | EFO:0004516 | Bone fracture related measurement |
| FOXN3 | 9606.ENSP00000343288 | Monarch | EFO:0004529 | Lipid measurement |
| FOXN3 | 9606.ENSP00000343288 | Monarch | EFO:0004530 | Triglyceride measurement |
| FOXN3 | 9606.ENSP00000343288 | Monarch | EFO:0004541 | HbA1c measurement |
| FOXN3 | 9606.ENSP00000343288 | Monarch | EFO:0004555 | Glycoprotein measurement |
| FOXN3 | 9606.ENSP00000343288 | Monarch | EFO:0004612 | High density lipoprotein cholesterol measurement |
| FOXN3 | 9606.ENSP00000343288 | Monarch | EFO:0004696 | Sex hormone-binding globulin measurement |
| FOXN3 | 9606.ENSP00000343288 | Monarch | EFO:0004730 | Hormone measurement |
| FOXN3 | 9606.ENSP00000343288 | Monarch | EFO:0004732 | Lipoprotein measurement |
| FOXN3 | 9606.ENSP00000343288 | Monarch | EFO:0004747 | Protein measurement |
| FOXN3 | 9606.ENSP00000343288 | Monarch | EFO:0004784 | Self reported educational attainment |
| FOXN3 | 9606.ENSP00000343288 | Monarch | EFO:0004875 | Mathematical ability |
| FOXN3 | 9606.ENSP00000343288 | Monarch | EFO:0004908 | Testosterone measurement |
| FOXN3 | 9606.ENSP00000343288 | Monarch | EFO:0005052 | Nervous system measurement |
| FOXN3 | 9606.ENSP00000343288 | Monarch | EFO:0005105 | Lipid or lipoprotein measurement |
| FOXN3 | 9606.ENSP00000343288 | Monarch | EFO:0005245 | Body weight loss |
| FOXN3 | 9606.ENSP00000343288 | Monarch | EFO:0005278 | Cardiovascular disease biomarker measurement |
| FOXN3 | 9606.ENSP00000343288 | Monarch | EFO:0005670 | Smoking initiation |
| FOXN3 | 9606.ENSP00000343288 | Monarch | EFO:0005671 | Smoking behaviour measurement |

| | | | | |
|---|---|---|---|---|
| FOXN3 | 9606.ENSP00000343288 | Monarch | EFO:0006527 | Smoking status measurement |
| FOXN3 | 9606.ENSP00000343288 | Monarch | EFO:0006842 | Diabetes mellitus biomarker |
| FOXN3 | 9606.ENSP00000343288 | Monarch | EFO:0006848 | Mental or behavioural disorder biomarker |
| FOXN3 | 9606.ENSP00000343288 | Monarch | EFO:0006930 | Brain volume measurement |
| FOXN3 | 9606.ENSP00000343288 | Monarch | EFO:0007655 | Clusterin measurement |
| FOXN3 | 9606.ENSP00000343288 | Monarch | EFO:0007657 | Cerebrospinal fluid clusterin measurement |
| FOXN3 | 9606.ENSP00000343288 | Monarch | EFO:0007860 | ADHD symptom measurement |
| FOXN3 | 9606.ENSP00000343288 | Monarch | EFO:0008354 | Cognitive function measurement |
| FOXN3 | 9606.ENSP00000343288 | Monarch | EFO:0008394 | Verbal-numerical reasoning measurement |
| FOXN3 | 9606.ENSP00000343288 | Monarch | EFO:0008579 | Risk-taking behaviour |
| FOXN3 | 9606.ENSP00000343288 | Monarch | EFO:0009270 | Heel bone mineral density |
| FOXN3 | 9606.ENSP00000343288 | Monarch | EFO:0010287 | Brain cortex volume measurement |
| FOXN3 | 9606.ENSP00000343288 | Monarch | EFO:0010309 | Insular cortex volume measurement |
| FOXN3 | 9606.ENSP00000343288 | Monarch | EFO:0010399 | Triacylglycerol 44:1 measurement |
| FOXN3 | 9606.ENSP00000343288 | Monarch | EFO:0011008 | Sex hormone measurement |
| FOXN3 | 9606.ENSP00000343288 | Monarch | EFO:0011015 | Educational attainment |
| FOXN3 | 9606.ENSP00000343288 | Monarch | HP:0000118 | Phenotypic abnormality |
| FOXN3 | 9606.ENSP00000343288 | Monarch | HP:0001392 | Abnormality of the liver |
| FOXN3 | 9606.ENSP00000343288 | Monarch | HP:0001395 | Hepatic fibrosis |
| FOXN3 | 9606.ENSP00000343288 | Monarch | HP:0001507 | Growth abnormality |
| FOXN3 | 9606.ENSP00000343288 | Monarch | HP:0002012 | Abnormality of the abdominal organs |
| FOXN3 | 9606.ENSP00000343288 | Monarch | HP:0004323 | Abnormality of body weight |
| FOXN3 | 9606.ENSP00000343288 | Monarch | HP:0025031 | Abnormality of the digestive system |
| FOXN3 | 9606.ENSP00000343288 | Monarch | HP:0410042 | Abnormal liver morphology |
| FOXN3 | 9606.ENSP00000343288 | UniProt Keywords | KW-0025 | Alternative splicing |
| FOXN3 | 9606.ENSP00000343288 | UniProt Keywords | KW-0131 | Cell cycle |
| FOXN3 | 9606.ENSP00000343288 | UniProt Keywords | KW-0238 | DNA-binding |
| FOXN3 | 9606.ENSP00000343288 | UniProt Keywords | KW-0539 | Nucleus |
| FOXN3 | 9606.ENSP00000343288 | UniProt Keywords | KW-0597 | Phosphoprotein |
| FOXN3 | 9606.ENSP00000343288 | UniProt Keywords | KW-0678 | Repressor |
| FOXN3 | 9606.ENSP00000343288 | UniProt Keywords | KW-0804 | Transcription |
| FOXN3 | 9606.ENSP00000343288 | UniProt Keywords | KW-0805 | Transcription regulation |
| FOXN3 | 9606.ENSP00000343288 | InterPro | IPR001766 | Fork head domain |
| FOXN3 | 9606.ENSP00000343288 | InterPro | IPR018122 | Fork head domain conserved site1 |
| FOXN3 | 9606.ENSP00000343288 | InterPro | IPR030456 | Fork head domain conserved site 2 |

| | | | | |
|---|---|---|---|---|
| FOXN3 | 9606.ENSP00000343288 | InterPro | IPR036388 | Winged helix-like DNA-binding domain superfamily |
| FOXN3 | 9606.ENSP00000343288 | InterPro | IPR036390 | Winged helix DNA-binding domain superfamily |
| FOXN3 | 9606.ENSP00000343288 | InterPro | IPR047119 | Forkhead box protein N2-4-like |
| FOXN3 | 9606.ENSP00000343288 | InterPro | IPR047404 | Forkhead box protein N3, forkhead domain |
| FOXN3 | 9606.ENSP00000343288 | SMART | SM00339 | FORKHEAD |
| KDR | 9606.ENSP00000263923 | GO Process | GO:0000003 | Reproduction |
| KDR | 9606.ENSP00000263923 | GO Process | GO:0000165 | MAPK cascade |
| KDR | 9606.ENSP00000263923 | GO Process | GO:0001525 | Angiogenesis |
| KDR | 9606.ENSP00000263923 | GO Process | GO:0001541 | Ovarian follicle development |
| KDR | 9606.ENSP00000263923 | GO Process | GO:0001568 | Blood vessel development |
| KDR | 9606.ENSP00000263923 | GO Process | GO:0001569 | Branching involved in blood vessel morphogenesis |
| KDR | 9606.ENSP00000263923 | GO Process | GO:0001570 | Vasculogenesis |
| KDR | 9606.ENSP00000263923 | GO Process | GO:0001654 | Eye development |
| KDR | 9606.ENSP00000263923 | GO Process | GO:0001667 | Ameboidal-type cell migration |
| KDR | 9606.ENSP00000263923 | GO Process | GO:0001763 | Morphogenesis of a branching structure |
| KDR | 9606.ENSP00000263923 | GO Process | GO:0001894 | Tissue homeostasis |
| KDR | 9606.ENSP00000263923 | GO Process | GO:0001932 | Regulation of protein phosphorylation |
| KDR | 9606.ENSP00000263923 | GO Process | GO:0001934 | Positive regulation of protein phosphorylation |
| KDR | 9606.ENSP00000263923 | GO Process | GO:0001936 | Regulation of endothelial cell proliferation |
| KDR | 9606.ENSP00000263923 | GO Process | GO:0001938 | Positive regulation of endothelial cell proliferation |
| KDR | 9606.ENSP00000263923 | GO Process | GO:0001944 | Vasculature development |
| KDR | 9606.ENSP00000263923 | GO Process | GO:0001945 | Lymph vessel development |
| KDR | 9606.ENSP00000263923 | GO Process | GO:0001952 | Regulation of cell-matrix adhesion |
| KDR | 9606.ENSP00000263923 | GO Process | GO:0001954 | Positive regulation of cell-matrix adhesion |
| KDR | 9606.ENSP00000263923 | GO Process | GO:0002009 | Morphogenesis of an epithelium |
| KDR | 9606.ENSP00000263923 | GO Process | GO:0002040 | Sprouting angiogenesis |
| KDR | 9606.ENSP00000263923 | GO Process | GO:0002042 | Cell migration involved in sprouting angiogenesis |
| KDR | 9606.ENSP00000263923 | GO Process | GO:0002053 | Positive regulation of mesenchymal cell proliferation |
| KDR | 9606.ENSP00000263923 | GO Process | GO:0002064 | Epithelial cell development |
| KDR | 9606.ENSP00000263923 | GO Process | GO:0002070 | Epithelial cell maturation |
| KDR | 9606.ENSP00000263923 | GO Process | GO:0002244 | Hematopoietic progenitor cell differentiation |
| KDR | 9606.ENSP00000263923 | GO Process | GO:0002376 | Immune system process |
| KDR | 9606.ENSP00000263923 | GO Process | GO:0002520 | Immune system development |
| KDR | 9606.ENSP00000263923 | GO Process | GO:0003006 | Developmental process involved in reproduction |
| KDR | 9606.ENSP00000263923 | GO Process | GO:0003157 | Endocardium development |

| | | | | |
|---|---|---|---|---|
| KDR | 9606.ENSP00000263923 | GO Process | GO:0003158 | Endothelium development |
| KDR | 9606.ENSP00000263923 | GO Process | GO:0003254 | Regulation of membrane depolarization |
| KDR | 9606.ENSP00000263923 | GO Process | GO:0006468 | Protein phosphorylation |
| KDR | 9606.ENSP00000263923 | GO Process | GO:0006793 | Phosphorus metabolic process |
| KDR | 9606.ENSP00000263923 | GO Process | GO:0006796 | Phosphate-containing compound metabolic process |
| KDR | 9606.ENSP00000263923 | GO Process | GO:0006807 | Nitrogen compound metabolic process |
| KDR | 9606.ENSP00000263923 | GO Process | GO:0006950 | Response to stress |
| KDR | 9606.ENSP00000263923 | GO Process | GO:0007154 | Cell communication |
| KDR | 9606.ENSP00000263923 | GO Process | GO:0007165 | Signal transduction |
| KDR | 9606.ENSP00000263923 | GO Process | GO:0007166 | Cell surface receptor signaling pathway |
| KDR | 9606.ENSP00000263923 | GO Process | GO:0007167 | Enzyme-linked receptor protein signaling pathway |
| KDR | 9606.ENSP00000263923 | GO Process | GO:0007169 | Transmembrane receptor protein tyrosine kinase signaling pathway |
| KDR | 9606.ENSP00000263923 | GO Process | GO:0007275 | Multicellular organism development |
| KDR | 9606.ENSP00000263923 | GO Process | GO:0007423 | Sensory organ development |
| KDR | 9606.ENSP00000263923 | GO Process | GO:0007507 | Heart development |
| KDR | 9606.ENSP00000263923 | GO Process | GO:0007548 | Sex differentiation |
| KDR | 9606.ENSP00000263923 | GO Process | GO:0008152 | Metabolic process |
| KDR | 9606.ENSP00000263923 | GO Process | GO:0008283 | Cell population proliferation |
| KDR | 9606.ENSP00000263923 | GO Process | GO:0008284 | Positive regulation of cell population proliferation |
| KDR | 9606.ENSP00000263923 | GO Process | GO:0008360 | Regulation of cell shape |
| KDR | 9606.ENSP00000263923 | GO Process | GO:0008406 | Gonad development |
| KDR | 9606.ENSP00000263923 | GO Process | GO:0008585 | Female gonad development |
| KDR | 9606.ENSP00000263923 | GO Process | GO:0009611 | Response to wounding |
| KDR | 9606.ENSP00000263923 | GO Process | GO:0009653 | Anatomical structure morphogenesis |
| KDR | 9606.ENSP00000263923 | GO Process | GO:0009790 | Embryo development |
| KDR | 9606.ENSP00000263923 | GO Process | GO:0009791 | Post-embryonic development |
| KDR | 9606.ENSP00000263923 | GO Process | GO:0009886 | Post-embryonic animal morphogenesis |
| KDR | 9606.ENSP00000263923 | GO Process | GO:0009887 | Animal organ morphogenesis |
| KDR | 9606.ENSP00000263923 | GO Process | GO:0009888 | Tissue development |
| KDR | 9606.ENSP00000263923 | GO Process | GO:0009889 | Regulation of biosynthetic process |
| KDR | 9606.ENSP00000263923 | GO Process | GO:0009891 | Positive regulation of biosynthetic process |
| KDR | 9606.ENSP00000263923 | GO Process | GO:0009892 | Negative regulation of metabolic process |
| KDR | 9606.ENSP00000263923 | GO Process | GO:0009893 | Positive regulation of metabolic process |
| KDR | 9606.ENSP00000263923 | GO Process | GO:0009894 | Regulation of catabolic process |
| KDR | 9606.ENSP00000263923 | GO Process | GO:0009896 | Positive regulation of catabolic process |

| | | | | |
|---|---|---|---|---|
| KDR | 9606.ENSP00000263923 | GO Process | GO:0009966 | Regulation of signal transduction |
| KDR | 9606.ENSP00000263923 | GO Process | GO:0009967 | Positive regulation of signal transduction |
| KDR | 9606.ENSP00000263923 | GO Process | GO:0009987 | Cellular process |
| KDR | 9606.ENSP00000263923 | GO Process | GO:0010033 | Response to organic substance |
| KDR | 9606.ENSP00000263923 | GO Process | GO:0010035 | Response to inorganic substance |
| KDR | 9606.ENSP00000263923 | GO Process | GO:0010463 | Mesenchymal cell proliferation |
| KDR | 9606.ENSP00000263923 | GO Process | GO:0010464 | Regulation of mesenchymal cell proliferation |
| KDR | 9606.ENSP00000263923 | GO Process | GO:0010468 | Regulation of gene expression |
| KDR | 9606.ENSP00000263923 | GO Process | GO:0010506 | Regulation of autophagy |
| KDR | 9606.ENSP00000263923 | GO Process | GO:0010508 | Positive regulation of autophagy |
| KDR | 9606.ENSP00000263923 | GO Process | GO:0010556 | Regulation of macromolecule biosynthetic process |
| KDR | 9606.ENSP00000263923 | GO Process | GO:0010557 | Positive regulation of macromolecule biosynthetic process |
| KDR | 9606.ENSP00000263923 | GO Process | GO:0010562 | Positive regulation of phosphorus metabolic process |
| KDR | 9606.ENSP00000263923 | GO Process | GO:0010594 | Regulation of endothelial cell migration |
| KDR | 9606.ENSP00000263923 | GO Process | GO:0010595 | Positive regulation of endothelial cell migration |
| KDR | 9606.ENSP00000263923 | GO Process | GO:0010604 | Positive regulation of macromolecule metabolic process |
| KDR | 9606.ENSP00000263923 | GO Process | GO:0010605 | Negative regulation of macromolecule metabolic process |
| KDR | 9606.ENSP00000263923 | GO Process | GO:0010629 | Negative regulation of gene expression |
| KDR | 9606.ENSP00000263923 | GO Process | GO:0010631 | Epithelial cell migration |
| KDR | 9606.ENSP00000263923 | GO Process | GO:0010632 | Regulation of epithelial cell migration |
| KDR | 9606.ENSP00000263923 | GO Process | GO:0010634 | Positive regulation of epithelial cell migration |
| KDR | 9606.ENSP00000263923 | GO Process | GO:0010638 | Positive regulation of organelle organization |
| KDR | 9606.ENSP00000263923 | GO Process | GO:0010646 | Regulation of cell communication |
| KDR | 9606.ENSP00000263923 | GO Process | GO:0010647 | Positive regulation of cell communication |
| KDR | 9606.ENSP00000263923 | GO Process | GO:0010810 | Regulation of cell-substrate adhesion |
| KDR | 9606.ENSP00000263923 | GO Process | GO:0010811 | Positive regulation of cell-substrate adhesion |
| KDR | 9606.ENSP00000263923 | GO Process | GO:0010821 | Regulation of mitochondrion organization |
| KDR | 9606.ENSP00000263923 | GO Process | GO:0010822 | Positive regulation of mitochondrion organization |
| KDR | 9606.ENSP00000263923 | GO Process | GO:0010941 | Regulation of cell death |
| KDR | 9606.ENSP00000263923 | GO Process | GO:0014066 | Regulation of phosphatidylinositol 3-kinase signaling |
| KDR | 9606.ENSP00000263923 | GO Process | GO:0014068 | Positive regulation of phosphatidylinositol 3-kinase signaling |
| KDR | 9606.ENSP00000263923 | GO Process | GO:0016239 | Positive regulation of macroautophagy |
| KDR | 9606.ENSP00000263923 | GO Process | GO:0016241 | Regulation of macroautophagy |
| KDR | 9606.ENSP00000263923 | GO Process | GO:0016310 | Phosphorylation |
| KDR | 9606.ENSP00000263923 | GO Process | GO:0016477 | Cell migration |

| | | | | |
|---|---|---|---|---|
| KDR | 9606.ENSP00000263923 | GO Process | GO:0018108 | Peptidyl-tyrosine phosphorylation |
| KDR | 9606.ENSP00000263923 | GO Process | GO:0018193 | Peptidyl-amino acid modification |
| KDR | 9606.ENSP00000263923 | GO Process | GO:0018212 | Peptidyl-tyrosine modification |
| KDR | 9606.ENSP00000263923 | GO Process | GO:0019220 | Regulation of phosphate metabolic process |
| KDR | 9606.ENSP00000263923 | GO Process | GO:0019222 | Regulation of metabolic process |
| KDR | 9606.ENSP00000263923 | GO Process | GO:0019538 | Protein metabolic process |
| KDR | 9606.ENSP00000263923 | GO Process | GO:0019722 | Calcium-mediated signaling |
| KDR | 9606.ENSP00000263923 | GO Process | GO:0019932 | Second-messenger-mediated signaling |
| KDR | 9606.ENSP00000263923 | GO Process | GO:0021700 | Developmental maturation |
| KDR | 9606.ENSP00000263923 | GO Process | GO:0022414 | Reproductive process |
| KDR | 9606.ENSP00000263923 | GO Process | GO:0022603 | Regulation of anatomical structure morphogenesis |
| KDR | 9606.ENSP00000263923 | GO Process | GO:0022604 | Regulation of cell morphogenesis |
| KDR | 9606.ENSP00000263923 | GO Process | GO:0023051 | Regulation of signaling |
| KDR | 9606.ENSP00000263923 | GO Process | GO:0023052 | Signaling |
| KDR | 9606.ENSP00000263923 | GO Process | GO:0023056 | Positive regulation of signaling |
| KDR | 9606.ENSP00000263923 | GO Process | GO:0030097 | Hemopoiesis |
| KDR | 9606.ENSP00000263923 | GO Process | GO:0030154 | Cell differentiation |
| KDR | 9606.ENSP00000263923 | GO Process | GO:0030155 | Regulation of cell adhesion |
| KDR | 9606.ENSP00000263923 | GO Process | GO:0030323 | Respiratory tube development |
| KDR | 9606.ENSP00000263923 | GO Process | GO:0030324 | Lung development |
| KDR | 9606.ENSP00000263923 | GO Process | GO:0030334 | Regulation of cell migration |
| KDR | 9606.ENSP00000263923 | GO Process | GO:0030335 | Positive regulation of cell migration |
| KDR | 9606.ENSP00000263923 | GO Process | GO:0030510 | Regulation of BMP signaling pathway |
| KDR | 9606.ENSP00000263923 | GO Process | GO:0030513 | Positive regulation of BMP signaling pathway |
| KDR | 9606.ENSP00000263923 | GO Process | GO:0030855 | Epithelial cell differentiation |
| KDR | 9606.ENSP00000263923 | GO Process | GO:0031077 | Post-embryonic camera-type eye development |
| KDR | 9606.ENSP00000263923 | GO Process | GO:0031323 | Regulation of cellular metabolic process |
| KDR | 9606.ENSP00000263923 | GO Process | GO:0031325 | Positive regulation of cellular metabolic process |
| KDR | 9606.ENSP00000263923 | GO Process | GO:0031329 | Regulation of cellular catabolic process |
| KDR | 9606.ENSP00000263923 | GO Process | GO:0031331 | Positive regulation of cellular catabolic process |
| KDR | 9606.ENSP00000263923 | GO Process | GO:0031399 | Regulation of protein modification process |
| KDR | 9606.ENSP00000263923 | GO Process | GO:0031401 | Positive regulation of protein modification process |
| KDR | 9606.ENSP00000263923 | GO Process | GO:0032101 | Regulation of response to external stimulus |
| KDR | 9606.ENSP00000263923 | GO Process | GO:0032103 | Positive regulation of response to external stimulus |
| KDR | 9606.ENSP00000263923 | GO Process | GO:0032501 | Multicellular organismal process |

| | | | | |
|---|---|---|---|---|
| KDR | 9606.ENSP00000263923 | GO Process | GO:0032502 | Developmental process |
| KDR | 9606.ENSP00000263923 | GO Process | GO:0033043 | Regulation of organelle organization |
| KDR | 9606.ENSP00000263923 | GO Process | GO:0033674 | Positive regulation of kinase activity |
| KDR | 9606.ENSP00000263923 | GO Process | GO:0035162 | Embryonic hemopoiesis |
| KDR | 9606.ENSP00000263923 | GO Process | GO:0035239 | Tube morphogenesis |
| KDR | 9606.ENSP00000263923 | GO Process | GO:0035295 | Tube development |
| KDR | 9606.ENSP00000263923 | GO Process | GO:0035556 | Intracellular signal transduction |
| KDR | 9606.ENSP00000263923 | GO Process | GO:0035584 | Calcium-mediated signaling using intracellular calcium source |
| KDR | 9606.ENSP00000263923 | GO Process | GO:0035924 | Cellular response to vascular endothelial growth factor stimulus |
| KDR | 9606.ENSP00000263923 | GO Process | GO:0036211 | Protein modification process |
| KDR | 9606.ENSP00000263923 | GO Process | GO:0036324 | Vascular endothelial growth factor receptor-2 signaling pathway |
| KDR | 9606.ENSP00000263923 | GO Process | GO:0038033 | Positive regulation of endothelial cell chemotaxis by VEGF-activated vascular endothelial growth factor receptor signaling pathway |
| KDR | 9606.ENSP00000263923 | GO Process | GO:0038083 | Peptidyl-tyrosine autophosphorylation |
| KDR | 9606.ENSP00000263923 | GO Process | GO:0038084 | Vascular endothelial growth factor signaling pathway |
| KDR | 9606.ENSP00000263923 | GO Process | GO:0038089 | Positive regulation of cell migration by vascular endothelial growth factor signaling pathway |
| KDR | 9606.ENSP00000263923 | GO Process | GO:0040012 | Regulation of locomotion |
| KDR | 9606.ENSP00000263923 | GO Process | GO:0040017 | Positive regulation of locomotion |
| KDR | 9606.ENSP00000263923 | GO Process | GO:0042060 | Wound healing |
| KDR | 9606.ENSP00000263923 | GO Process | GO:0042127 | Regulation of cell population proliferation |
| KDR | 9606.ENSP00000263923 | GO Process | GO:0042221 | Response to chemical |
| KDR | 9606.ENSP00000263923 | GO Process | GO:0042325 | Regulation of phosphorylation |
| KDR | 9606.ENSP00000263923 | GO Process | GO:0042327 | Positive regulation of phosphorylation |
| KDR | 9606.ENSP00000263923 | GO Process | GO:0042391 | Regulation of membrane potential |
| KDR | 9606.ENSP00000263923 | GO Process | GO:0042592 | Homeostatic process |
| KDR | 9606.ENSP00000263923 | GO Process | GO:0042981 | Regulation of apoptotic process |
| KDR | 9606.ENSP00000263923 | GO Process | GO:0043010 | Camera-type eye development |
| KDR | 9606.ENSP00000263923 | GO Process | GO:0043066 | Negative regulation of apoptotic process |
| KDR | 9606.ENSP00000263923 | GO Process | GO:0043067 | Regulation of programmed cell death |
| KDR | 9606.ENSP00000263923 | GO Process | GO:0043069 | Negative regulation of programmed cell death |
| KDR | 9606.ENSP00000263923 | GO Process | GO:0043085 | Positive regulation of catalytic activity |
| KDR | 9606.ENSP00000263923 | GO Process | GO:0043129 | Surfactant homeostasis |
| KDR | 9606.ENSP00000263923 | GO Process | GO:0043170 | Macromolecule metabolic process |
| KDR | 9606.ENSP00000263923 | GO Process | GO:0043408 | Regulation of MAPK cascade |

| | | | | |
|---|---|---|---|---|
| KDR | 9606.ENSP00000263923 | GO Process | GO:0043410 | Positive regulation of MAPK cascade |
| KDR | 9606.ENSP00000263923 | GO Process | GO:0043412 | Macromolecule modification |
| KDR | 9606.ENSP00000263923 | GO Process | GO:0043491 | Protein kinase B signaling |
| KDR | 9606.ENSP00000263923 | GO Process | GO:0043523 | Regulation of neuron apoptotic process |
| KDR | 9606.ENSP00000263923 | GO Process | GO:0043524 | Negative regulation of neuron apoptotic process |
| KDR | 9606.ENSP00000263923 | GO Process | GO:0043534 | Blood vessel endothelial cell migration |
| KDR | 9606.ENSP00000263923 | GO Process | GO:0043535 | Regulation of blood vessel endothelial cell migration |
| KDR | 9606.ENSP00000263923 | GO Process | GO:0043536 | Positive regulation of blood vessel endothelial cell migration |
| KDR | 9606.ENSP00000263923 | GO Process | GO:0043542 | Endothelial cell migration |
| KDR | 9606.ENSP00000263923 | GO Process | GO:0043549 | Regulation of kinase activity |
| KDR | 9606.ENSP00000263923 | GO Process | GO:0044087 | Regulation of cellular component biogenesis |
| KDR | 9606.ENSP00000263923 | GO Process | GO:0044089 | Positive regulation of cellular component biogenesis |
| KDR | 9606.ENSP00000263923 | GO Process | GO:0044093 | Positive regulation of molecular function |
| KDR | 9606.ENSP00000263923 | GO Process | GO:0044237 | Cellular metabolic process |
| KDR | 9606.ENSP00000263923 | GO Process | GO:0044238 | Primary metabolic process |
| KDR | 9606.ENSP00000263923 | GO Process | GO:0045137 | Development of primary sexual characteristics |
| KDR | 9606.ENSP00000263923 | GO Process | GO:0045165 | Cell fate commitment |
| KDR | 9606.ENSP00000263923 | GO Process | GO:0045446 | Endothelial cell differentiation |
| KDR | 9606.ENSP00000263923 | GO Process | GO:0045595 | Regulation of cell differentiation |
| KDR | 9606.ENSP00000263923 | GO Process | GO:0045597 | Positive regulation of cell differentiation |
| KDR | 9606.ENSP00000263923 | GO Process | GO:0045765 | Regulation of angiogenesis |
| KDR | 9606.ENSP00000263923 | GO Process | GO:0045766 | Positive regulation of angiogenesis |
| KDR | 9606.ENSP00000263923 | GO Process | GO:0045785 | Positive regulation of cell adhesion |
| KDR | 9606.ENSP00000263923 | GO Process | GO:0045937 | Positive regulation of phosphate metabolic process |
| KDR | 9606.ENSP00000263923 | GO Process | GO:0046545 | Development of primary female sexual characteristics |
| KDR | 9606.ENSP00000263923 | GO Process | GO:0046660 | Female sex differentiation |
| KDR | 9606.ENSP00000263923 | GO Process | GO:0046777 | Protein autophosphorylation |
| KDR | 9606.ENSP00000263923 | GO Process | GO:0048010 | Vascular endothelial growth factor receptor signaling pathway |
| KDR | 9606.ENSP00000263923 | GO Process | GO:0048050 | Post-embryonic eye morphogenesis |
| KDR | 9606.ENSP00000263923 | GO Process | GO:0048286 | Lung alveolus development |
| KDR | 9606.ENSP00000263923 | GO Process | GO:0048468 | Cell development |
| KDR | 9606.ENSP00000263923 | GO Process | GO:0048469 | Cell maturation |
| KDR | 9606.ENSP00000263923 | GO Process | GO:0048513 | Animal organ development |
| KDR | 9606.ENSP00000263923 | GO Process | GO:0048514 | Blood vessel morphogenesis |
| KDR | 9606.ENSP00000263923 | GO Process | GO:0048518 | Positive regulation of biological process |

| | | | | |
|---|---|---|---|---|
| KDR | 9606.ENSP00000263923 | GO Process | GO:0048519 | Negative regulation of biological process |
| KDR | 9606.ENSP00000263923 | GO Process | GO:0048522 | Positive regulation of cellular process |
| KDR | 9606.ENSP00000263923 | GO Process | GO:0048523 | Negative regulation of cellular process |
| KDR | 9606.ENSP00000263923 | GO Process | GO:0048534 | Hematopoietic or lymphoid organ development |
| KDR | 9606.ENSP00000263923 | GO Process | GO:0048563 | Post-embryonic animal organ morphogenesis |
| KDR | 9606.ENSP00000263923 | GO Process | GO:0048568 | Embryonic organ development |
| KDR | 9606.ENSP00000263923 | GO Process | GO:0048569 | Post-embryonic animal organ development |
| KDR | 9606.ENSP00000263923 | GO Process | GO:0048583 | Regulation of response to stimulus |
| KDR | 9606.ENSP00000263923 | GO Process | GO:0048584 | Positive regulation of response to stimulus |
| KDR | 9606.ENSP00000263923 | GO Process | GO:0048592 | Eye morphogenesis |
| KDR | 9606.ENSP00000263923 | GO Process | GO:0048593 | Camera-type eye morphogenesis |
| KDR | 9606.ENSP00000263923 | GO Process | GO:0048597 | Post-embryonic camera-type eye morphogenesis |
| KDR | 9606.ENSP00000263923 | GO Process | GO:0048608 | Reproductive structure development |
| KDR | 9606.ENSP00000263923 | GO Process | GO:0048646 | Anatomical structure formation involved in morphogenesis |
| KDR | 9606.ENSP00000263923 | GO Process | GO:0048729 | Tissue morphogenesis |
| KDR | 9606.ENSP00000263923 | GO Process | GO:0048731 | System development |
| KDR | 9606.ENSP00000263923 | GO Process | GO:0048754 | Branching morphogenesis of an epithelial tube |
| KDR | 9606.ENSP00000263923 | GO Process | GO:0048856 | Anatomical structure development |
| KDR | 9606.ENSP00000263923 | GO Process | GO:0048869 | Cellular developmental process |
| KDR | 9606.ENSP00000263923 | GO Process | GO:0048870 | Cell motility |
| KDR | 9606.ENSP00000263923 | GO Process | GO:0048871 | Multicellular organismal homeostasis |
| KDR | 9606.ENSP00000263923 | GO Process | GO:0048875 | Chemical homeostasis within a tissue |
| KDR | 9606.ENSP00000263923 | GO Process | GO:0048878 | Chemical homeostasis |
| KDR | 9606.ENSP00000263923 | GO Process | GO:0048880 | Sensory system development |
| KDR | 9606.ENSP00000263923 | GO Process | GO:0050673 | Epithelial cell proliferation |
| KDR | 9606.ENSP00000263923 | GO Process | GO:0050678 | Regulation of epithelial cell proliferation |
| KDR | 9606.ENSP00000263923 | GO Process | GO:0050679 | Positive regulation of epithelial cell proliferation |
| KDR | 9606.ENSP00000263923 | GO Process | GO:0050789 | Regulation of biological process |
| KDR | 9606.ENSP00000263923 | GO Process | GO:0050790 | Regulation of catalytic activity |
| KDR | 9606.ENSP00000263923 | GO Process | GO:0050793 | Regulation of developmental process |
| KDR | 9606.ENSP00000263923 | GO Process | GO:0050794 | Regulation of cellular process |
| KDR | 9606.ENSP00000263923 | GO Process | GO:0050801 | Ion homeostasis |
| KDR | 9606.ENSP00000263923 | GO Process | GO:0050896 | Response to stimulus |
| KDR | 9606.ENSP00000263923 | GO Process | GO:0050920 | Regulation of chemotaxis |
| KDR | 9606.ENSP00000263923 | GO Process | GO:0050921 | Positive regulation of chemotaxis |

| | | | | |
|---|---|---|---|---|
| KDR | 9606.ENSP00000263923 | GO Process | GO:0050926 | Regulation of positive chemotaxis |
| KDR | 9606.ENSP00000263923 | GO Process | GO:0050927 | Positive regulation of positive chemotaxis |
| KDR | 9606.ENSP00000263923 | GO Process | GO:0051094 | Positive regulation of developmental process |
| KDR | 9606.ENSP00000263923 | GO Process | GO:0051128 | Regulation of cellular component organization |
| KDR | 9606.ENSP00000263923 | GO Process | GO:0051130 | Positive regulation of cellular component organization |
| KDR | 9606.ENSP00000263923 | GO Process | GO:0051171 | Regulation of nitrogen compound metabolic process |
| KDR | 9606.ENSP00000263923 | GO Process | GO:0051173 | Positive regulation of nitrogen compound metabolic process |
| KDR | 9606.ENSP00000263923 | GO Process | GO:0051174 | Regulation of phosphorus metabolic process |
| KDR | 9606.ENSP00000263923 | GO Process | GO:0051239 | Regulation of multicellular organismal process |
| KDR | 9606.ENSP00000263923 | GO Process | GO:0051240 | Positive regulation of multicellular organismal process |
| KDR | 9606.ENSP00000263923 | GO Process | GO:0051246 | Regulation of protein metabolic process |
| KDR | 9606.ENSP00000263923 | GO Process | GO:0051247 | Positive regulation of protein metabolic process |
| KDR | 9606.ENSP00000263923 | GO Process | GO:0051338 | Regulation of transferase activity |
| KDR | 9606.ENSP00000263923 | GO Process | GO:0051347 | Positive regulation of transferase activity |
| KDR | 9606.ENSP00000263923 | GO Process | GO:0051716 | Cellular response to stimulus |
| KDR | 9606.ENSP00000263923 | GO Process | GO:0051769 | Regulation of nitric-oxide synthase biosynthetic process |
| KDR | 9606.ENSP00000263923 | GO Process | GO:0051770 | Positive regulation of nitric-oxide synthase biosynthetic process |
| KDR | 9606.ENSP00000263923 | GO Process | GO:0051881 | Regulation of mitochondrial membrane potential |
| KDR | 9606.ENSP00000263923 | GO Process | GO:0051893 | Regulation of focal adhesion assembly |
| KDR | 9606.ENSP00000263923 | GO Process | GO:0051894 | Positive regulation of focal adhesion assembly |
| KDR | 9606.ENSP00000263923 | GO Process | GO:0051900 | Regulation of mitochondrial depolarization |
| KDR | 9606.ENSP00000263923 | GO Process | GO:0051901 | Positive regulation of mitochondrial depolarization |
| KDR | 9606.ENSP00000263923 | GO Process | GO:0055065 | Metal ion homeostasis |
| KDR | 9606.ENSP00000263923 | GO Process | GO:0055074 | Calcium ion homeostasis |
| KDR | 9606.ENSP00000263923 | GO Process | GO:0055080 | Cation homeostasis |
| KDR | 9606.ENSP00000263923 | GO Process | GO:0060055 | Angiogenesis involved in wound healing |
| KDR | 9606.ENSP00000263923 | GO Process | GO:0060249 | Anatomical structure homeostasis |
| KDR | 9606.ENSP00000263923 | GO Process | GO:0060255 | Regulation of macromolecule metabolic process |
| KDR | 9606.ENSP00000263923 | GO Process | GO:0060429 | Epithelium development |
| KDR | 9606.ENSP00000263923 | GO Process | GO:0060541 | Respiratory system development |
| KDR | 9606.ENSP00000263923 | GO Process | GO:0060548 | Negative regulation of cell death |
| KDR | 9606.ENSP00000263923 | GO Process | GO:0060562 | Epithelial tube morphogenesis |
| KDR | 9606.ENSP00000263923 | GO Process | GO:0060837 | Blood vessel endothelial cell differentiation |
| KDR | 9606.ENSP00000263923 | GO Process | GO:0061042 | Vascular wound healing |
| KDR | 9606.ENSP00000263923 | GO Process | GO:0061138 | Morphogenesis of a branching epithelium |

| | | | | |
|---|---|---|---|---|
| KDR | 9606.ENSP00000263923 | GO Process | GO:0061458 | Reproductive system development |
| KDR | 9606.ENSP00000263923 | GO Process | GO:0065007 | Biological regulation |
| KDR | 9606.ENSP00000263923 | GO Process | GO:0065008 | Regulation of biological quality |
| KDR | 9606.ENSP00000263923 | GO Process | GO:0065009 | Regulation of molecular function |
| KDR | 9606.ENSP00000263923 | GO Process | GO:0070371 | ERK1 and ERK2 cascade |
| KDR | 9606.ENSP00000263923 | GO Process | GO:0070372 | Regulation of ERK1 and ERK2 cascade |
| KDR | 9606.ENSP00000263923 | GO Process | GO:0070374 | Positive regulation of ERK1 and ERK2 cascade |
| KDR | 9606.ENSP00000263923 | GO Process | GO:0070848 | Response to growth factor |
| KDR | 9606.ENSP00000263923 | GO Process | GO:0070887 | Cellular response to chemical stimulus |
| KDR | 9606.ENSP00000263923 | GO Process | GO:0071310 | Cellular response to organic substance |
| KDR | 9606.ENSP00000263923 | GO Process | GO:0071363 | Cellular response to growth factor stimulus |
| KDR | 9606.ENSP00000263923 | GO Process | GO:0071526 | Semaphorin-plexin signaling pathway |
| KDR | 9606.ENSP00000263923 | GO Process | GO:0071695 | Anatomical structure maturation |
| KDR | 9606.ENSP00000263923 | GO Process | GO:0071704 | Organic substance metabolic process |
| KDR | 9606.ENSP00000263923 | GO Process | GO:0072089 | Stem cell proliferation |
| KDR | 9606.ENSP00000263923 | GO Process | GO:0072091 | Regulation of stem cell proliferation |
| KDR | 9606.ENSP00000263923 | GO Process | GO:0072359 | Circulatory system development |
| KDR | 9606.ENSP00000263923 | GO Process | GO:0072507 | Divalent inorganic cation homeostasis |
| KDR | 9606.ENSP00000263923 | GO Process | GO:0080090 | Regulation of primary metabolic process |
| KDR | 9606.ENSP00000263923 | GO Process | GO:0090049 | Regulation of cell migration involved in sprouting angiogenesis |
| KDR | 9606.ENSP00000263923 | GO Process | GO:0090050 | Positive regulation of cell migration involved in sprouting angiogenesis |
| KDR | 9606.ENSP00000263923 | GO Process | GO:0090092 | Regulation of transmembrane receptor protein serine/threonine kinase signaling pathway |
| KDR | 9606.ENSP00000263923 | GO Process | GO:0090100 | Positive regulation of transmembrane receptor protein serine/threonine kinase signaling pathway |
| KDR | 9606.ENSP00000263923 | GO Process | GO:0090109 | Regulation of cell-substrate junction assembly |
| KDR | 9606.ENSP00000263923 | GO Process | GO:0090130 | Tissue migration |
| KDR | 9606.ENSP00000263923 | GO Process | GO:0090132 | Epithelium migration |
| KDR | 9606.ENSP00000263923 | GO Process | GO:0090140 | Regulation of mitochondrial fission |
| KDR | 9606.ENSP00000263923 | GO Process | GO:0090141 | Positive regulation of mitochondrial fission |
| KDR | 9606.ENSP00000263923 | GO Process | GO:0090287 | Regulation of cellular response to growth factor stimulus |
| KDR | 9606.ENSP00000263923 | GO Process | GO:0090596 | Sensory organ morphogenesis |
| KDR | 9606.ENSP00000263923 | GO Process | GO:0098771 | Inorganic ion homeostasis |
| KDR | 9606.ENSP00000263923 | GO Process | GO:0150063 | Visual system development |
| KDR | 9606.ENSP00000263923 | GO Process | GO:0150116 | Regulation of cell-substrate junction organization |

| | | | | |
|---|---|---|---|---|
| KDR | 9606.ENSP00000263923 | GO Process | GO:0150117 | Positive regulation of cell-substrate junction organization |
| KDR | 9606.ENSP00000263923 | GO Process | GO:1901214 | Regulation of neuron death |
| KDR | 9606.ENSP00000263923 | GO Process | GO:1901215 | Negative regulation of neuron death |
| KDR | 9606.ENSP00000263923 | GO Process | GO:1901342 | Regulation of vasculature development |
| KDR | 9606.ENSP00000263923 | GO Process | GO:1901532 | Regulation of hematopoietic progenitor cell differentiation |
| KDR | 9606.ENSP00000263923 | GO Process | GO:1901564 | Organonitrogen compound metabolic process |
| KDR | 9606.ENSP00000263923 | GO Process | GO:1901888 | Regulation of cell junction assembly |
| KDR | 9606.ENSP00000263923 | GO Process | GO:1901890 | Positive regulation of cell junction assembly |
| KDR | 9606.ENSP00000263923 | GO Process | GO:1902531 | Regulation of intracellular signal transduction |
| KDR | 9606.ENSP00000263923 | GO Process | GO:1902533 | Positive regulation of intracellular signal transduction |
| KDR | 9606.ENSP00000263923 | GO Process | GO:1903010 | Regulation of bone development |
| KDR | 9606.ENSP00000263923 | GO Process | GO:1904018 | Positive regulation of vasculature development |
| KDR | 9606.ENSP00000263923 | GO Process | GO:1904035 | Regulation of epithelial cell apoptotic process |
| KDR | 9606.ENSP00000263923 | GO Process | GO:1904036 | Negative regulation of epithelial cell apoptotic process |
| KDR | 9606.ENSP00000263923 | GO Process | GO:1904181 | Positive regulation of membrane depolarization |
| KDR | 9606.ENSP00000263923 | GO Process | GO:1904880 | Response to hydrogen sulfide |
| KDR | 9606.ENSP00000263923 | GO Process | GO:1904881 | Cellular response to hydrogen sulfide |
| KDR | 9606.ENSP00000263923 | GO Process | GO:2000026 | Regulation of multicellular organismal development |
| KDR | 9606.ENSP00000263923 | GO Process | GO:2000145 | Regulation of cell motility |
| KDR | 9606.ENSP00000263923 | GO Process | GO:2000147 | Positive regulation of cell motility |
| KDR | 9606.ENSP00000263923 | GO Process | GO:2000351 | Regulation of endothelial cell apoptotic process |
| KDR | 9606.ENSP00000263923 | GO Process | GO:2000352 | Negative regulation of endothelial cell apoptotic process |
| KDR | 9606.ENSP00000263923 | GO Process | GO:2000648 | Positive regulation of stem cell proliferation |
| KDR | 9606.ENSP00000263923 | GO Process | GO:2001026 | Regulation of endothelial cell chemotaxis |
| KDR | 9606.ENSP00000263923 | GO Process | GO:2001028 | Positive regulation of endothelial cell chemotaxis |
| KDR | 9606.ENSP00000263923 | GO Process | GO:2001212 | Regulation of vasculogenesis |
| KDR | 9606.ENSP00000263923 | GO Process | GO:2001214 | Positive regulation of vasculogenesis |
| KDR | 9606.ENSP00000263923 | GO Function | GO:0000166 | Nucleotide binding |
| KDR | 9606.ENSP00000263923 | GO Function | GO:0003824 | Catalytic activity |
| KDR | 9606.ENSP00000263923 | GO Function | GO:0004672 | Protein kinase activity |
| KDR | 9606.ENSP00000263923 | GO Function | GO:0004713 | Protein tyrosine kinase activity |
| KDR | 9606.ENSP00000263923 | GO Function | GO:0004714 | Transmembrane receptor protein tyrosine kinase activity |
| KDR | 9606.ENSP00000263923 | GO Function | GO:0004888 | Transmembrane signaling receptor activity |
| KDR | 9606.ENSP00000263923 | GO Function | GO:0005021 | Vascular endothelial growth factor receptor activity |
| KDR | 9606.ENSP00000263923 | GO Function | GO:0005102 | Signaling receptor binding |

| | | | | |
|---|---|---|---|---|
| KDR | 9606.ENSP00000263923 | GO Function | GO:0005178 | Integrin binding |
| KDR | 9606.ENSP00000263923 | GO Function | GO:0005488 | Binding |
| KDR | 9606.ENSP00000263923 | GO Function | GO:0005515 | Protein binding |
| KDR | 9606.ENSP00000263923 | GO Function | GO:0005524 | ATP binding |
| KDR | 9606.ENSP00000263923 | GO Function | GO:0015026 | Coreceptor activity |
| KDR | 9606.ENSP00000263923 | GO Function | GO:0016301 | Kinase activity |
| KDR | 9606.ENSP00000263923 | GO Function | GO:0016740 | Transferase activity |
| KDR | 9606.ENSP00000263923 | GO Function | GO:0016772 | Transferase activity, transferring phosphorus-containing groups |
| KDR | 9606.ENSP00000263923 | GO Function | GO:0016773 | Phosphotransferase activity, alcohol group as acceptor |
| KDR | 9606.ENSP00000263923 | GO Function | GO:0017076 | Purine nucleotide binding |
| KDR | 9606.ENSP00000263923 | GO Function | GO:0019199 | Transmembrane receptor protein kinase activity |
| KDR | 9606.ENSP00000263923 | GO Function | GO:0019838 | Growth factor binding |
| KDR | 9606.ENSP00000263923 | GO Function | GO:0030554 | Adenyl nucleotide binding |
| KDR | 9606.ENSP00000263923 | GO Function | GO:0031072 | Heat shock protein binding |
| KDR | 9606.ENSP00000263923 | GO Function | GO:0032553 | Ribonucleotide binding |
| KDR | 9606.ENSP00000263923 | GO Function | GO:0032555 | Purine ribonucleotide binding |
| KDR | 9606.ENSP00000263923 | GO Function | GO:0032559 | Adenyl ribonucleotide binding |
| KDR | 9606.ENSP00000263923 | GO Function | GO:0035639 | Purine ribonucleoside triphosphate binding |
| KDR | 9606.ENSP00000263923 | GO Function | GO:0036094 | Small molecule binding |
| KDR | 9606.ENSP00000263923 | GO Function | GO:0038023 | Signaling receptor activity |
| KDR | 9606.ENSP00000263923 | GO Function | GO:0038085 | Vascular endothelial growth factor binding |
| KDR | 9606.ENSP00000263923 | GO Function | GO:0042802 | Identical protein binding |
| KDR | 9606.ENSP00000263923 | GO Function | GO:0043167 | Ion binding |
| KDR | 9606.ENSP00000263923 | GO Function | GO:0043168 | Anion binding |
| KDR | 9606.ENSP00000263923 | GO Function | GO:0044877 | Protein-containing complex binding |
| KDR | 9606.ENSP00000263923 | GO Function | GO:0045296 | Cadherin binding |
| KDR | 9606.ENSP00000263923 | GO Function | GO:0050839 | Cell adhesion molecule binding |
| KDR | 9606.ENSP00000263923 | GO Function | GO:0051879 | Hsp90 protein binding |
| KDR | 9606.ENSP00000263923 | GO Function | GO:0060089 | Molecular transducer activity |
| KDR | 9606.ENSP00000263923 | GO Function | GO:0097159 | Organic cyclic compound binding |
| KDR | 9606.ENSP00000263923 | GO Function | GO:0097367 | Carbohydrate derivative binding |
| KDR | 9606.ENSP00000263923 | GO Function | GO:0140096 | Catalytic activity, acting on a protein |
| KDR | 9606.ENSP00000263923 | GO Function | GO:1901265 | Nucleoside phosphate binding |
| KDR | 9606.ENSP00000263923 | GO Function | GO:1901363 | Heterocyclic compound binding |
| KDR | 9606.ENSP00000263923 | GO Component | GO:0005576 | Extracellular region |

| | | | | |
|---|---|---|---|---|
| KDR | 9606.ENSP00000263923 | GO Component | GO:0005622 | Intracellular anatomical structure |
| KDR | 9606.ENSP00000263923 | GO Component | GO:0005634 | Nucleus |
| KDR | 9606.ENSP00000263923 | GO Component | GO:0005737 | Cytoplasm |
| KDR | 9606.ENSP00000263923 | GO Component | GO:0005768 | Endosome |
| KDR | 9606.ENSP00000263923 | GO Component | GO:0005769 | Early endosome |
| KDR | 9606.ENSP00000263923 | GO Component | GO:0005783 | Endoplasmic reticulum |
| KDR | 9606.ENSP00000263923 | GO Component | GO:0005794 | Golgi apparatus |
| KDR | 9606.ENSP00000263923 | GO Component | GO:0005886 | Plasma membrane |
| KDR | 9606.ENSP00000263923 | GO Component | GO:0005887 | Integral component of plasma membrane |
| KDR | 9606.ENSP00000263923 | GO Component | GO:0009897 | External side of plasma membrane |
| KDR | 9606.ENSP00000263923 | GO Component | GO:0009986 | Cell surface |
| KDR | 9606.ENSP00000263923 | GO Component | GO:0012505 | Endomembrane system |
| KDR | 9606.ENSP00000263923 | GO Component | GO:0016020 | Membrane |
| KDR | 9606.ENSP00000263923 | GO Component | GO:0016021 | Integral component of membrane |
| KDR | 9606.ENSP00000263923 | GO Component | GO:0030054 | Cell junction |
| KDR | 9606.ENSP00000263923 | GO Component | GO:0031224 | Intrinsic component of membrane |
| KDR | 9606.ENSP00000263923 | GO Component | GO:0031226 | Intrinsic component of plasma membrane |
| KDR | 9606.ENSP00000263923 | GO Component | GO:0031410 | Cytoplasmic vesicle |
| KDR | 9606.ENSP00000263923 | GO Component | GO:0031982 | Vesicle |
| KDR | 9606.ENSP00000263923 | GO Component | GO:0032991 | Protein-containing complex |
| KDR | 9606.ENSP00000263923 | GO Component | GO:0043226 | Organelle |
| KDR | 9606.ENSP00000263923 | GO Component | GO:0043227 | Membrane-bounded organelle |
| KDR | 9606.ENSP00000263923 | GO Component | GO:0043229 | Intracellular organelle |
| KDR | 9606.ENSP00000263923 | GO Component | GO:0043231 | Intracellular membrane-bounded organelle |
| KDR | 9606.ENSP00000263923 | GO Component | GO:0043235 | Receptor complex |
| KDR | 9606.ENSP00000263923 | GO Component | GO:0045121 | Membrane raft |
| KDR | 9606.ENSP00000263923 | GO Component | GO:0070161 | Anchoring junction |
| KDR | 9606.ENSP00000263923 | GO Component | GO:0071944 | Cell periphery |
| KDR | 9606.ENSP00000263923 | GO Component | GO:0097443 | Sorting endosome |
| KDR | 9606.ENSP00000263923 | GO Component | GO:0097708 | Intracellular vesicle |
| KDR | 9606.ENSP00000263923 | GO Component | GO:0098552 | Side of membrane |
| KDR | 9606.ENSP00000263923 | GO Component | GO:0098857 | Membrane microdomain |
| KDR | 9606.ENSP00000263923 | GO Component | GO:0110165 | Cellular anatomical entity |
| KDR | 9606.ENSP00000263923 | STRING clusters | CL:17325 | Mixed, incl. Constitutive Signaling by Aberrant PI3K in Cancer, and SH2 domain superfamily |

| | | | | |
|---|---|---|---|---|
| KDR | 9606.ENSP00000263923 | STRING clusters | CL:17326 | Mixed, incl. Constitutive Signaling by Aberrant PI3K in Cancer, and SH2 domain superfamily |
| KDR | 9606.ENSP00000263923 | STRING clusters | CL:17327 | Mixed, incl. Constitutive Signaling by Aberrant PI3K in Cancer, and Tyrosine-protein kinase, catalytic domain |
| KDR | 9606.ENSP00000263923 | STRING clusters | CL:17328 | Mixed, incl. Constitutive Signaling by Aberrant PI3K in Cancer, and FCERI mediated Ca+2 mobilization |
| KDR | 9606.ENSP00000263923 | STRING clusters | CL:17329 | Constitutive Signaling by Aberrant PI3K in Cancer, and VEGF ligand-receptor interactions |
| KDR | 9606.ENSP00000263923 | STRING clusters | CL:17330 | Constitutive Signaling by Aberrant PI3K in Cancer, and VEGF ligand-receptor interactions |
| KDR | 9606.ENSP00000263923 | STRING clusters | CL:17331 | Mixed, incl. FGFR2 ligand binding and activation, and PDGF/VEGF domain |
| KDR | 9606.ENSP00000263923 | STRING clusters | CL:17381 | PDGF/VEGF domain, and Vascular endothelial growth factor receptor activity |
| KDR | 9606.ENSP00000263923 | STRING clusters | CL:17382 | VEGF ligand-receptor interactions, and Tie signaling pathway |
| KDR | 9606.ENSP00000263923 | KEGG | hsa01521 | EGFR tyrosine kinase inhibitor resistance |
| KDR | 9606.ENSP00000263923 | KEGG | hsa04010 | MAPK signaling pathway |
| KDR | 9606.ENSP00000263923 | KEGG | hsa04014 | Ras signaling pathway |
| KDR | 9606.ENSP00000263923 | KEGG | hsa04015 | Rap1 signaling pathway |
| KDR | 9606.ENSP00000263923 | KEGG | hsa04151 | PI3K-Akt signaling pathway |
| KDR | 9606.ENSP00000263923 | KEGG | hsa04370 | VEGF signaling pathway |
| KDR | 9606.ENSP00000263923 | KEGG | hsa04510 | Focal adhesion |
| KDR | 9606.ENSP00000263923 | KEGG | hsa05205 | Proteoglycans in cancer |
| KDR | 9606.ENSP00000263923 | KEGG | hsa05418 | Fluid shear stress and atherosclerosis |
| KDR | 9606.ENSP00000263923 | Reactome | HSA-1474244 | Extracellular matrix organization |
| KDR | 9606.ENSP00000263923 | Reactome | HSA-162582 | Signal Transduction |
| KDR | 9606.ENSP00000263923 | Reactome | HSA-1643685 | Disease |
| KDR | 9606.ENSP00000263923 | Reactome | HSA-194138 | Signaling by VEGF |
| KDR | 9606.ENSP00000263923 | Reactome | HSA-194306 | Neurophilin interactions with VEGF and VEGFR |
| KDR | 9606.ENSP00000263923 | Reactome | HSA-194313 | VEGF ligand-receptor interactions |
| KDR | 9606.ENSP00000263923 | Reactome | HSA-195399 | VEGF binds to VEGFR leading to receptor dimerization |
| KDR | 9606.ENSP00000263923 | Reactome | HSA-216083 | Integrin cell surface interactions |
| KDR | 9606.ENSP00000263923 | Reactome | HSA-4420097 | VEGFA-VEGFR2 Pathway |
| KDR | 9606.ENSP00000263923 | Reactome | HSA-5218921 | VEGFR2 mediated cell proliferation |
| KDR | 9606.ENSP00000263923 | Reactome | HSA-5663202 | Diseases of signal transduction by growth factor receptors and second messengers |

| | | | | |
|---|---|---|---|---|
| KDR | 9606.ENSP00000263923 | Reactome | HSA-9006934 | Signaling by Receptor Tyrosine Kinases |
| KDR | 9606.ENSP00000263923 | Reactome | HSA-9671555 | Signaling by PDGFR in disease |
| KDR | 9606.ENSP00000263923 | Reactome | HSA-9673768 | Signaling by membrane-tethered fusions of PDGFRA or PDGFRB |
| KDR | 9606.ENSP00000263923 | WikiPathways | WP1539 | Angiogenesis |
| KDR | 9606.ENSP00000263923 | WikiPathways | WP2406 | Cardiac progenitor differentiation |
| KDR | 9606.ENSP00000263923 | WikiPathways | WP306 | Focal adhesion |
| KDR | 9606.ENSP00000263923 | WikiPathways | WP3888 | VEGFA-VEGFR2 signaling |
| KDR | 9606.ENSP00000263923 | WikiPathways | WP3932 | Focal adhesion: PI3K-Akt-mTOR-signaling pathway |
| KDR | 9606.ENSP00000263923 | WikiPathways | WP3943 | Robo4 and VEGF signaling pathways crosstalk |
| KDR | 9606.ENSP00000263923 | WikiPathways | WP4018 | Clear cell renal cell carcinoma pathways |
| KDR | 9606.ENSP00000263923 | WikiPathways | WP4172 | PI3K-Akt signaling pathway |
| KDR | 9606.ENSP00000263923 | WikiPathways | WP4223 | Ras signaling |
| KDR | 9606.ENSP00000263923 | WikiPathways | WP4300 | Extracellular vesicles in the crosstalk of cardiac cells |
| KDR | 9606.ENSP00000263923 | WikiPathways | WP4331 | Neovascularisation processes |
| KDR | 9606.ENSP00000263923 | WikiPathways | WP4540 | Hippo signaling regulation pathways |
| KDR | 9606.ENSP00000263923 | WikiPathways | WP4541 | Hippo-Merlin signaling dysregulation |
| KDR | 9606.ENSP00000263923 | WikiPathways | WP4685 | Melanoma |
| KDR | 9606.ENSP00000263923 | WikiPathways | WP4747 | Netrin-UNC5B signaling pathway |
| KDR | 9606.ENSP00000263923 | WikiPathways | WP4806 | EGFR tyrosine kinase inhibitor resistance |
| KDR | 9606.ENSP00000263923 | WikiPathways | WP4823 | Genes controlling nephrogenesis |
| KDR | 9606.ENSP00000263923 | WikiPathways | WP4950 | 16p11.2 distal deletion syndrome |
| KDR | 9606.ENSP00000263923 | WikiPathways | WP5055 | Burn wound healing |
| KDR | 9606.ENSP00000263923 | WikiPathways | WP5065 | SARS-CoV-2 altering angiogenesis via NRP1 |
| KDR | 9606.ENSP00000263923 | WikiPathways | WP5087 | Malignant pleural mesothelioma |
| KDR | 9606.ENSP00000263923 | WikiPathways | WP5144 | NRP1-triggered signaling pathways in pancreatic cancer |
| KDR | 9606.ENSP00000263923 | WikiPathways | WP5236 | Markers of kidney cell lineage |
| KDR | 9606.ENSP00000263923 | WikiPathways | WP5284 | Cell interactions of the pancreatic cancer microenvironment |
| KDR | 9606.ENSP00000263923 | WikiPathways | WP5316 | Primary ovarian insufficiency |
| KDR | 9606.ENSP00000263923 | WikiPathways | WP5322 | CKAP4 signaling pathway map |
| KDR | 9606.ENSP00000263923 | DISEASES | DOID:0050117 | Disease by infectious agent |
| KDR | 9606.ENSP00000263923 | DISEASES | DOID:0050686 | Organ system cancer |
| KDR | 9606.ENSP00000263923 | DISEASES | DOID:0050687 | Cell type cancer |
| KDR | 9606.ENSP00000263923 | DISEASES | DOID:0080374 | Gastroesophageal cancer |
| KDR | 9606.ENSP00000263923 | DISEASES | DOID:0080375 | Gastroesophageal adenocarcinoma |
| KDR | 9606.ENSP00000263923 | DISEASES | DOID:1398 | Parasitic infectious disease |

| | | | | |
|---|---|---|---|---|
| KDR | 9606.ENSP00000263923 | DISEASES | DOID:14566 | Disease of cellular proliferation |
| KDR | 9606.ENSP00000263923 | DISEASES | DOID:162 | Cancer |
| KDR | 9606.ENSP00000263923 | DISEASES | DOID:299 | Adenocarcinoma |
| KDR | 9606.ENSP00000263923 | DISEASES | DOID:305 | Carcinoma |
| KDR | 9606.ENSP00000263923 | DISEASES | DOID:3119 | Gastrointestinal system cancer |
| KDR | 9606.ENSP00000263923 | DISEASES | DOID:4 | Disease |
| KDR | 9606.ENSP00000263923 | DISEASES | DOID:4110 | Parasitic ectoparasitic infectious disease |
| KDR | 9606.ENSP00000263923 | DISEASES | DOID:4944 | Gastroesophageal junction adenocarcinoma |
| KDR | 9606.ENSP00000263923 | DISEASES | DOID:5501 | Pediculus humanus capitis infestation |
| KDR | 9606.ENSP00000263923 | DISEASES | DOID:5502 | Lice infestation |
| KDR | 9606.ENSP00000263923 | DISEASES | DOID:7 | Disease of anatomical entity |
| KDR | 9606.ENSP00000263923 | DISEASES | DOID:77 | Gastrointestinal system disease |
| KDR | 9606.ENSP00000263923 | TISSUES | BTO:0000000 | Tissues, cell types and enzyme sources |
| KDR | 9606.ENSP00000263923 | TISSUES | BTO:0000042 | Animal |
| KDR | 9606.ENSP00000263923 | TISSUES | BTO:0000081 | Reproductive system |
| KDR | 9606.ENSP00000263923 | TISSUES | BTO:0000083 | Female reproductive system |
| KDR | 9606.ENSP00000263923 | TISSUES | BTO:0000088 | Cardiovascular system |
| KDR | 9606.ENSP00000263923 | TISSUES | BTO:0000089 | Blood |
| KDR | 9606.ENSP00000263923 | TISSUES | BTO:0000135 | Aorta |
| KDR | 9606.ENSP00000263923 | TISSUES | BTO:0000174 | Embryonic structure |
| KDR | 9606.ENSP00000263923 | TISSUES | BTO:0000202 | Sense organ |
| KDR | 9606.ENSP00000263923 | TISSUES | BTO:0000251 | Chorioallantois |
| KDR | 9606.ENSP00000263923 | TISSUES | BTO:0000282 | Head |
| KDR | 9606.ENSP00000263923 | TISSUES | BTO:0000284 | Organism form |
| KDR | 9606.ENSP00000263923 | TISSUES | BTO:0000379 | Embryo |
| KDR | 9606.ENSP00000263923 | TISSUES | BTO:0000381 | Embryonic blood |
| KDR | 9606.ENSP00000263923 | TISSUES | BTO:0000393 | Endothelium |
| KDR | 9606.ENSP00000263923 | TISSUES | BTO:0000416 | Epithelium |
| KDR | 9606.ENSP00000263923 | TISSUES | BTO:0000439 | Eye |
| KDR | 9606.ENSP00000263923 | TISSUES | BTO:0000449 | Fetus |
| KDR | 9606.ENSP00000263923 | TISSUES | BTO:0000473 | Fetal membrane |
| KDR | 9606.ENSP00000263923 | TISSUES | BTO:0000553 | Peripheral blood |
| KDR | 9606.ENSP00000263923 | TISSUES | BTO:0000556 | Germ layer |
| KDR | 9606.ENSP00000263923 | TISSUES | BTO:0000562 | Heart |
| KDR | 9606.ENSP00000263923 | TISSUES | BTO:0000570 | Hematopoietic system |

| | | | | |
|---|---|---|---|---|
| KDR | 9606.ENSP00000263923 | TISSUES | BTO:0000573 | Artery |
| KDR | 9606.ENSP00000263923 | TISSUES | BTO:0000634 | Integument |
| KDR | 9606.ENSP00000263923 | TISSUES | BTO:0000720 | Plant form |
| KDR | 9606.ENSP00000263923 | TISSUES | BTO:0000766 | Blood vessel endothelium |
| KDR | 9606.ENSP00000263923 | TISSUES | BTO:0000839 | Mesoderm |
| KDR | 9606.ENSP00000263923 | TISSUES | BTO:0001078 | Placenta |
| KDR | 9606.ENSP00000263923 | TISSUES | BTO:0001085 | Vascular system |
| KDR | 9606.ENSP00000263923 | TISSUES | BTO:0001102 | Blood vessel |
| KDR | 9606.ENSP00000263923 | TISSUES | BTO:0001120 | Epithelial cell line |
| KDR | 9606.ENSP00000263923 | TISSUES | BTO:0001175 | Retina |
| KDR | 9606.ENSP00000263923 | TISSUES | BTO:0001176 | Endothelial cell |
| KDR | 9606.ENSP00000263923 | TISSUES | BTO:0001228 | Seedling |
| KDR | 9606.ENSP00000263923 | TISSUES | BTO:0001296 | Sprout |
| KDR | 9606.ENSP00000263923 | TISSUES | BTO:0001393 | Mesenchyme |
| KDR | 9606.ENSP00000263923 | TISSUES | BTO:0001415 | Umbilical cord |
| KDR | 9606.ENSP00000263923 | TISSUES | BTO:0001461 | Whole plant |
| KDR | 9606.ENSP00000263923 | TISSUES | BTO:0001481 | Plant |
| KDR | 9606.ENSP00000263923 | TISSUES | BTO:0001489 | Whole body |
| KDR | 9606.ENSP00000263923 | TISSUES | BTO:0001509 | Umbilical vein |
| KDR | 9606.ENSP00000263923 | TISSUES | BTO:0001519 | Endothelial cell line |
| KDR | 9606.ENSP00000263923 | TISSUES | BTO:0001520 | Umbilical vein endothelial cell line |
| KDR | 9606.ENSP00000263923 | TISSUES | BTO:0001853 | Vascular endothelium |
| KDR | 9606.ENSP00000263923 | TISSUES | BTO:0001854 | Vascular endothelial cell |
| KDR | 9606.ENSP00000263923 | TISSUES | BTO:0001949 | HUVEC cell |
| KDR | 9606.ENSP00000263923 | TISSUES | BTO:0002045 | Capillary |
| KDR | 9606.ENSP00000263923 | TISSUES | BTO:0003091 | Urogenital system |
| KDR | 9606.ENSP00000263923 | TISSUES | BTO:0003099 | Internal female genital organ |
| KDR | 9606.ENSP00000263923 | TISSUES | BTO:0003122 | Microvascular endothelium |
| KDR | 9606.ENSP00000263923 | TISSUES | BTO:0003123 | Microvascular endothelial cell |
| KDR | 9606.ENSP00000263923 | TISSUES | BTO:0003494 | Peripheral blood cell |
| KDR | 9606.ENSP00000263923 | TISSUES | BTO:0003718 | Vasculature |
| KDR | 9606.ENSP00000263923 | TISSUES | BTO:0004092 | Endogenous progenitor cell |
| KDR | 9606.ENSP00000263923 | TISSUES | BTO:0004093 | Endothelial progenitor cell |
| KDR | 9606.ENSP00000263923 | TISSUES | BTO:0004395 | Microvessel |
| KDR | 9606.ENSP00000263923 | TISSUES | BTO:0004673 | Dorsal aorta |

| | | | | |
|---|---|---|---|---|
| KDR | 9606.ENSP00000263923 | TISSUES | BTO:0004985 | Angioblast |
| KDR | 9606.ENSP00000263923 | TISSUES | BTO:0005001 | Retinal microvascular endothelial cell |
| KDR | 9606.ENSP00000263923 | TISSUES | BTO:0005587 | Stalk cell |
| KDR | 9606.ENSP00000263923 | COMPARTMENTS | GOCC:0005576 | Extracellular region |
| KDR | 9606.ENSP00000263923 | COMPARTMENTS | GOCC:0005622 | Intracellular |
| KDR | 9606.ENSP00000263923 | COMPARTMENTS | GOCC:0005634 | Nucleus |
| KDR | 9606.ENSP00000263923 | COMPARTMENTS | GOCC:0005737 | Cytoplasm |
| KDR | 9606.ENSP00000263923 | COMPARTMENTS | GOCC:0005768 | Endosome |
| KDR | 9606.ENSP00000263923 | COMPARTMENTS | GOCC:0005769 | Early endosome |
| KDR | 9606.ENSP00000263923 | COMPARTMENTS | GOCC:0005783 | Endoplasmic reticulum |
| KDR | 9606.ENSP00000263923 | COMPARTMENTS | GOCC:0005794 | Golgi apparatus |
| KDR | 9606.ENSP00000263923 | COMPARTMENTS | GOCC:0005886 | Plasma membrane |
| KDR | 9606.ENSP00000263923 | COMPARTMENTS | GOCC:0005887 | Integral component of plasma membrane |
| KDR | 9606.ENSP00000263923 | COMPARTMENTS | GOCC:0012505 | Endomembrane system |
| KDR | 9606.ENSP00000263923 | COMPARTMENTS | GOCC:0016020 | Membrane |
| KDR | 9606.ENSP00000263923 | COMPARTMENTS | GOCC:0016021 | Integral component of membrane |
| KDR | 9606.ENSP00000263923 | COMPARTMENTS | GOCC:0030054 | Cell junction |
| KDR | 9606.ENSP00000263923 | COMPARTMENTS | GOCC:0031224 | Intrinsic component of membrane |
| KDR | 9606.ENSP00000263923 | COMPARTMENTS | GOCC:0031226 | Intrinsic component of plasma membrane |
| KDR | 9606.ENSP00000263923 | COMPARTMENTS | GOCC:0031410 | Cytoplasmic vesicle |
| KDR | 9606.ENSP00000263923 | COMPARTMENTS | GOCC:0031982 | Vesicle |
| KDR | 9606.ENSP00000263923 | COMPARTMENTS | GOCC:0032991 | Protein-containing complex |
| KDR | 9606.ENSP00000263923 | COMPARTMENTS | GOCC:0036454 | Growth factor complex |
| KDR | 9606.ENSP00000263923 | COMPARTMENTS | GOCC:0043226 | Organelle |
| KDR | 9606.ENSP00000263923 | COMPARTMENTS | GOCC:0043227 | Membrane-bounded organelle |
| KDR | 9606.ENSP00000263923 | COMPARTMENTS | GOCC:0043229 | Intracellular organelle |
| KDR | 9606.ENSP00000263923 | COMPARTMENTS | GOCC:0043231 | Intracellular membrane-bounded organelle |
| KDR | 9606.ENSP00000263923 | COMPARTMENTS | GOCC:0044232 | Organelle membrane contact site |
| KDR | 9606.ENSP00000263923 | COMPARTMENTS | GOCC:0045121 | Membrane raft |
| KDR | 9606.ENSP00000263923 | COMPARTMENTS | GOCC:0071944 | Cell periphery |
| KDR | 9606.ENSP00000263923 | COMPARTMENTS | GOCC:0097443 | Sorting endosome |
| KDR | 9606.ENSP00000263923 | COMPARTMENTS | GOCC:0097708 | Intracellular vesicle |
| KDR | 9606.ENSP00000263923 | COMPARTMENTS | GOCC:0098857 | Membrane microdomain |
| KDR | 9606.ENSP00000263923 | COMPARTMENTS | GOCC:0110165 | Cellular anatomical entity |
| KDR | 9606.ENSP00000263923 | COMPARTMENTS | GOCC:0140268 | Endoplasmic reticulum-plasma membrane contact site |

| | | | | |
|---|---|---|---|---|
| KDR | 9606.ENSP00000263923 | COMPARTMENTS | GOCC:1990150 | VEGF-A complex |
| KDR | 9606.ENSP00000263923 | Monarch | EFO:0003892 | Pulmonary function measurement |
| KDR | 9606.ENSP00000263923 | Monarch | EFO:0004312 | Vital capacity |
| KDR | 9606.ENSP00000263923 | Monarch | EFO:0004747 | Protein measurement |
| KDR | 9606.ENSP00000263923 | Monarch | EFO:0007937 | Blood protein measurement |
| KDR | 9606.ENSP00000263923 | Monarch | EFO:0008314 | Vascular endothelial growth factor receptor 2 measurement |
| KDR | 9606.ENSP00000263923 | Monarch | EFO:0008376 | Mosquito bite measurement |
| KDR | 9606.ENSP00000263923 | Monarch | EFO:0008378 | Mosquito bite reaction size measurement |
| KDR | 9606.ENSP00000263923 | Monarch | HP:0000118 | Phenotypic abnormality |
| KDR | 9606.ENSP00000263923 | Monarch | HP:0001028 | Hemangioma |
| KDR | 9606.ENSP00000263923 | Monarch | HP:0001626 | Abnormality of the cardiovascular system |
| KDR | 9606.ENSP00000263923 | Monarch | HP:0002597 | Abnormality of the vasculature |
| KDR | 9606.ENSP00000263923 | Monarch | HP:0002664 | Neoplasm |
| KDR | 9606.ENSP00000263923 | Monarch | HP:0005306 | Capillary hemangioma |
| KDR | 9606.ENSP00000263923 | Monarch | HP:0011793 | Neoplasm by anatomical site |
| KDR | 9606.ENSP00000263923 | Monarch | HP:0100742 | Vascular neoplasm |
| KDR | 9606.ENSP00000263923 | UniProt Keywords | KW-0025 | Alternative splicing |
| KDR | 9606.ENSP00000263923 | UniProt Keywords | KW-0037 | Angiogenesis |
| KDR | 9606.ENSP00000263923 | UniProt Keywords | KW-0067 | ATP-binding |
| KDR | 9606.ENSP00000263923 | UniProt Keywords | KW-0217 | Developmental protein |
| KDR | 9606.ENSP00000263923 | UniProt Keywords | KW-0221 | Differentiation |
| KDR | 9606.ENSP00000263923 | UniProt Keywords | KW-0256 | Endoplasmic reticulum |
| KDR | 9606.ENSP00000263923 | UniProt Keywords | KW-0325 | Glycoprotein |
| KDR | 9606.ENSP00000263923 | UniProt Keywords | KW-0393 | Immunoglobulin domain |
| KDR | 9606.ENSP00000263923 | UniProt Keywords | KW-0418 | Kinase |
| KDR | 9606.ENSP00000263923 | UniProt Keywords | KW-0472 | Membrane |
| KDR | 9606.ENSP00000263923 | UniProt Keywords | KW-0539 | Nucleus |
| KDR | 9606.ENSP00000263923 | UniProt Keywords | KW-0547 | Nucleotide-binding |
| KDR | 9606.ENSP00000263923 | UniProt Keywords | KW-0597 | Phosphoprotein |
| KDR | 9606.ENSP00000263923 | UniProt Keywords | KW-0675 | Receptor |
| KDR | 9606.ENSP00000263923 | UniProt Keywords | KW-0677 | Repeat |
| KDR | 9606.ENSP00000263923 | UniProt Keywords | KW-0732 | Signal |
| KDR | 9606.ENSP00000263923 | UniProt Keywords | KW-0808 | Transferase |
| KDR | 9606.ENSP00000263923 | UniProt Keywords | KW-0812 | Transmembrane |
| KDR | 9606.ENSP00000263923 | UniProt Keywords | KW-0829 | Tyrosine-protein kinase |

| | | | | |
|---|---|---|---|---|
| KDR | 9606.ENSP00000263923 | UniProt Keywords | KW-0832 | Ubl conjugation |
| KDR | 9606.ENSP00000263923 | UniProt Keywords | KW-0945 | Host-virus interaction |
| KDR | 9606.ENSP00000263923 | UniProt Keywords | KW-0963 | Cytoplasm |
| KDR | 9606.ENSP00000263923 | UniProt Keywords | KW-0964 | Secreted |
| KDR | 9606.ENSP00000263923 | UniProt Keywords | KW-0965 | Cell junction |
| KDR | 9606.ENSP00000263923 | UniProt Keywords | KW-0967 | Endosome |
| KDR | 9606.ENSP00000263923 | UniProt Keywords | KW-0968 | Cytoplasmic vesicle |
| KDR | 9606.ENSP00000263923 | UniProt Keywords | KW-1003 | Cell membrane |
| KDR | 9606.ENSP00000263923 | UniProt Keywords | KW-1015 | Disulfide bond |
| KDR | 9606.ENSP00000263923 | UniProt Keywords | KW-1133 | Transmembrane helix |
| KDR | 9606.ENSP00000263923 | InterPro | IPR000719 | Protein kinase domain |
| KDR | 9606.ENSP00000263923 | InterPro | IPR001245 | Serine-threonine/tyrosine-protein kinase, catalytic domain |
| KDR | 9606.ENSP00000263923 | InterPro | IPR001824 | Tyrosine-protein kinase, receptor class III, conserved site |
| KDR | 9606.ENSP00000263923 | InterPro | IPR003598 | Immunoglobulin subtype 2 |
| KDR | 9606.ENSP00000263923 | InterPro | IPR003599 | Immunoglobulin subtype |
| KDR | 9606.ENSP00000263923 | InterPro | IPR007110 | Immunoglobulin-like domain |
| KDR | 9606.ENSP00000263923 | InterPro | IPR008266 | Tyrosine-protein kinase, active site |
| KDR | 9606.ENSP00000263923 | InterPro | IPR009136 | Vascular endothelial growth factor receptor 2 (VEGFR2) |
| KDR | 9606.ENSP00000263923 | InterPro | IPR011009 | Protein kinase-like domain superfamily |
| KDR | 9606.ENSP00000263923 | InterPro | IPR013098 | Immunoglobulin I-set |
| KDR | 9606.ENSP00000263923 | InterPro | IPR013783 | Immunoglobulin-like fold |
| KDR | 9606.ENSP00000263923 | InterPro | IPR017441 | Protein kinase, ATP binding site |
| KDR | 9606.ENSP00000263923 | InterPro | IPR020635 | Tyrosine-protein kinase, catalytic domain |
| KDR | 9606.ENSP00000263923 | InterPro | IPR036179 | Immunoglobulin-like domain superfamily |
| KDR | 9606.ENSP00000263923 | InterPro | IPR041348 | VEGFR-2, transmembrane domain |
| KDR | 9606.ENSP00000263923 | SMART | SM00219 | Tyrosine kinase, catalytic domain |
| KDR | 9606.ENSP00000263923 | SMART | SM00408 | Immunoglobulin C-2 Type |
| KDR | 9606.ENSP00000263923 | SMART | SM00409 | Immunoglobulin |
| KDR | 9606.ENSP00000263923 | SMART | SM00410 | Immunoglobulin like |
| LPAR4 | 9606.ENSP00000408205 | GO Process | GO:0007154 | Cell communication |
| LPAR4 | 9606.ENSP00000408205 | GO Process | GO:0007165 | Signal transduction |
| LPAR4 | 9606.ENSP00000408205 | GO Process | GO:0007186 | G protein-coupled receptor signaling pathway |
| LPAR4 | 9606.ENSP00000408205 | GO Process | GO:0007200 | Phospholipase C-activating G protein-coupled receptor signaling pathway |
| LPAR4 | 9606.ENSP00000408205 | GO Process | GO:0007204 | Positive regulation of cytosolic calcium ion concentration |
| LPAR4 | 9606.ENSP00000408205 | GO Process | GO:0009966 | Regulation of signal transduction |

| | | | | |
|---|---|---|---|---|
| LPAR4 | 9606.ENSP00000408205 | GO Process | GO:0009967 | Positive regulation of signal transduction |
| LPAR4 | 9606.ENSP00000408205 | GO Process | GO:0009987 | Cellular process |
| LPAR4 | 9606.ENSP00000408205 | GO Process | GO:0010646 | Regulation of cell communication |
| LPAR4 | 9606.ENSP00000408205 | GO Process | GO:0010647 | Positive regulation of cell communication |
| LPAR4 | 9606.ENSP00000408205 | GO Process | GO:0023051 | Regulation of signaling |
| LPAR4 | 9606.ENSP00000408205 | GO Process | GO:0023052 | Signaling |
| LPAR4 | 9606.ENSP00000408205 | GO Process | GO:0023056 | Positive regulation of signaling |
| LPAR4 | 9606.ENSP00000408205 | GO Process | GO:0035023 | Regulation of Rho protein signal transduction |
| LPAR4 | 9606.ENSP00000408205 | GO Process | GO:0035025 | Positive regulation of Rho protein signal transduction |
| LPAR4 | 9606.ENSP00000408205 | GO Process | GO:0046578 | Regulation of Ras protein signal transduction |
| LPAR4 | 9606.ENSP00000408205 | GO Process | GO:0046579 | Positive regulation of Ras protein signal transduction |
| LPAR4 | 9606.ENSP00000408205 | GO Process | GO:0048518 | Positive regulation of biological process |
| LPAR4 | 9606.ENSP00000408205 | GO Process | GO:0048522 | Positive regulation of cellular process |
| LPAR4 | 9606.ENSP00000408205 | GO Process | GO:0048583 | Regulation of response to stimulus |
| LPAR4 | 9606.ENSP00000408205 | GO Process | GO:0048584 | Positive regulation of response to stimulus |
| LPAR4 | 9606.ENSP00000408205 | GO Process | GO:0050789 | Regulation of biological process |
| LPAR4 | 9606.ENSP00000408205 | GO Process | GO:0050794 | Regulation of cellular process |
| LPAR4 | 9606.ENSP00000408205 | GO Process | GO:0050896 | Response to stimulus |
| LPAR4 | 9606.ENSP00000408205 | GO Process | GO:0051056 | Regulation of small GTPase mediated signal transduction |
| LPAR4 | 9606.ENSP00000408205 | GO Process | GO:0051057 | Positive regulation of small GTPase mediated signal transduction |
| LPAR4 | 9606.ENSP00000408205 | GO Process | GO:0051482 | Positive regulation of cytosolic calcium ion concentration involved in phospholipase C-activating G protein-coupled signaling pathway |
| LPAR4 | 9606.ENSP00000408205 | GO Process | GO:0051716 | Cellular response to stimulus |
| LPAR4 | 9606.ENSP00000408205 | GO Process | GO:0065007 | Biological regulation |
| LPAR4 | 9606.ENSP00000408205 | GO Process | GO:0065008 | Regulation of biological quality |
| LPAR4 | 9606.ENSP00000408205 | GO Process | GO:1902531 | Regulation of intracellular signal transduction |
| LPAR4 | 9606.ENSP00000408205 | GO Process | GO:1902533 | Positive regulation of intracellular signal transduction |
| LPAR4 | 9606.ENSP00000408205 | GO Function | GO:0004888 | Transmembrane signaling receptor activity |
| LPAR4 | 9606.ENSP00000408205 | GO Function | GO:0004930 | G protein-coupled receptor activity |
| LPAR4 | 9606.ENSP00000408205 | GO Function | GO:0005488 | Binding |
| LPAR4 | 9606.ENSP00000408205 | GO Function | GO:0005543 | Phospholipid binding |
| LPAR4 | 9606.ENSP00000408205 | GO Function | GO:0008289 | Lipid binding |
| LPAR4 | 9606.ENSP00000408205 | GO Function | GO:0035727 | Lysophosphatidic acid binding |
| LPAR4 | 9606.ENSP00000408205 | GO Function | GO:0038023 | Signaling receptor activity |
| LPAR4 | 9606.ENSP00000408205 | GO Function | GO:0043167 | Ion binding |

| | | | | |
|---|---|---|---|---|
| LPAR4 | 9606.ENSP00000408205 | GO Function | GO:0043168 | Anion binding |
| LPAR4 | 9606.ENSP00000408205 | GO Function | GO:0045125 | Bioactive lipid receptor activity |
| LPAR4 | 9606.ENSP00000408205 | GO Function | GO:0060089 | Molecular transducer activity |
| LPAR4 | 9606.ENSP00000408205 | GO Function | GO:0070915 | Lysophosphatidic acid receptor activity |
| LPAR4 | 9606.ENSP00000408205 | GO Function | GO:0097367 | Carbohydrate derivative binding |
| LPAR4 | 9606.ENSP00000408205 | GO Component | GO:0005622 | Intracellular anatomical structure |
| LPAR4 | 9606.ENSP00000408205 | GO Component | GO:0005634 | Nucleus |
| LPAR4 | 9606.ENSP00000408205 | GO Component | GO:0005654 | Nucleoplasm |
| LPAR4 | 9606.ENSP00000408205 | GO Component | GO:0005886 | Plasma membrane |
| LPAR4 | 9606.ENSP00000408205 | GO Component | GO:0005887 | Integral component of plasma membrane |
| LPAR4 | 9606.ENSP00000408205 | GO Component | GO:0016020 | Membrane |
| LPAR4 | 9606.ENSP00000408205 | GO Component | GO:0016021 | Integral component of membrane |
| LPAR4 | 9606.ENSP00000408205 | GO Component | GO:0016604 | Nuclear body |
| LPAR4 | 9606.ENSP00000408205 | GO Component | GO:0031224 | Intrinsic component of membrane |
| LPAR4 | 9606.ENSP00000408205 | GO Component | GO:0031226 | Intrinsic component of plasma membrane |
| LPAR4 | 9606.ENSP00000408205 | GO Component | GO:0031974 | Membrane-enclosed lumen |
| LPAR4 | 9606.ENSP00000408205 | GO Component | GO:0031981 | Nuclear lumen |
| LPAR4 | 9606.ENSP00000408205 | GO Component | GO:0043226 | Organelle |
| LPAR4 | 9606.ENSP00000408205 | GO Component | GO:0043227 | Membrane-bounded organelle |
| LPAR4 | 9606.ENSP00000408205 | GO Component | GO:0043229 | Intracellular organelle |
| LPAR4 | 9606.ENSP00000408205 | GO Component | GO:0043231 | Intracellular membrane-bounded organelle |
| LPAR4 | 9606.ENSP00000408205 | GO Component | GO:0043233 | Organelle lumen |
| LPAR4 | 9606.ENSP00000408205 | GO Component | GO:0070013 | Intracellular organelle lumen |
| LPAR4 | 9606.ENSP00000408205 | GO Component | GO:0071944 | Cell periphery |
| LPAR4 | 9606.ENSP00000408205 | GO Component | GO:0110165 | Cellular anatomical entity |
| LPAR4 | 9606.ENSP00000408205 | STRING clusters | CL:24306 | Mixed, incl. Photoreceptor outer segment, and Calcium regulation in cardiac cells |
| LPAR4 | 9606.ENSP00000408205 | STRING clusters | CL:24307 | Mixed, incl. Heterotrimeric G-protein complex, and Signal transduction inhibitor |
| LPAR4 | 9606.ENSP00000408205 | STRING clusters | CL:24458 | Bioactive lipid receptor activity, and Phospholipase C-activating G protein-coupled acetylcholine receptor signaling pathway |
| LPAR4 | 9606.ENSP00000408205 | STRING clusters | CL:24461 | Lysophosphatidic acid receptor activity |
| LPAR4 | 9606.ENSP00000408205 | KEGG | hsa04015 | Rap1 signaling pathway |
| LPAR4 | 9606.ENSP00000408205 | KEGG | hsa04072 | Phospholipase D signaling pathway |
| LPAR4 | 9606.ENSP00000408205 | KEGG | hsa04080 | Neuroactive ligand-receptor interaction |

| | | | | |
|---|---|---|---|---|
| LPAR4 | 9606.ENSP00000408205 | KEGG | hsa04151 | PI3K-Akt signaling pathway |
| LPAR4 | 9606.ENSP00000408205 | KEGG | hsa04810 | Regulation of actin cytoskeleton |
| LPAR4 | 9606.ENSP00000408205 | KEGG | hsa05130 | Pathogenic Escherichia coli infection |
| LPAR4 | 9606.ENSP00000408205 | KEGG | hsa05200 | Pathways in cancer |
| LPAR4 | 9606.ENSP00000408205 | Reactome | HSA-162582 | Signal Transduction |
| LPAR4 | 9606.ENSP00000408205 | Reactome | HSA-372790 | Signaling by GPCR |
| LPAR4 | 9606.ENSP00000408205 | Reactome | HSA-373076 | Class A/1 (Rhodopsin-like receptors) |
| LPAR4 | 9606.ENSP00000408205 | Reactome | HSA-388396 | GPCR downstream signalling |
| LPAR4 | 9606.ENSP00000408205 | Reactome | HSA-416476 | G alpha (q) signalling events |
| LPAR4 | 9606.ENSP00000408205 | Reactome | HSA-417957 | P2Y receptors |
| LPAR4 | 9606.ENSP00000408205 | Reactome | HSA-418038 | Nucleotide-like (purinergic) receptors |
| LPAR4 | 9606.ENSP00000408205 | Reactome | HSA-500792 | GPCR ligand binding |
| LPAR4 | 9606.ENSP00000408205 | WikiPathways | WP3932 | Focal adhesion: PI3K-Akt-mTOR-signaling pathway |
| LPAR4 | 9606.ENSP00000408205 | WikiPathways | WP4172 | PI3K-Akt signaling pathway |
| LPAR4 | 9606.ENSP00000408205 | WikiPathways | WP455 | GPCRs, class A rhodopsin-like |
| LPAR4 | 9606.ENSP00000408205 | WikiPathways | WP4900 | Purinergic signaling |
| LPAR4 | 9606.ENSP00000408205 | WikiPathways | WP80 | Nucleotide GPCRs |
| LPAR4 | 9606.ENSP00000408205 | COMPARTMENTS | GOCC:0005886 | Plasma membrane |
| LPAR4 | 9606.ENSP00000408205 | COMPARTMENTS | GOCC:0016020 | Membrane |
| LPAR4 | 9606.ENSP00000408205 | COMPARTMENTS | GOCC:0071944 | Cell periphery |
| LPAR4 | 9606.ENSP00000408205 | COMPARTMENTS | GOCC:0110165 | Cellular anatomical entity |
| LPAR4 | 9606.ENSP00000408205 | UniProt Keywords | KW-0297 | G-protein coupled receptor |
| LPAR4 | 9606.ENSP00000408205 | UniProt Keywords | KW-0325 | Glycoprotein |
| LPAR4 | 9606.ENSP00000408205 | UniProt Keywords | KW-0446 | Lipid-binding |
| LPAR4 | 9606.ENSP00000408205 | UniProt Keywords | KW-0472 | Membrane |
| LPAR4 | 9606.ENSP00000408205 | UniProt Keywords | KW-0675 | Receptor |
| LPAR4 | 9606.ENSP00000408205 | UniProt Keywords | KW-0807 | Transducer |
| LPAR4 | 9606.ENSP00000408205 | UniProt Keywords | KW-0812 | Transmembrane |
| LPAR4 | 9606.ENSP00000408205 | UniProt Keywords | KW-1003 | Cell membrane |
| LPAR4 | 9606.ENSP00000408205 | UniProt Keywords | KW-1015 | Disulfide bond |
| LPAR4 | 9606.ENSP00000408205 | UniProt Keywords | KW-1133 | Transmembrane helix |
| LPAR4 | 9606.ENSP00000408205 | Pfam | PF00001 | 7 transmembrane receptor (rhodopsin family) |
| LPAR4 | 9606.ENSP00000408205 | InterPro | IPR000276 | G protein-coupled receptor, rhodopsin-like |
| LPAR4 | 9606.ENSP00000408205 | InterPro | IPR017452 | GPCR, rhodopsin-like, 7TM |
| LPP | 9606.ENSP00000491657 | GO Process | GO:0007155 | Cell adhesion |

| | | | | |
|---|---|---|---|---|
| LPP | 9606.ENSP00000491657 | GO Process | GO:0009987 | Cellular process |
| LPP | 9606.ENSP00000491657 | GO Process | GO:0098609 | Cell-cell adhesion |
| LPP | 9606.ENSP00000491657 | GO Function | GO:0005488 | Binding |
| LPP | 9606.ENSP00000491657 | GO Function | GO:0043167 | Ion binding |
| LPP | 9606.ENSP00000491657 | GO Function | GO:0043169 | Cation binding |
| LPP | 9606.ENSP00000491657 | GO Function | GO:0046872 | Metal ion binding |
| LPP | 9606.ENSP00000491657 | GO Component | GO:0001725 | Stress fiber |
| LPP | 9606.ENSP00000491657 | GO Component | GO:0005622 | Intracellular anatomical structure |
| LPP | 9606.ENSP00000491657 | GO Component | GO:0005634 | Nucleus |
| LPP | 9606.ENSP00000491657 | GO Component | GO:0005737 | Cytoplasm |
| LPP | 9606.ENSP00000491657 | GO Component | GO:0005829 | Cytosol |
| LPP | 9606.ENSP00000491657 | GO Component | GO:0005856 | Cytoskeleton |
| LPP | 9606.ENSP00000491657 | GO Component | GO:0005886 | Plasma membrane |
| LPP | 9606.ENSP00000491657 | GO Component | GO:0005925 | Focal adhesion |
| LPP | 9606.ENSP00000491657 | GO Component | GO:0015629 | Actin cytoskeleton |
| LPP | 9606.ENSP00000491657 | GO Component | GO:0016020 | Membrane |
| LPP | 9606.ENSP00000491657 | GO Component | GO:0030054 | Cell junction |
| LPP | 9606.ENSP00000491657 | GO Component | GO:0030055 | Cell-substrate junction |
| LPP | 9606.ENSP00000491657 | GO Component | GO:0032432 | Actin filament bundle |
| LPP | 9606.ENSP00000491657 | GO Component | GO:0042641 | Actomyosin |
| LPP | 9606.ENSP00000491657 | GO Component | GO:0043226 | Organelle |
| LPP | 9606.ENSP00000491657 | GO Component | GO:0043227 | Membrane-bounded organelle |
| LPP | 9606.ENSP00000491657 | GO Component | GO:0043228 | Non-membrane-bounded organelle |
| LPP | 9606.ENSP00000491657 | GO Component | GO:0043229 | Intracellular organelle |
| LPP | 9606.ENSP00000491657 | GO Component | GO:0043231 | Intracellular membrane-bounded organelle |
| LPP | 9606.ENSP00000491657 | GO Component | GO:0043232 | Intracellular non-membrane-bounded organelle |
| LPP | 9606.ENSP00000491657 | GO Component | GO:0070161 | Anchoring junction |
| LPP | 9606.ENSP00000491657 | GO Component | GO:0071944 | Cell periphery |
| LPP | 9606.ENSP00000491657 | GO Component | GO:0097517 | Contractile actin filament bundle |
| LPP | 9606.ENSP00000491657 | GO Component | GO:0110165 | Cellular anatomical entity |
| LPP | 9606.ENSP00000491657 | STRING clusters | CL:37667 | Mostly uncharacterized, incl. Cell cycle regulatory protein, and EPM2A-interacting protein 1/ZBED8-like |
| LPP | 9606.ENSP00000491657 | STRING clusters | CL:37755 | Mostly uncharacterized, incl. FAM212 family, and Regulation of epithelium regeneration |

| | | | | |
|---|---|---|---|---|
| LPP | 9606.ENSP00000491657 | STRING clusters | CL:37757 | Mixed, incl. Regulation of epithelium regeneration, and FAM43A/B, phosphotyrosine-binding domain |
| LPP | 9606.ENSP00000491657 | WikiPathways | WP4560 | MFAP5 effect on permeability and motility of endothelial cells via cytoskeleton rearrangement |
| LPP | 9606.ENSP00000491657 | TISSUES | BTO:0000000 | Tissues, cell types and enzyme sources |
| LPP | 9606.ENSP00000491657 | TISSUES | BTO:0000042 | Animal |
| LPP | 9606.ENSP00000491657 | TISSUES | BTO:0000058 | Alimentary canal |
| LPP | 9606.ENSP00000491657 | TISSUES | BTO:0000081 | Reproductive system |
| LPP | 9606.ENSP00000491657 | TISSUES | BTO:0000083 | Female reproductive system |
| LPP | 9606.ENSP00000491657 | TISSUES | BTO:0000345 | Digestive gland |
| LPP | 9606.ENSP00000491657 | TISSUES | BTO:0000511 | Gastrointestinal tract |
| LPP | 9606.ENSP00000491657 | TISSUES | BTO:0000522 | Gland |
| LPP | 9606.ENSP00000491657 | TISSUES | BTO:0000648 | Intestine |
| LPP | 9606.ENSP00000491657 | TISSUES | BTO:0000651 | Small intestine |
| LPP | 9606.ENSP00000491657 | TISSUES | BTO:0001488 | Endocrine gland |
| LPP | 9606.ENSP00000491657 | TISSUES | BTO:0001489 | Whole body |
| LPP | 9606.ENSP00000491657 | TISSUES | BTO:0001491 | Viscus |
| LPP | 9606.ENSP00000491657 | TISSUES | BTO:0003091 | Urogenital system |
| LPP | 9606.ENSP00000491657 | COMPARTMENTS | GOCC:0005622 | Intracellular |
| LPP | 9606.ENSP00000491657 | COMPARTMENTS | GOCC:0005634 | Nucleus |
| LPP | 9606.ENSP00000491657 | COMPARTMENTS | GOCC:0005737 | Cytoplasm |
| LPP | 9606.ENSP00000491657 | COMPARTMENTS | GOCC:0005829 | Cytosol |
| LPP | 9606.ENSP00000491657 | COMPARTMENTS | GOCC:0005886 | Plasma membrane |
| LPP | 9606.ENSP00000491657 | COMPARTMENTS | GOCC:0005925 | Focal adhesion |
| LPP | 9606.ENSP00000491657 | COMPARTMENTS | GOCC:0016020 | Membrane |
| LPP | 9606.ENSP00000491657 | COMPARTMENTS | GOCC:0030054 | Cell junction |
| LPP | 9606.ENSP00000491657 | COMPARTMENTS | GOCC:0030055 | Cell-substrate junction |
| LPP | 9606.ENSP00000491657 | COMPARTMENTS | GOCC:0043226 | Organelle |
| LPP | 9606.ENSP00000491657 | COMPARTMENTS | GOCC:0043227 | Membrane-bounded organelle |
| LPP | 9606.ENSP00000491657 | COMPARTMENTS | GOCC:0043229 | Intracellular organelle |
| LPP | 9606.ENSP00000491657 | COMPARTMENTS | GOCC:0043231 | Intracellular membrane-bounded organelle |
| LPP | 9606.ENSP00000491657 | COMPARTMENTS | GOCC:0070161 | Anchoring junction |
| LPP | 9606.ENSP00000491657 | COMPARTMENTS | GOCC:0071944 | Cell periphery |
| LPP | 9606.ENSP00000491657 | COMPARTMENTS | GOCC:0110165 | Cellular anatomical entity |
| LPP | 9606.ENSP00000491657 | Monarch | EFO:0000246 | Age |

| | | | | |
|---|---|---|---|---|
| LPP | 9606.ENSP00000491657 | Monarch | EFO:0000313 | Carcinoma |
| LPP | 9606.ENSP00000491657 | Monarch | EFO:0000408 | Disease |
| LPP | 9606.ENSP00000491657 | Monarch | EFO:0000616 | Neoplasm |
| LPP | 9606.ENSP00000491657 | Monarch | EFO:0000684 | Respiratory system disease |
| LPP | 9606.ENSP00000491657 | Monarch | EFO:0000701 | Skin disease |
| LPP | 9606.ENSP00000491657 | Monarch | EFO:0000719 | Temporal measurement |
| LPP | 9606.ENSP00000491657 | Monarch | EFO:0003917 | Premature birth |
| LPP | 9606.ENSP00000491657 | Monarch | EFO:0004198 | Skin neoplasm |
| LPP | 9606.ENSP00000491657 | Monarch | EFO:0004300 | Longevity |
| LPP | 9606.ENSP00000491657 | Monarch | EFO:0004302 | Anthropometric measurement |
| LPP | 9606.ENSP00000491657 | Monarch | EFO:0004303 | Vital signs |
| LPP | 9606.ENSP00000491657 | Monarch | EFO:0004305 | Erythrocyte count |
| LPP | 9606.ENSP00000491657 | Monarch | EFO:0004306 | Erythrocyte indices |
| LPP | 9606.ENSP00000491657 | Monarch | EFO:0004308 | Leukocyte count |
| LPP | 9606.ENSP00000491657 | Monarch | EFO:0004324 | Body weights and measures |
| LPP | 9606.ENSP00000491657 | Monarch | EFO:0004325 | Blood pressure |
| LPP | 9606.ENSP00000491657 | Monarch | EFO:0004338 | Body weight |
| LPP | 9606.ENSP00000491657 | Monarch | EFO:0004339 | Body height |
| LPP | 9606.ENSP00000491657 | Monarch | EFO:0004340 | Body mass index |
| LPP | 9606.ENSP00000491657 | Monarch | EFO:0004352 | Mortality |
| LPP | 9606.ENSP00000491657 | Monarch | EFO:0004464 | Brain measurement |
| LPP | 9606.ENSP00000491657 | Monarch | EFO:0004467 | Insulin measurement |
| LPP | 9606.ENSP00000491657 | Monarch | EFO:0004471 | Insulin sensitivity measurement |
| LPP | 9606.ENSP00000491657 | Monarch | EFO:0004503 | Hematological measurement |
| LPP | 9606.ENSP00000491657 | Monarch | EFO:0004518 | Creatinine measurement |
| LPP | 9606.ENSP00000491657 | Monarch | EFO:0004526 | Mean corpuscular volume |
| LPP | 9606.ENSP00000491657 | Monarch | EFO:0004529 | Lipid measurement |
| LPP | 9606.ENSP00000491657 | Monarch | EFO:0004530 | Triglyceride measurement |
| LPP | 9606.ENSP00000491657 | Monarch | EFO:0004556 | Antibody measurement |
| LPP | 9606.ENSP00000491657 | Monarch | EFO:0004557 | Population measurement |
| LPP | 9606.ENSP00000491657 | Monarch | EFO:0004579 | Serum IgE measurement |
| LPP | 9606.ENSP00000491657 | Monarch | EFO:0004586 | Complete blood cell count |
| LPP | 9606.ENSP00000491657 | Monarch | EFO:0004587 | Lymphocyte count |
| LPP | 9606.ENSP00000491657 | Monarch | EFO:0004641 | White matter integrity |
| LPP | 9606.ENSP00000491657 | Monarch | EFO:0004645 | Response to vaccine |

| | | | | |
|---|---|---|---|---|
| LPP | 9606.ENSP00000491657 | Monarch | EFO:0004695 | Intraocular pressure measurement |
| LPP | 9606.ENSP00000491657 | Monarch | EFO:0004725 | Metabolite measurement |
| LPP | 9606.ENSP00000491657 | Monarch | EFO:0004730 | Hormone measurement |
| LPP | 9606.ENSP00000491657 | Monarch | EFO:0004731 | Eye measurement |
| LPP | 9606.ENSP00000491657 | Monarch | EFO:0004742 | Renal system measurement |
| LPP | 9606.ENSP00000491657 | Monarch | EFO:0004747 | Protein measurement |
| LPP | 9606.ENSP00000491657 | Monarch | EFO:0004748 | Thyroid stimulating hormone measurement |
| LPP | 9606.ENSP00000491657 | Monarch | EFO:0004833 | Neutrophil count |
| LPP | 9606.ENSP00000491657 | Monarch | EFO:0004842 | Eosinophil count |
| LPP | 9606.ENSP00000491657 | Monarch | EFO:0004847 | Age at onset |
| LPP | 9606.ENSP00000491657 | Monarch | EFO:0004870 | Sleep measurement |
| LPP | 9606.ENSP00000491657 | Monarch | EFO:0004872 | Inflammatory biomarker measurement |
| LPP | 9606.ENSP00000491657 | Monarch | EFO:0004873 | Cytokine measurement |
| LPP | 9606.ENSP00000491657 | Monarch | EFO:0004874 | Memory performance |
| LPP | 9606.ENSP00000491657 | Monarch | EFO:0005036 | Platelet measurement |
| LPP | 9606.ENSP00000491657 | Monarch | EFO:0005047 | Erythrocyte measurement |
| LPP | 9606.ENSP00000491657 | Monarch | EFO:0005090 | Basophil count |
| LPP | 9606.ENSP00000491657 | Monarch | EFO:0005091 | Monocyte count |
| LPP | 9606.ENSP00000491657 | Monarch | EFO:0005105 | Lipid or lipoprotein measurement |
| LPP | 9606.ENSP00000491657 | Monarch | EFO:0005188 | CCL11 measurement |
| LPP | 9606.ENSP00000491657 | Monarch | EFO:0005208 | Glomerular filtration rate |
| LPP | 9606.ENSP00000491657 | Monarch | EFO:0005298 | Allergic sensitization measurement |
| LPP | 9606.ENSP00000491657 | Monarch | EFO:0006336 | Diastolic blood pressure |
| LPP | 9606.ENSP00000491657 | Monarch | EFO:0006842 | Diabetes mellitus biomarker |
| LPP | 9606.ENSP00000491657 | Monarch | EFO:0006843 | Infectious disease biomarker |
| LPP | 9606.ENSP00000491657 | Monarch | EFO:0006848 | Mental or behavioural disorder biomarker |
| LPP | 9606.ENSP00000491657 | Monarch | EFO:0006858 | Epithelial neoplasm |
| LPP | 9606.ENSP00000491657 | Monarch | EFO:0006896 | Glucose homeostasis measurement |
| LPP | 9606.ENSP00000491657 | Monarch | EFO:0006917 | Spontaneous preterm birth |
| LPP | 9606.ENSP00000491657 | Monarch | EFO:0007008 | Allergy measurement |
| LPP | 9606.ENSP00000491657 | Monarch | EFO:0007010 | Drug use measurement |
| LPP | 9606.ENSP00000491657 | Monarch | EFO:0007796 | Parental longevity |
| LPP | 9606.ENSP00000491657 | Monarch | EFO:0007863 | Illness severity status |
| LPP | 9606.ENSP00000491657 | Monarch | EFO:0007874 | Gut microbiome measurement |
| LPP | 9606.ENSP00000491657 | Monarch | EFO:0007882 | Microbiome measurement |

| | | | | |
|---|---|---|---|---|
| LPP | 9606.ENSP00000491657 | Monarch | EFO:0007937 | Blood protein measurement |
| LPP | 9606.ENSP00000491657 | Monarch | EFO:0007984 | Platelet component distribution width |
| LPP | 9606.ENSP00000491657 | Monarch | EFO:0007987 | Granulocyte count |
| LPP | 9606.ENSP00000491657 | Monarch | EFO:0007988 | Myeloid white cell count |
| LPP | 9606.ENSP00000491657 | Monarch | EFO:0007989 | Monocyte percentage of leukocytes |
| LPP | 9606.ENSP00000491657 | Monarch | EFO:0007991 | Eosinophil percentage of leukocytes |
| LPP | 9606.ENSP00000491657 | Monarch | EFO:0007994 | Neutrophil percentage of granulocytes |
| LPP | 9606.ENSP00000491657 | Monarch | EFO:0007996 | Eosinophil percentage of granulocytes |
| LPP | 9606.ENSP00000491657 | Monarch | EFO:0007997 | Granulocyte percentage of myeloid white cells |
| LPP | 9606.ENSP00000491657 | Monarch | EFO:0008328 | Chronotype measurement |
| LPP | 9606.ENSP00000491657 | Monarch | EFO:0008376 | Mosquito bite measurement |
| LPP | 9606.ENSP00000491657 | Monarch | EFO:0008377 | Mosquito bite reaction itch intensity measurement |
| LPP | 9606.ENSP00000491657 | Monarch | EFO:0008378 | Mosquito bite reaction size measurement |
| LPP | 9606.ENSP00000491657 | Monarch | EFO:0009259 | Skin carcinoma |
| LPP | 9606.ENSP00000491657 | Monarch | EFO:0009260 | Non-melanoma skin carcinoma |
| LPP | 9606.ENSP00000491657 | Monarch | EFO:0009433 | Lower respiratory tract disease |
| LPP | 9606.ENSP00000491657 | Monarch | EFO:0009933 | Thyroid preparation use measurement |
| LPP | 9606.ENSP00000491657 | Monarch | EFO:0009941 | Inhalant adrenergic use measurement |
| LPP | 9606.ENSP00000491657 | Monarch | EFO:0010176 | Keratinocyte carcinoma |
| LPP | 9606.ENSP00000491657 | Monarch | EFO:0010285 | Integumentary system disease |
| LPP | 9606.ENSP00000491657 | Monarch | EFO:0010638 | Atopic asthma |
| LPP | 9606.ENSP00000491657 | Monarch | EFO:1002018 | Bronchial disease |
| LPP | 9606.ENSP00000491657 | Monarch | HP:0000118 | Phenotypic abnormality |
| LPP | 9606.ENSP00000491657 | Monarch | HP:0000951 | Abnormality of the skin |
| LPP | 9606.ENSP00000491657 | Monarch | HP:0000964 | Eczema |
| LPP | 9606.ENSP00000491657 | Monarch | HP:0001574 | Abnormality of the integument |
| LPP | 9606.ENSP00000491657 | Monarch | HP:0001871 | Abnormality of blood and blood-forming tissues |
| LPP | 9606.ENSP00000491657 | Monarch | HP:0001881 | Abnormal leukocyte morphology |
| LPP | 9606.ENSP00000491657 | Monarch | HP:0001909 | Leukemia |
| LPP | 9606.ENSP00000491657 | Monarch | HP:0002488 | Acute leukemia |
| LPP | 9606.ENSP00000491657 | Monarch | HP:0002664 | Neoplasm |
| LPP | 9606.ENSP00000491657 | Monarch | HP:0002715 | Abnormality of the immune system |
| LPP | 9606.ENSP00000491657 | Monarch | HP:0004377 | Hematological neoplasm |
| LPP | 9606.ENSP00000491657 | Monarch | HP:0004808 | Acute myeloid leukemia |
| LPP | 9606.ENSP00000491657 | Monarch | HP:0010978 | Abnormality of immune system physiology |

| | | | | |
|---|---|---|---|---|
| LPP | 9606.ENSP00000491657 | Monarch | HP:0010987 | Abnormal cellular immune system morphology |
| LPP | 9606.ENSP00000491657 | Monarch | HP:0011122 | Abnormality of skin physiology |
| LPP | 9606.ENSP00000491657 | Monarch | HP:0011123 | Inflammatory abnormality of the skin |
| LPP | 9606.ENSP00000491657 | Monarch | HP:0011793 | Neoplasm by anatomical site |
| LPP | 9606.ENSP00000491657 | Monarch | HP:0012647 | Abnormal inflammatory response |
| LPP | 9606.ENSP00000491657 | Monarch | HP:0012649 | Increased inflammatory response |
| LPP | 9606.ENSP00000491657 | Monarch | HP:0032251 | Abnormal immune system morphology |
| LPP | 9606.ENSP00000491657 | Monarch | MONDO:0000653 | Integumentary system cancer |
| LPP | 9606.ENSP00000491657 | Monarch | MONDO:0002898 | Skin cancer |
| LPP | 9606.ENSP00000491657 | Monarch | MONDO:0004979 | Asthma |
| LPP | 9606.ENSP00000491657 | Monarch | MONDO:0004992 | Cancer |
| LPP | 9606.ENSP00000491657 | Monarch | MONDO:0021634 | Epithelial skin neoplasm |
| LPP | 9606.ENSP00000491657 | Monarch | MONDO:0023370 | Neoplastic disease or syndrome |
| LPP | 9606.ENSP00000491657 | Monarch | MONDO:0045024 | Cancer or benign tumor |
| LPP | 9606.ENSP00000491657 | UniProt Keywords | KW-0007 | Acetylation |
| LPP | 9606.ENSP00000491657 | UniProt Keywords | KW-0010 | Activator |
| LPP | 9606.ENSP00000491657 | UniProt Keywords | KW-0130 | Cell adhesion |
| LPP | 9606.ENSP00000491657 | UniProt Keywords | KW-0160 | Chromosomal rearrangement |
| LPP | 9606.ENSP00000491657 | UniProt Keywords | KW-0440 | LIM domain |
| LPP | 9606.ENSP00000491657 | UniProt Keywords | KW-0472 | Membrane |
| LPP | 9606.ENSP00000491657 | UniProt Keywords | KW-0479 | Metal-binding |
| LPP | 9606.ENSP00000491657 | UniProt Keywords | KW-0539 | Nucleus |
| LPP | 9606.ENSP00000491657 | UniProt Keywords | KW-0597 | Phosphoprotein |
| LPP | 9606.ENSP00000491657 | UniProt Keywords | KW-0677 | Repeat |
| LPP | 9606.ENSP00000491657 | UniProt Keywords | KW-0832 | Ubl conjugation |
| LPP | 9606.ENSP00000491657 | UniProt Keywords | KW-0862 | Zinc |
| LPP | 9606.ENSP00000491657 | UniProt Keywords | KW-0963 | Cytoplasm |
| LPP | 9606.ENSP00000491657 | UniProt Keywords | KW-0965 | Cell junction |

| | | | | |
|---|---|---|---|---|
| LPP | 9606.ENSP00000491657 | UniProt Keywords | KW-1003 | Cell membrane |
| LPP | 9606.ENSP00000491657 | UniProt Keywords | KW-1017 | Isopeptide bond |
| LPP | 9606.ENSP00000491657 | InterPro | IPR001781 | Zinc finger, LIM-type |
| LPP | 9606.ENSP00000491657 | SMART | SM00132 | Zinc-binding domain present in Lin-11, Isl-1, Mec-3. |
| PHLDB1 | 9606.ENSP00000354498 | GO Process | GO:0010470 | Regulation of gastrulation |
| PHLDB1 | 9606.ENSP00000354498 | GO Process | GO:0010717 | Regulation of epithelial to mesenchymal transition |
| PHLDB1 | 9606.ENSP00000354498 | GO Process | GO:0022603 | Regulation of anatomical structure morphogenesis |
| PHLDB1 | 9606.ENSP00000354498 | GO Process | GO:0032886 | Regulation of microtubule-based process |
| PHLDB1 | 9606.ENSP00000354498 | GO Process | GO:0033043 | Regulation of organelle organization |
| PHLDB1 | 9606.ENSP00000354498 | GO Process | GO:0044087 | Regulation of cellular component biogenesis |
| PHLDB1 | 9606.ENSP00000354498 | GO Process | GO:0044089 | Positive regulation of cellular component biogenesis |
| PHLDB1 | 9606.ENSP00000354498 | GO Process | GO:0045595 | Regulation of cell differentiation |
| PHLDB1 | 9606.ENSP00000354498 | GO Process | GO:0045995 | Regulation of embryonic development |
| PHLDB1 | 9606.ENSP00000354498 | GO Process | GO:0048518 | Positive regulation of biological process |
| PHLDB1 | 9606.ENSP00000354498 | GO Process | GO:0048522 | Positive regulation of cellular process |
| PHLDB1 | 9606.ENSP00000354498 | GO Process | GO:0050789 | Regulation of biological process |
| PHLDB1 | 9606.ENSP00000354498 | GO Process | GO:0050793 | Regulation of developmental process |
| PHLDB1 | 9606.ENSP00000354498 | GO Process | GO:0050794 | Regulation of cellular process |
| PHLDB1 | 9606.ENSP00000354498 | GO Process | GO:0051094 | Positive regulation of developmental process |
| PHLDB1 | 9606.ENSP00000354498 | GO Process | GO:0051128 | Regulation of cellular component organization |
| PHLDB1 | 9606.ENSP00000354498 | GO Process | GO:0051130 | Positive regulation of cellular component organization |
| PHLDB1 | 9606.ENSP00000354498 | GO Process | GO:0051239 | Regulation of multicellular organismal process |
| PHLDB1 | 9606.ENSP00000354498 | GO Process | GO:0051493 | Regulation of cytoskeleton organization |
| PHLDB1 | 9606.ENSP00000354498 | GO Process | GO:0065007 | Biological regulation |
| PHLDB1 | 9606.ENSP00000354498 | GO Process | GO:0070507 | Regulation of microtubule cytoskeleton organization |
| PHLDB1 | 9606.ENSP00000354498 | GO Process | GO:0110011 | Regulation of basement membrane organization |
| PHLDB1 | 9606.ENSP00000354498 | GO Process | GO:1901201 | Regulation of extracellular matrix assembly |
| PHLDB1 | 9606.ENSP00000354498 | GO Process | GO:1901203 | Positive regulation of extracellular matrix assembly |
| PHLDB1 | 9606.ENSP00000354498 | GO Process | GO:1903053 | Regulation of extracellular matrix organization |
| PHLDB1 | 9606.ENSP00000354498 | GO Process | GO:1903055 | Positive regulation of extracellular matrix organization |
| PHLDB1 | 9606.ENSP00000354498 | GO Process | GO:1904259 | Regulation of basement membrane assembly involved in embryonic body morphogenesis |
| PHLDB1 | 9606.ENSP00000354498 | GO Process | GO:1904261 | Positive regulation of basement membrane assembly involved in embryonic body morphogenesis |
| PHLDB1 | 9606.ENSP00000354498 | GO Process | GO:2000026 | Regulation of multicellular organismal development |

| | | | | |
|---|---|---|---|---|
| PHLDB1 | 9606.ENSP00000354498 | GO Component | GO:0005576 | Extracellular region |
| PHLDB1 | 9606.ENSP00000354498 | GO Component | GO:0005622 | Intracellular anatomical structure |
| PHLDB1 | 9606.ENSP00000354498 | GO Component | GO:0005737 | Cytoplasm |
| PHLDB1 | 9606.ENSP00000354498 | GO Component | GO:0005829 | Cytosol |
| PHLDB1 | 9606.ENSP00000354498 | GO Component | GO:0005886 | Plasma membrane |
| PHLDB1 | 9606.ENSP00000354498 | GO Component | GO:0005938 | Cell cortex |
| PHLDB1 | 9606.ENSP00000354498 | GO Component | GO:0016020 | Membrane |
| PHLDB1 | 9606.ENSP00000354498 | GO Component | GO:0043226 | Organelle |
| PHLDB1 | 9606.ENSP00000354498 | GO Component | GO:0043227 | Membrane-bounded organelle |
| PHLDB1 | 9606.ENSP00000354498 | GO Component | GO:0043229 | Intracellular organelle |
| PHLDB1 | 9606.ENSP00000354498 | GO Component | GO:0043231 | Intracellular membrane-bounded organelle |
| PHLDB1 | 9606.ENSP00000354498 | GO Component | GO:0045171 | Intercellular bridge |
| PHLDB1 | 9606.ENSP00000354498 | GO Component | GO:0045178 | Basal part of cell |
| PHLDB1 | 9606.ENSP00000354498 | GO Component | GO:0045180 | Basal cortex |
| PHLDB1 | 9606.ENSP00000354498 | GO Component | GO:0071944 | Cell periphery |
| PHLDB1 | 9606.ENSP00000354498 | GO Component | GO:0099568 | Cytoplasmic region |
| PHLDB1 | 9606.ENSP00000354498 | GO Component | GO:0099738 | Cell cortex region |
| PHLDB1 | 9606.ENSP00000354498 | GO Component | GO:0110165 | Cellular anatomical entity |
| PHLDB1 | 9606.ENSP00000354498 | STRING clusters | CL:9272 | Mixed, incl. Starch and sucrose metabolism, and Amino sugar biosynthetic process |
| PHLDB1 | 9606.ENSP00000354498 | TISSUES | BTO:0000000 | Tissues, cell types and enzyme sources |
| PHLDB1 | 9606.ENSP00000354498 | TISSUES | BTO:0000042 | Animal |
| PHLDB1 | 9606.ENSP00000354498 | TISSUES | BTO:0000081 | Reproductive system |
| PHLDB1 | 9606.ENSP00000354498 | TISSUES | BTO:0000083 | Female reproductive system |
| PHLDB1 | 9606.ENSP00000354498 | TISSUES | BTO:0000142 | Brain |
| PHLDB1 | 9606.ENSP00000354498 | TISSUES | BTO:0000227 | Central nervous system |
| PHLDB1 | 9606.ENSP00000354498 | TISSUES | BTO:0000282 | Head |
| PHLDB1 | 9606.ENSP00000354498 | TISSUES | BTO:0001484 | Nervous system |
| PHLDB1 | 9606.ENSP00000354498 | TISSUES | BTO:0001489 | Whole body |
| PHLDB1 | 9606.ENSP00000354498 | TISSUES | BTO:0003091 | Urogenital system |
| PHLDB1 | 9606.ENSP00000354498 | COMPARTMENTS | GOCC:0005576 | Extracellular region |
| PHLDB1 | 9606.ENSP00000354498 | COMPARTMENTS | GOCC:0005622 | Intracellular |
| PHLDB1 | 9606.ENSP00000354498 | COMPARTMENTS | GOCC:0005737 | Cytoplasm |
| PHLDB1 | 9606.ENSP00000354498 | COMPARTMENTS | GOCC:0005829 | Cytosol |
| PHLDB1 | 9606.ENSP00000354498 | COMPARTMENTS | GOCC:0005938 | Cell cortex |

| | | | | |
|---|---|---|---|---|
| PHLDB1 | 9606.ENSP00000354498 | COMPARTMENTS | GOCC:0043226 | Organelle |
| PHLDB1 | 9606.ENSP00000354498 | COMPARTMENTS | GOCC:0043227 | Membrane-bounded organelle |
| PHLDB1 | 9606.ENSP00000354498 | COMPARTMENTS | GOCC:0043229 | Intracellular organelle |
| PHLDB1 | 9606.ENSP00000354498 | COMPARTMENTS | GOCC:0043231 | Intracellular membrane-bounded organelle |
| PHLDB1 | 9606.ENSP00000354498 | COMPARTMENTS | GOCC:0045171 | Intercellular bridge |
| PHLDB1 | 9606.ENSP00000354498 | COMPARTMENTS | GOCC:0045178 | Basal part of cell |
| PHLDB1 | 9606.ENSP00000354498 | COMPARTMENTS | GOCC:0045180 | Basal cortex |
| PHLDB1 | 9606.ENSP00000354498 | COMPARTMENTS | GOCC:0071944 | Cell periphery |
| PHLDB1 | 9606.ENSP00000354498 | COMPARTMENTS | GOCC:0099568 | Cytoplasmic region |
| PHLDB1 | 9606.ENSP00000354498 | COMPARTMENTS | GOCC:0099738 | Cell cortex region |
| PHLDB1 | 9606.ENSP00000354498 | COMPARTMENTS | GOCC:0110165 | Cellular anatomical entity |
| PHLDB1 | 9606.ENSP00000354498 | Monarch | EFO:0003923 | Bone density |
| PHLDB1 | 9606.ENSP00000354498 | Monarch | EFO:0004512 | Bone measurement |
| PHLDB1 | 9606.ENSP00000354498 | Monarch | EFO:0004516 | Bone fracture related measurement |
| PHLDB1 | 9606.ENSP00000354498 | Monarch | EFO:0004529 | Lipid measurement |
| PHLDB1 | 9606.ENSP00000354498 | Monarch | EFO:0004533 | Alkaline phosphatase measurement |
| PHLDB1 | 9606.ENSP00000354498 | Monarch | EFO:0004574 | Total cholesterol measurement |
| PHLDB1 | 9606.ENSP00000354498 | Monarch | EFO:0004582 | Liver enzyme measurement |
| PHLDB1 | 9606.ENSP00000354498 | Monarch | EFO:0004732 | Lipoprotein measurement |
| PHLDB1 | 9606.ENSP00000354498 | Monarch | EFO:0004747 | Protein measurement |
| PHLDB1 | 9606.ENSP00000354498 | Monarch | EFO:0005105 | Lipid or lipoprotein measurement |
| PHLDB1 | 9606.ENSP00000354498 | Monarch | EFO:0009270 | Heel bone mineral density |
| PHLDB1 | 9606.ENSP00000354498 | UniProt Keywords | KW-0025 | Alternative splicing |
| PHLDB1 | 9606.ENSP00000354498 | UniProt Keywords | KW-0175 | Coiled coil |
| PHLDB1 | 9606.ENSP00000354498 | UniProt Keywords | KW-0488 | Methylation |
| PHLDB1 | 9606.ENSP00000354498 | UniProt Keywords | KW-0597 | Phosphoprotein |
| PHLDB1 | 9606.ENSP00000354498 | InterPro | IPR001849 | Pleckstrin homology domain |
| PHLDB1 | 9606.ENSP00000354498 | InterPro | IPR008984 | SMAD/FHA domain superfamily |
| PHLDB1 | 9606.ENSP00000354498 | InterPro | IPR011993 | PH-like domain superfamily |
| PHLDB1 | 9606.ENSP00000354498 | InterPro | IPR037810 | PHLDB1/2/3, PH domain |
| PHLDB1 | 9606.ENSP00000354498 | SMART | SM00233 | Pleckstrin homology domain. |
| PXYLP1 | 9606.ENSP00000286353 | GO Process | GO:0006022 | Aminoglycan metabolic process |
| PXYLP1 | 9606.ENSP00000286353 | GO Process | GO:0006023 | Aminoglycan biosynthetic process |
| PXYLP1 | 9606.ENSP00000286353 | GO Process | GO:0006024 | Glycosaminoglycan biosynthetic process |
| PXYLP1 | 9606.ENSP00000286353 | GO Process | GO:0006029 | Proteoglycan metabolic process |

| | | | | |
|---|---|---|---|---|
| PXYLP1 | 9606.ENSP00000286353 | GO Process | GO:0006793 | Phosphorus metabolic process |
| PXYLP1 | 9606.ENSP00000286353 | GO Process | GO:0006796 | Phosphate-containing compound metabolic process |
| PXYLP1 | 9606.ENSP00000286353 | GO Process | GO:0006807 | Nitrogen compound metabolic process |
| PXYLP1 | 9606.ENSP00000286353 | GO Process | GO:0008152 | Metabolic process |
| PXYLP1 | 9606.ENSP00000286353 | GO Process | GO:0009058 | Biosynthetic process |
| PXYLP1 | 9606.ENSP00000286353 | GO Process | GO:0009059 | Macromolecule biosynthetic process |
| PXYLP1 | 9606.ENSP00000286353 | GO Process | GO:0009100 | Glycoprotein metabolic process |
| PXYLP1 | 9606.ENSP00000286353 | GO Process | GO:0009101 | Glycoprotein biosynthetic process |
| PXYLP1 | 9606.ENSP00000286353 | GO Process | GO:0009889 | Regulation of biosynthetic process |
| PXYLP1 | 9606.ENSP00000286353 | GO Process | GO:0009891 | Positive regulation of biosynthetic process |
| PXYLP1 | 9606.ENSP00000286353 | GO Process | GO:0009893 | Positive regulation of metabolic process |
| PXYLP1 | 9606.ENSP00000286353 | GO Process | GO:0009987 | Cellular process |
| PXYLP1 | 9606.ENSP00000286353 | GO Process | GO:0010556 | Regulation of macromolecule biosynthetic process |
| PXYLP1 | 9606.ENSP00000286353 | GO Process | GO:0010557 | Positive regulation of macromolecule biosynthetic process |
| PXYLP1 | 9606.ENSP00000286353 | GO Process | GO:0010559 | Regulation of glycoprotein biosynthetic process |
| PXYLP1 | 9606.ENSP00000286353 | GO Process | GO:0010560 | Positive regulation of glycoprotein biosynthetic process |
| PXYLP1 | 9606.ENSP00000286353 | GO Process | GO:0010604 | Positive regulation of macromolecule metabolic process |
| PXYLP1 | 9606.ENSP00000286353 | GO Process | GO:0010908 | Regulation of heparan sulfate proteoglycan biosynthetic process |
| PXYLP1 | 9606.ENSP00000286353 | GO Process | GO:0010909 | Positive regulation of heparan sulfate proteoglycan biosynthetic process |
| PXYLP1 | 9606.ENSP00000286353 | GO Process | GO:0016311 | Dephosphorylation |
| PXYLP1 | 9606.ENSP00000286353 | GO Process | GO:0019222 | Regulation of metabolic process |
| PXYLP1 | 9606.ENSP00000286353 | GO Process | GO:0019538 | Protein metabolic process |
| PXYLP1 | 9606.ENSP00000286353 | GO Process | GO:0030166 | Proteoglycan biosynthetic process |
| PXYLP1 | 9606.ENSP00000286353 | GO Process | GO:0030203 | Glycosaminoglycan metabolic process |
| PXYLP1 | 9606.ENSP00000286353 | GO Process | GO:0031323 | Regulation of cellular metabolic process |
| PXYLP1 | 9606.ENSP00000286353 | GO Process | GO:0031325 | Positive regulation of cellular metabolic process |
| PXYLP1 | 9606.ENSP00000286353 | GO Process | GO:0031326 | Regulation of cellular biosynthetic process |
| PXYLP1 | 9606.ENSP00000286353 | GO Process | GO:0031328 | Positive regulation of cellular biosynthetic process |
| PXYLP1 | 9606.ENSP00000286353 | GO Process | GO:0034645 | Cellular macromolecule biosynthetic process |
| PXYLP1 | 9606.ENSP00000286353 | GO Process | GO:0043170 | Macromolecule metabolic process |
| PXYLP1 | 9606.ENSP00000286353 | GO Process | GO:0044237 | Cellular metabolic process |
| PXYLP1 | 9606.ENSP00000286353 | GO Process | GO:0044238 | Primary metabolic process |
| PXYLP1 | 9606.ENSP00000286353 | GO Process | GO:0044249 | Cellular biosynthetic process |
| PXYLP1 | 9606.ENSP00000286353 | GO Process | GO:0044260 | Cellular macromolecule metabolic process |
| PXYLP1 | 9606.ENSP00000286353 | GO Process | GO:0048518 | Positive regulation of biological process |

| | | | | |
|---|---|---|---|---|
| PXYLP1 | 9606.ENSP00000286353 | GO Process | GO:0048522 | Positive regulation of cellular process |
| PXYLP1 | 9606.ENSP00000286353 | GO Process | GO:0050650 | Chondroitin sulfate proteoglycan biosynthetic process |
| PXYLP1 | 9606.ENSP00000286353 | GO Process | GO:0050654 | Chondroitin sulfate proteoglycan metabolic process |
| PXYLP1 | 9606.ENSP00000286353 | GO Process | GO:0050789 | Regulation of biological process |
| PXYLP1 | 9606.ENSP00000286353 | GO Process | GO:0050794 | Regulation of cellular process |
| PXYLP1 | 9606.ENSP00000286353 | GO Process | GO:0051171 | Regulation of nitrogen compound metabolic process |
| PXYLP1 | 9606.ENSP00000286353 | GO Process | GO:0051173 | Positive regulation of nitrogen compound metabolic process |
| PXYLP1 | 9606.ENSP00000286353 | GO Process | GO:0051246 | Regulation of protein metabolic process |
| PXYLP1 | 9606.ENSP00000286353 | GO Process | GO:0051247 | Positive regulation of protein metabolic process |
| PXYLP1 | 9606.ENSP00000286353 | GO Process | GO:0060255 | Regulation of macromolecule metabolic process |
| PXYLP1 | 9606.ENSP00000286353 | GO Process | GO:0065007 | Biological regulation |
| PXYLP1 | 9606.ENSP00000286353 | GO Process | GO:0071704 | Organic substance metabolic process |
| PXYLP1 | 9606.ENSP00000286353 | GO Process | GO:0080090 | Regulation of primary metabolic process |
| PXYLP1 | 9606.ENSP00000286353 | GO Process | GO:1901135 | Carbohydrate derivative metabolic process |
| PXYLP1 | 9606.ENSP00000286353 | GO Process | GO:1901137 | Carbohydrate derivative biosynthetic process |
| PXYLP1 | 9606.ENSP00000286353 | GO Process | GO:1901564 | Organonitrogen compound metabolic process |
| PXYLP1 | 9606.ENSP00000286353 | GO Process | GO:1901566 | Organonitrogen compound biosynthetic process |
| PXYLP1 | 9606.ENSP00000286353 | GO Process | GO:1901576 | Organic substance biosynthetic process |
| PXYLP1 | 9606.ENSP00000286353 | GO Process | GO:1902730 | Positive regulation of proteoglycan biosynthetic process |
| PXYLP1 | 9606.ENSP00000286353 | GO Process | GO:1903018 | Regulation of glycoprotein metabolic process |
| PXYLP1 | 9606.ENSP00000286353 | GO Process | GO:1903020 | Positive regulation of glycoprotein metabolic process |
| PXYLP1 | 9606.ENSP00000286353 | GO Process | GO:2000112 | Regulation of cellular macromolecule biosynthetic process |
| PXYLP1 | 9606.ENSP00000286353 | GO Function | GO:0003824 | Catalytic activity |
| PXYLP1 | 9606.ENSP00000286353 | GO Function | GO:0016787 | Hydrolase activity |
| PXYLP1 | 9606.ENSP00000286353 | GO Function | GO:0016788 | Hydrolase activity, acting on ester bonds |
| PXYLP1 | 9606.ENSP00000286353 | GO Function | GO:0016791 | Phosphatase activity |
| PXYLP1 | 9606.ENSP00000286353 | GO Function | GO:0042578 | Phosphoric ester hydrolase activity |
| PXYLP1 | 9606.ENSP00000286353 | GO Component | GO:0000139 | Golgi membrane |
| PXYLP1 | 9606.ENSP00000286353 | GO Component | GO:0005622 | Intracellular anatomical structure |
| PXYLP1 | 9606.ENSP00000286353 | GO Component | GO:0005737 | Cytoplasm |
| PXYLP1 | 9606.ENSP00000286353 | GO Component | GO:0005794 | Golgi apparatus |
| PXYLP1 | 9606.ENSP00000286353 | GO Component | GO:0012505 | Endomembrane system |
| PXYLP1 | 9606.ENSP00000286353 | GO Component | GO:0016020 | Membrane |
| PXYLP1 | 9606.ENSP00000286353 | GO Component | GO:0016021 | Integral component of membrane |
| PXYLP1 | 9606.ENSP00000286353 | GO Component | GO:0031090 | Organelle membrane |

| | | | | |
|---|---|---|---|---|
| PXYLP1 | 9606.ENSP00000286353 | GO Component | GO:0031224 | Intrinsic component of membrane |
| PXYLP1 | 9606.ENSP00000286353 | GO Component | GO:0043226 | Organelle |
| PXYLP1 | 9606.ENSP00000286353 | GO Component | GO:0043227 | Membrane-bounded organelle |
| PXYLP1 | 9606.ENSP00000286353 | GO Component | GO:0043229 | Intracellular organelle |
| PXYLP1 | 9606.ENSP00000286353 | GO Component | GO:0043231 | Intracellular membrane-bounded organelle |
| PXYLP1 | 9606.ENSP00000286353 | GO Component | GO:0098588 | Bounding membrane of organelle |
| PXYLP1 | 9606.ENSP00000286353 | GO Component | GO:0110165 | Cellular anatomical entity |
| PXYLP1 | 9606.ENSP00000286353 | STRING clusters | CL:8023 | Mixed, incl. Protein serine/threonine phosphatase complex, and Phosphatase regulator activity |
| PXYLP1 | 9606.ENSP00000286353 | STRING clusters | CL:8253 | Mixed, incl. Cyclin-dependent protein kinase holoenzyme complex, and Cyclin-dependent protein kinase activity |
| PXYLP1 | 9606.ENSP00000286353 | STRING clusters | CL:8255 | Mixed, incl. Cyclin-dependent protein kinase holoenzyme complex, and Dual specificity phosphatase, catalytic domain |
| PXYLP1 | 9606.ENSP00000286353 | STRING clusters | CL:8257 | Mixed, incl. Cyclin-dependent protein kinase holoenzyme complex, and Dual specificity phosphatase, catalytic domain |
| PXYLP1 | 9606.ENSP00000286353 | STRING clusters | CL:8327 | Mixed, incl. Ubiquitin family, and Domain of unknown function (DUF4062) |
| PXYLP1 | 9606.ENSP00000286353 | TISSUES | BTO:0000000 | Tissues, cell types and enzyme sources |
| PXYLP1 | 9606.ENSP00000286353 | TISSUES | BTO:0000042 | Animal |
| PXYLP1 | 9606.ENSP00000286353 | TISSUES | BTO:0000081 | Reproductive system |
| PXYLP1 | 9606.ENSP00000286353 | TISSUES | BTO:0000083 | Female reproductive system |
| PXYLP1 | 9606.ENSP00000286353 | TISSUES | BTO:0000254 | Female reproductive gland |
| PXYLP1 | 9606.ENSP00000286353 | TISSUES | BTO:0000522 | Gland |
| PXYLP1 | 9606.ENSP00000286353 | TISSUES | BTO:0000975 | Ovary |
| PXYLP1 | 9606.ENSP00000286353 | TISSUES | BTO:0001488 | Endocrine gland |
| PXYLP1 | 9606.ENSP00000286353 | TISSUES | BTO:0001489 | Whole body |
| PXYLP1 | 9606.ENSP00000286353 | TISSUES | BTO:0003091 | Urogenital system |
| PXYLP1 | 9606.ENSP00000286353 | TISSUES | BTO:0003099 | Internal female genital organ |
| PXYLP1 | 9606.ENSP00000286353 | TISSUES | BTO:0004617 | Mucilage |
| PXYLP1 | 9606.ENSP00000286353 | COMPARTMENTS | GOCC:0005618 | Cell wall |
| PXYLP1 | 9606.ENSP00000286353 | COMPARTMENTS | GOCC:0005622 | Intracellular |
| PXYLP1 | 9606.ENSP00000286353 | COMPARTMENTS | GOCC:0005737 | Cytoplasm |
| PXYLP1 | 9606.ENSP00000286353 | COMPARTMENTS | GOCC:0005794 | Golgi apparatus |
| PXYLP1 | 9606.ENSP00000286353 | COMPARTMENTS | GOCC:0009505 | Plant-type cell wall |
| PXYLP1 | 9606.ENSP00000286353 | COMPARTMENTS | GOCC:0009530 | Primary cell wall |
| PXYLP1 | 9606.ENSP00000286353 | COMPARTMENTS | GOCC:0009531 | Secondary cell wall |

| | | | | |
|---|---|---|---|---|
| PXYLP1 | 9606.ENSP00000286353 | COMPARTMENTS | GOCC:0009549 | Cellulose microfibril |
| PXYLP1 | 9606.ENSP00000286353 | COMPARTMENTS | GOCC:0012505 | Endomembrane system |
| PXYLP1 | 9606.ENSP00000286353 | COMPARTMENTS | GOCC:0030312 | External encapsulating structure |
| PXYLP1 | 9606.ENSP00000286353 | COMPARTMENTS | GOCC:0043226 | Organelle |
| PXYLP1 | 9606.ENSP00000286353 | COMPARTMENTS | GOCC:0043227 | Membrane-bounded organelle |
| PXYLP1 | 9606.ENSP00000286353 | COMPARTMENTS | GOCC:0043229 | Intracellular organelle |
| PXYLP1 | 9606.ENSP00000286353 | COMPARTMENTS | GOCC:0043231 | Intracellular membrane-bounded organelle |
| PXYLP1 | 9606.ENSP00000286353 | COMPARTMENTS | GOCC:0071944 | Cell periphery |
| PXYLP1 | 9606.ENSP00000286353 | COMPARTMENTS | GOCC:0110165 | Cellular anatomical entity |
| PXYLP1 | 9606.ENSP00000286353 | Monarch | EFO:0003926 | Neuropsychological test |
| PXYLP1 | 9606.ENSP00000286353 | Monarch | EFO:0004298 | Cardiovascular measurement |
| PXYLP1 | 9606.ENSP00000286353 | Monarch | EFO:0004302 | Anthropometric measurement |
| PXYLP1 | 9606.ENSP00000286353 | Monarch | EFO:0004306 | Erythrocyte indices |
| PXYLP1 | 9606.ENSP00000286353 | Monarch | EFO:0004308 | Leukocyte count |
| PXYLP1 | 9606.ENSP00000286353 | Monarch | EFO:0004324 | Body weights and measures |
| PXYLP1 | 9606.ENSP00000286353 | Monarch | EFO:0004338 | Body weight |
| PXYLP1 | 9606.ENSP00000286353 | Monarch | EFO:0004339 | Body height |
| PXYLP1 | 9606.ENSP00000286353 | Monarch | EFO:0004342 | Waist circumference |
| PXYLP1 | 9606.ENSP00000286353 | Monarch | EFO:0004468 | Glucose measurement |
| PXYLP1 | 9606.ENSP00000286353 | Monarch | EFO:0004503 | Hematological measurement |
| PXYLP1 | 9606.ENSP00000286353 | Monarch | EFO:0004509 | Hemoglobin measurement |
| PXYLP1 | 9606.ENSP00000286353 | Monarch | EFO:0004512 | Bone measurement |
| PXYLP1 | 9606.ENSP00000286353 | Monarch | EFO:0004527 | Mean corpuscular hemoglobin |
| PXYLP1 | 9606.ENSP00000286353 | Monarch | EFO:0004529 | Lipid measurement |
| PXYLP1 | 9606.ENSP00000286353 | Monarch | EFO:0004541 | HbA1c measurement |
| PXYLP1 | 9606.ENSP00000286353 | Monarch | EFO:0004555 | Glycoprotein measurement |
| PXYLP1 | 9606.ENSP00000286353 | Monarch | EFO:0004586 | Complete blood cell count |
| PXYLP1 | 9606.ENSP00000286353 | Monarch | EFO:0004747 | Protein measurement |
| PXYLP1 | 9606.ENSP00000286353 | Monarch | EFO:0004833 | Neutrophil count |
| PXYLP1 | 9606.ENSP00000286353 | Monarch | EFO:0004872 | Inflammatory biomarker measurement |
| PXYLP1 | 9606.ENSP00000286353 | Monarch | EFO:0005047 | Erythrocyte measurement |
| PXYLP1 | 9606.ENSP00000286353 | Monarch | EFO:0005090 | Basophil count |
| PXYLP1 | 9606.ENSP00000286353 | Monarch | EFO:0005093 | Hip circumference |
| PXYLP1 | 9606.ENSP00000286353 | Monarch | EFO:0005105 | Lipid or lipoprotein measurement |
| PXYLP1 | 9606.ENSP00000286353 | Monarch | EFO:0005278 | Cardiovascular disease biomarker measurement |

| | | | | |
|---|---|---|---|---|
| PXYLP1 | 9606.ENSP00000286353 | Monarch | EFO:0006842 | Diabetes mellitus biomarker |
| PXYLP1 | 9606.ENSP00000286353 | Monarch | EFO:0006848 | Mental or behavioural disorder biomarker |
| PXYLP1 | 9606.ENSP00000286353 | Monarch | EFO:0007789 | BMI-adjusted waist circumference |
| PXYLP1 | 9606.ENSP00000286353 | Monarch | EFO:0007819 | Advanced glycation end-product measurement |
| PXYLP1 | 9606.ENSP00000286353 | Monarch | EFO:0007987 | Granulocyte count |
| PXYLP1 | 9606.ENSP00000286353 | Monarch | EFO:0007988 | Myeloid white cell count |
| PXYLP1 | 9606.ENSP00000286353 | Monarch | EFO:0008039 | BMI-adjusted hip circumference |
| PXYLP1 | 9606.ENSP00000286353 | Monarch | EFO:0010226 | Phosphatidylcholine measurement |
| PXYLP1 | 9606.ENSP00000286353 | Monarch | EFO:0010384 | Phosphatidylcholine 38:2 measurement |
| PXYLP1 | 9606.ENSP00000286353 | Monarch | EFO:0010473 | Cyclic adenosine monophosphate measurement |
| PXYLP1 | 9606.ENSP00000286353 | Monarch | EFO:0010513 | Nucleotide measurement |
| PXYLP1 | 9606.ENSP00000286353 | Monarch | EFO:0010968 | Phosphate measurement |
| PXYLP1 | 9606.ENSP00000286353 | Monarch | HP:0000118 | Phenotypic abnormality |
| PXYLP1 | 9606.ENSP00000286353 | Monarch | HP:0000478 | Abnormality of the eye |
| PXYLP1 | 9606.ENSP00000286353 | Monarch | HP:0000539 | Abnormality of refraction |
| PXYLP1 | 9606.ENSP00000286353 | Monarch | HP:0000545 | Myopia |
| PXYLP1 | 9606.ENSP00000286353 | Monarch | HP:0012373 | Abnormal eye physiology |
| PXYLP1 | 9606.ENSP00000286353 | UniProt Keywords | KW-0025 | Alternative splicing |
| PXYLP1 | 9606.ENSP00000286353 | UniProt Keywords | KW-0325 | Glycoprotein |
| PXYLP1 | 9606.ENSP00000286353 | UniProt Keywords | KW-0333 | Golgi apparatus |
| PXYLP1 | 9606.ENSP00000286353 | UniProt Keywords | KW-0378 | Hydrolase |
| PXYLP1 | 9606.ENSP00000286353 | UniProt Keywords | KW-0472 | Membrane |
| PXYLP1 | 9606.ENSP00000286353 | UniProt Keywords | KW-0735 | Signal-anchor |
| PXYLP1 | 9606.ENSP00000286353 | UniProt Keywords | KW-0812 | Transmembrane |
| PXYLP1 | 9606.ENSP00000286353 | UniProt Keywords | KW-1133 | Transmembrane helix |
| PXYLP1 | 9606.ENSP00000286353 | Pfam | PF00328 | Histidine phosphatase superfamily (branch 2) |
| PXYLP1 | 9606.ENSP00000286353 | InterPro | IPR000560 | Histidine phosphatase superfamily, clade-2 |
| PXYLP1 | 9606.ENSP00000286353 | InterPro | IPR029033 | Histidine phosphatase superfamily |
| RHOBTB3 | 9606.ENSP00000369318 | GO Process | GO:0000003 | Reproduction |
| RHOBTB3 | 9606.ENSP00000369318 | GO Process | GO:0003006 | Developmental process involved in reproduction |
| RHOBTB3 | 9606.ENSP00000369318 | GO Process | GO:0006810 | Transport |
| RHOBTB3 | 9606.ENSP00000369318 | GO Process | GO:0006996 | Organelle organization |
| RHOBTB3 | 9606.ENSP00000369318 | GO Process | GO:0007010 | Cytoskeleton organization |
| RHOBTB3 | 9606.ENSP00000369318 | GO Process | GO:0007015 | Actin filament organization |
| RHOBTB3 | 9606.ENSP00000369318 | GO Process | GO:0007154 | Cell communication |

| | | | | |
|---|---|---|---|---|
| RHOBTB3 | 9606.ENSP00000369318 | GO Process | GO:0007163 | Establishment or maintenance of cell polarity |
| RHOBTB3 | 9606.ENSP00000369318 | GO Process | GO:0007165 | Signal transduction |
| RHOBTB3 | 9606.ENSP00000369318 | GO Process | GO:0007264 | Small GTPase mediated signal transduction |
| RHOBTB3 | 9606.ENSP00000369318 | GO Process | GO:0007275 | Multicellular organism development |
| RHOBTB3 | 9606.ENSP00000369318 | GO Process | GO:0007548 | Sex differentiation |
| RHOBTB3 | 9606.ENSP00000369318 | GO Process | GO:0008360 | Regulation of cell shape |
| RHOBTB3 | 9606.ENSP00000369318 | GO Process | GO:0008406 | Gonad development |
| RHOBTB3 | 9606.ENSP00000369318 | GO Process | GO:0008584 | Male gonad development |
| RHOBTB3 | 9606.ENSP00000369318 | GO Process | GO:0009987 | Cellular process |
| RHOBTB3 | 9606.ENSP00000369318 | GO Process | GO:0016043 | Cellular component organization |
| RHOBTB3 | 9606.ENSP00000369318 | GO Process | GO:0016192 | Vesicle-mediated transport |
| RHOBTB3 | 9606.ENSP00000369318 | GO Process | GO:0016197 | Endosomal transport |
| RHOBTB3 | 9606.ENSP00000369318 | GO Process | GO:0016477 | Cell migration |
| RHOBTB3 | 9606.ENSP00000369318 | GO Process | GO:0016482 | Cytosolic transport |
| RHOBTB3 | 9606.ENSP00000369318 | GO Process | GO:0022414 | Reproductive process |
| RHOBTB3 | 9606.ENSP00000369318 | GO Process | GO:0022603 | Regulation of anatomical structure morphogenesis |
| RHOBTB3 | 9606.ENSP00000369318 | GO Process | GO:0022604 | Regulation of cell morphogenesis |
| RHOBTB3 | 9606.ENSP00000369318 | GO Process | GO:0023052 | Signaling |
| RHOBTB3 | 9606.ENSP00000369318 | GO Process | GO:0030029 | Actin filament-based process |
| RHOBTB3 | 9606.ENSP00000369318 | GO Process | GO:0030036 | Actin cytoskeleton organization |
| RHOBTB3 | 9606.ENSP00000369318 | GO Process | GO:0030865 | Cortical cytoskeleton organization |
| RHOBTB3 | 9606.ENSP00000369318 | GO Process | GO:0032501 | Multicellular organismal process |
| RHOBTB3 | 9606.ENSP00000369318 | GO Process | GO:0032502 | Developmental process |
| RHOBTB3 | 9606.ENSP00000369318 | GO Process | GO:0032956 | Regulation of actin cytoskeleton organization |
| RHOBTB3 | 9606.ENSP00000369318 | GO Process | GO:0032970 | Regulation of actin filament-based process |
| RHOBTB3 | 9606.ENSP00000369318 | GO Process | GO:0033043 | Regulation of organelle organization |
| RHOBTB3 | 9606.ENSP00000369318 | GO Process | GO:0035556 | Intracellular signal transduction |
| RHOBTB3 | 9606.ENSP00000369318 | GO Process | GO:0042147 | Retrograde transport, endosome to Golgi |
| RHOBTB3 | 9606.ENSP00000369318 | GO Process | GO:0045137 | Development of primary sexual characteristics |
| RHOBTB3 | 9606.ENSP00000369318 | GO Process | GO:0046546 | Development of primary male sexual characteristics |
| RHOBTB3 | 9606.ENSP00000369318 | GO Process | GO:0046661 | Male sex differentiation |
| RHOBTB3 | 9606.ENSP00000369318 | GO Process | GO:0046907 | Intracellular transport |
| RHOBTB3 | 9606.ENSP00000369318 | GO Process | GO:0048513 | Animal organ development |
| RHOBTB3 | 9606.ENSP00000369318 | GO Process | GO:0048608 | Reproductive structure development |
| RHOBTB3 | 9606.ENSP00000369318 | GO Process | GO:0048731 | System development |

| | | | | |
|---|---|---|---|---|
| RHOBTB3 | 9606.ENSP00000369318 | GO Process | GO:0048856 | Anatomical structure development |
| RHOBTB3 | 9606.ENSP00000369318 | GO Process | GO:0048870 | Cell motility |
| RHOBTB3 | 9606.ENSP00000369318 | GO Process | GO:0050789 | Regulation of biological process |
| RHOBTB3 | 9606.ENSP00000369318 | GO Process | GO:0050793 | Regulation of developmental process |
| RHOBTB3 | 9606.ENSP00000369318 | GO Process | GO:0050794 | Regulation of cellular process |
| RHOBTB3 | 9606.ENSP00000369318 | GO Process | GO:0050896 | Response to stimulus |
| RHOBTB3 | 9606.ENSP00000369318 | GO Process | GO:0051128 | Regulation of cellular component organization |
| RHOBTB3 | 9606.ENSP00000369318 | GO Process | GO:0051179 | Localization |
| RHOBTB3 | 9606.ENSP00000369318 | GO Process | GO:0051234 | Establishment of localization |
| RHOBTB3 | 9606.ENSP00000369318 | GO Process | GO:0051493 | Regulation of cytoskeleton organization |
| RHOBTB3 | 9606.ENSP00000369318 | GO Process | GO:0051641 | Cellular localization |
| RHOBTB3 | 9606.ENSP00000369318 | GO Process | GO:0051649 | Establishment of localization in cell |
| RHOBTB3 | 9606.ENSP00000369318 | GO Process | GO:0051716 | Cellular response to stimulus |
| RHOBTB3 | 9606.ENSP00000369318 | GO Process | GO:0061458 | Reproductive system development |
| RHOBTB3 | 9606.ENSP00000369318 | GO Process | GO:0065007 | Biological regulation |
| RHOBTB3 | 9606.ENSP00000369318 | GO Process | GO:0071840 | Cellular component organization or biogenesis |
| RHOBTB3 | 9606.ENSP00000369318 | GO Process | GO:0097435 | Supramolecular fiber organization |
| RHOBTB3 | 9606.ENSP00000369318 | GO Function | GO:0000166 | Nucleotide binding |
| RHOBTB3 | 9606.ENSP00000369318 | GO Function | GO:0003824 | Catalytic activity |
| RHOBTB3 | 9606.ENSP00000369318 | GO Function | GO:0003924 | GTPase activity |
| RHOBTB3 | 9606.ENSP00000369318 | GO Function | GO:0005488 | Binding |
| RHOBTB3 | 9606.ENSP00000369318 | GO Function | GO:0005515 | Protein binding |
| RHOBTB3 | 9606.ENSP00000369318 | GO Function | GO:0005524 | ATP binding |
| RHOBTB3 | 9606.ENSP00000369318 | GO Function | GO:0005525 | GTP binding |
| RHOBTB3 | 9606.ENSP00000369318 | GO Function | GO:0016462 | Pyrophosphatase activity |
| RHOBTB3 | 9606.ENSP00000369318 | GO Function | GO:0016787 | Hydrolase activity |
| RHOBTB3 | 9606.ENSP00000369318 | GO Function | GO:0016817 | Hydrolase activity, acting on acid anhydrides |
| RHOBTB3 | 9606.ENSP00000369318 | GO Function | GO:0016818 | Hydrolase activity, acting on acid anhydrides, in phosphorus-containing anhydrides |
| RHOBTB3 | 9606.ENSP00000369318 | GO Function | GO:0016887 | ATP hydrolysis activity |
| RHOBTB3 | 9606.ENSP00000369318 | GO Function | GO:0017076 | Purine nucleotide binding |
| RHOBTB3 | 9606.ENSP00000369318 | GO Function | GO:0017111 | Nucleoside-triphosphatase activity |
| RHOBTB3 | 9606.ENSP00000369318 | GO Function | GO:0019001 | Guanyl nucleotide binding |
| RHOBTB3 | 9606.ENSP00000369318 | GO Function | GO:0019899 | Enzyme binding |
| RHOBTB3 | 9606.ENSP00000369318 | GO Function | GO:0019900 | Kinase binding |

| | | | | |
|---|---|---|---|---|
| RHOBTB3 | 9606.ENSP00000369318 | GO Function | GO:0019901 | Protein kinase binding |
| RHOBTB3 | 9606.ENSP00000369318 | GO Function | GO:0030554 | Adenyl nucleotide binding |
| RHOBTB3 | 9606.ENSP00000369318 | GO Function | GO:0031267 | Small GTPase binding |
| RHOBTB3 | 9606.ENSP00000369318 | GO Function | GO:0032553 | Ribonucleotide binding |
| RHOBTB3 | 9606.ENSP00000369318 | GO Function | GO:0032555 | Purine ribonucleotide binding |
| RHOBTB3 | 9606.ENSP00000369318 | GO Function | GO:0032559 | Adenyl ribonucleotide binding |
| RHOBTB3 | 9606.ENSP00000369318 | GO Function | GO:0032561 | Guanyl ribonucleotide binding |
| RHOBTB3 | 9606.ENSP00000369318 | GO Function | GO:0035639 | Purine ribonucleoside triphosphate binding |
| RHOBTB3 | 9606.ENSP00000369318 | GO Function | GO:0036094 | Small molecule binding |
| RHOBTB3 | 9606.ENSP00000369318 | GO Function | GO:0043167 | Ion binding |
| RHOBTB3 | 9606.ENSP00000369318 | GO Function | GO:0043168 | Anion binding |
| RHOBTB3 | 9606.ENSP00000369318 | GO Function | GO:0051020 | GTPase binding |
| RHOBTB3 | 9606.ENSP00000369318 | GO Function | GO:0097159 | Organic cyclic compound binding |
| RHOBTB3 | 9606.ENSP00000369318 | GO Function | GO:0097367 | Carbohydrate derivative binding |
| RHOBTB3 | 9606.ENSP00000369318 | GO Function | GO:0140657 | ATP-dependent activity |
| RHOBTB3 | 9606.ENSP00000369318 | GO Function | GO:1901265 | Nucleoside phosphate binding |
| RHOBTB3 | 9606.ENSP00000369318 | GO Function | GO:1901363 | Heterocyclic compound binding |
| RHOBTB3 | 9606.ENSP00000369318 | GO Component | GO:0005576 | Extracellular region |
| RHOBTB3 | 9606.ENSP00000369318 | GO Component | GO:0005615 | Extracellular space |
| RHOBTB3 | 9606.ENSP00000369318 | GO Component | GO:0005622 | Intracellular anatomical structure |
| RHOBTB3 | 9606.ENSP00000369318 | GO Component | GO:0005737 | Cytoplasm |
| RHOBTB3 | 9606.ENSP00000369318 | GO Component | GO:0005794 | Golgi apparatus |
| RHOBTB3 | 9606.ENSP00000369318 | GO Component | GO:0005802 | trans-Golgi network |
| RHOBTB3 | 9606.ENSP00000369318 | GO Component | GO:0005829 | Cytosol |
| RHOBTB3 | 9606.ENSP00000369318 | GO Component | GO:0005856 | Cytoskeleton |
| RHOBTB3 | 9606.ENSP00000369318 | GO Component | GO:0005886 | Plasma membrane |
| RHOBTB3 | 9606.ENSP00000369318 | GO Component | GO:0005938 | Cell cortex |
| RHOBTB3 | 9606.ENSP00000369318 | GO Component | GO:0012505 | Endomembrane system |
| RHOBTB3 | 9606.ENSP00000369318 | GO Component | GO:0016020 | Membrane |
| RHOBTB3 | 9606.ENSP00000369318 | GO Component | GO:0031090 | Organelle membrane |
| RHOBTB3 | 9606.ENSP00000369318 | GO Component | GO:0031410 | Cytoplasmic vesicle |
| RHOBTB3 | 9606.ENSP00000369318 | GO Component | GO:0031982 | Vesicle |
| RHOBTB3 | 9606.ENSP00000369318 | GO Component | GO:0031984 | Organelle subcompartment |
| RHOBTB3 | 9606.ENSP00000369318 | GO Component | GO:0032588 | trans-Golgi network membrane |
| RHOBTB3 | 9606.ENSP00000369318 | GO Component | GO:0042995 | Cell projection |

| | | | | |
|---|---|---|---|---|
| RHOBTB3 | 9606.ENSP00000369318 | GO Component | GO:0043226 | Organelle |
| RHOBTB3 | 9606.ENSP00000369318 | GO Component | GO:0043227 | Membrane-bounded organelle |
| RHOBTB3 | 9606.ENSP00000369318 | GO Component | GO:0043228 | Non-membrane-bounded organelle |
| RHOBTB3 | 9606.ENSP00000369318 | GO Component | GO:0043229 | Intracellular organelle |
| RHOBTB3 | 9606.ENSP00000369318 | GO Component | GO:0043230 | Extracellular organelle |
| RHOBTB3 | 9606.ENSP00000369318 | GO Component | GO:0043231 | Intracellular membrane-bounded organelle |
| RHOBTB3 | 9606.ENSP00000369318 | GO Component | GO:0043232 | Intracellular non-membrane-bounded organelle |
| RHOBTB3 | 9606.ENSP00000369318 | GO Component | GO:0065010 | Extracellular membrane-bounded organelle |
| RHOBTB3 | 9606.ENSP00000369318 | GO Component | GO:0070062 | Extracellular exosome |
| RHOBTB3 | 9606.ENSP00000369318 | GO Component | GO:0071944 | Cell periphery |
| RHOBTB3 | 9606.ENSP00000369318 | GO Component | GO:0097708 | Intracellular vesicle |
| RHOBTB3 | 9606.ENSP00000369318 | GO Component | GO:0098791 | Golgi apparatus subcompartment |
| RHOBTB3 | 9606.ENSP00000369318 | GO Component | GO:0110165 | Cellular anatomical entity |
| RHOBTB3 | 9606.ENSP00000369318 | GO Component | GO:1903561 | Extracellular vesicle |
| RHOBTB3 | 9606.ENSP00000369318 | STRING clusters | CL:13437 | Mixed, incl. RAB geranylgeranylation, and Vesicle tethering complex |
| RHOBTB3 | 9606.ENSP00000369318 | STRING clusters | CL:13649 | Mixed, incl. N-terminal extension of GAT domain, and RHOBTB3 ATPase cycle |
| RHOBTB3 | 9606.ENSP00000369318 | STRING clusters | CL:13662 | Rab9, and Kelch repeat type 2 |
| RHOBTB3 | 9606.ENSP00000369318 | Reactome | HSA-162582 | Signal Transduction |
| RHOBTB3 | 9606.ENSP00000369318 | Reactome | HSA-199991 | Membrane Trafficking |
| RHOBTB3 | 9606.ENSP00000369318 | Reactome | HSA-5653656 | Vesicle-mediated transport |
| RHOBTB3 | 9606.ENSP00000369318 | Reactome | HSA-6811440 | Retrograde transport at the Trans-Golgi-Network |
| RHOBTB3 | 9606.ENSP00000369318 | Reactome | HSA-6811442 | Intra-Golgi and retrograde Golgi-to-ER traffic |
| RHOBTB3 | 9606.ENSP00000369318 | Reactome | HSA-9706019 | RHOBTB3 ATPase cycle |
| RHOBTB3 | 9606.ENSP00000369318 | Reactome | HSA-9716542 | Signaling by Rho GTPases, Miro GTPases and RHOBTB3 |
| RHOBTB3 | 9606.ENSP00000369318 | TISSUES | BTO:0000000 | Tissues, cell types and enzyme sources |
| RHOBTB3 | 9606.ENSP00000369318 | TISSUES | BTO:0000042 | Animal |
| RHOBTB3 | 9606.ENSP00000369318 | TISSUES | BTO:0000081 | Reproductive system |
| RHOBTB3 | 9606.ENSP00000369318 | TISSUES | BTO:0000083 | Female reproductive system |
| RHOBTB3 | 9606.ENSP00000369318 | TISSUES | BTO:0000142 | Brain |
| RHOBTB3 | 9606.ENSP00000369318 | TISSUES | BTO:0000174 | Embryonic structure |
| RHOBTB3 | 9606.ENSP00000369318 | TISSUES | BTO:0000227 | Central nervous system |
| RHOBTB3 | 9606.ENSP00000369318 | TISSUES | BTO:0000231 | Cerebral hemisphere |
| RHOBTB3 | 9606.ENSP00000369318 | TISSUES | BTO:0000233 | Cerebral cortex |
| RHOBTB3 | 9606.ENSP00000369318 | TISSUES | BTO:0000239 | Telencephalon |

| | | | | |
|---|---|---|---|---|
| RHOBTB3 | 9606.ENSP00000369318 | TISSUES | BTO:0000282 | Head |
| RHOBTB3 | 9606.ENSP00000369318 | TISSUES | BTO:0000284 | Organism form |
| RHOBTB3 | 9606.ENSP00000369318 | TISSUES | BTO:0000449 | Fetus |
| RHOBTB3 | 9606.ENSP00000369318 | TISSUES | BTO:0000478 | Forebrain |
| RHOBTB3 | 9606.ENSP00000369318 | TISSUES | BTO:0000887 | Muscle |
| RHOBTB3 | 9606.ENSP00000369318 | TISSUES | BTO:0000928 | Limbic system |
| RHOBTB3 | 9606.ENSP00000369318 | TISSUES | BTO:0001078 | Placenta |
| RHOBTB3 | 9606.ENSP00000369318 | TISSUES | BTO:0001484 | Nervous system |
| RHOBTB3 | 9606.ENSP00000369318 | TISSUES | BTO:0001485 | Muscular system |
| RHOBTB3 | 9606.ENSP00000369318 | TISSUES | BTO:0001489 | Whole body |
| RHOBTB3 | 9606.ENSP00000369318 | TISSUES | BTO:0003091 | Urogenital system |
| RHOBTB3 | 9606.ENSP00000369318 | TISSUES | BTO:0003099 | Internal female genital organ |
| RHOBTB3 | 9606.ENSP00000369318 | COMPARTMENTS | GOCC:0000139 | Golgi membrane |
| RHOBTB3 | 9606.ENSP00000369318 | COMPARTMENTS | GOCC:0005622 | Intracellular |
| RHOBTB3 | 9606.ENSP00000369318 | COMPARTMENTS | GOCC:0005737 | Cytoplasm |
| RHOBTB3 | 9606.ENSP00000369318 | COMPARTMENTS | GOCC:0005794 | Golgi apparatus |
| RHOBTB3 | 9606.ENSP00000369318 | COMPARTMENTS | GOCC:0005802 | trans-Golgi network |
| RHOBTB3 | 9606.ENSP00000369318 | COMPARTMENTS | GOCC:0012505 | Endomembrane system |
| RHOBTB3 | 9606.ENSP00000369318 | COMPARTMENTS | GOCC:0016020 | Membrane |
| RHOBTB3 | 9606.ENSP00000369318 | COMPARTMENTS | GOCC:0031090 | Organelle membrane |
| RHOBTB3 | 9606.ENSP00000369318 | COMPARTMENTS | GOCC:0031984 | Organelle subcompartment |
| RHOBTB3 | 9606.ENSP00000369318 | COMPARTMENTS | GOCC:0032588 | trans-Golgi network membrane |
| RHOBTB3 | 9606.ENSP00000369318 | COMPARTMENTS | GOCC:0043226 | Organelle |
| RHOBTB3 | 9606.ENSP00000369318 | COMPARTMENTS | GOCC:0043227 | Membrane-bounded organelle |
| RHOBTB3 | 9606.ENSP00000369318 | COMPARTMENTS | GOCC:0043229 | Intracellular organelle |
| RHOBTB3 | 9606.ENSP00000369318 | COMPARTMENTS | GOCC:0043231 | Intracellular membrane-bounded organelle |
| RHOBTB3 | 9606.ENSP00000369318 | COMPARTMENTS | GOCC:0098588 | Bounding membrane of organelle |
| RHOBTB3 | 9606.ENSP00000369318 | COMPARTMENTS | GOCC:0098791 | Golgi apparatus subcompartment |
| RHOBTB3 | 9606.ENSP00000369318 | COMPARTMENTS | GOCC:0110165 | Cellular anatomical entity |
| RHOBTB3 | 9606.ENSP00000369318 | Monarch | EFO:0005134 | Amino acid measurement |
| RHOBTB3 | 9606.ENSP00000369318 | Monarch | EFO:0010487 | Glutamate measurement |
| RHOBTB3 | 9606.ENSP00000369318 | UniProt Keywords | KW-0067 | ATP-binding |
| RHOBTB3 | 9606.ENSP00000369318 | UniProt Keywords | KW-0333 | Golgi apparatus |
| RHOBTB3 | 9606.ENSP00000369318 | UniProt Keywords | KW-0378 | Hydrolase |
| RHOBTB3 | 9606.ENSP00000369318 | UniProt Keywords | KW-0547 | Nucleotide-binding |

| | | | | |
|---|---|---|---|---|
| RHOBTB3 | 9606.ENSP00000369318 | UniProt Keywords | KW-0677 | Repeat |
| RHOBTB3 | 9606.ENSP00000369318 | UniProt Keywords | KW-0813 | Transport |
| RHOBTB3 | 9606.ENSP00000369318 | Pfam | PF00071 | Ras family |
| RHOBTB3 | 9606.ENSP00000369318 | InterPro | IPR000210 | BTB/POZ domain |
| RHOBTB3 | 9606.ENSP00000369318 | InterPro | IPR001806 | Small GTPase |
| RHOBTB3 | 9606.ENSP00000369318 | InterPro | IPR011333 | SKP1/BTB/POZ domain superfamily |
| RHOBTB3 | 9606.ENSP00000369318 | InterPro | IPR027417 | P-loop containing nucleoside triphosphate hydrolase |
| RHOBTB3 | 9606.ENSP00000369318 | SMART | SM00225 | Broad-Complex, Tramtrack and Bric a brac |
| RPL28 | 9606.ENSP00000452763 | GO Process | GO:0002181 | Cytoplasmic translation |
| RPL28 | 9606.ENSP00000452763 | GO Process | GO:0006412 | Translation |
| RPL28 | 9606.ENSP00000452763 | GO Process | GO:0006518 | Peptide metabolic process |
| RPL28 | 9606.ENSP00000452763 | GO Process | GO:0006807 | Nitrogen compound metabolic process |
| RPL28 | 9606.ENSP00000452763 | GO Process | GO:0008152 | Metabolic process |
| RPL28 | 9606.ENSP00000452763 | GO Process | GO:0009058 | Biosynthetic process |
| RPL28 | 9606.ENSP00000452763 | GO Process | GO:0009059 | Macromolecule biosynthetic process |
| RPL28 | 9606.ENSP00000452763 | GO Process | GO:0009987 | Cellular process |
| RPL28 | 9606.ENSP00000452763 | GO Process | GO:0010467 | Gene expression |
| RPL28 | 9606.ENSP00000452763 | GO Process | GO:0019538 | Protein metabolic process |
| RPL28 | 9606.ENSP00000452763 | GO Process | GO:0034641 | Cellular nitrogen compound metabolic process |
| RPL28 | 9606.ENSP00000452763 | GO Process | GO:0034645 | Cellular macromolecule biosynthetic process |
| RPL28 | 9606.ENSP00000452763 | GO Process | GO:0043043 | Peptide biosynthetic process |
| RPL28 | 9606.ENSP00000452763 | GO Process | GO:0043170 | Macromolecule metabolic process |
| RPL28 | 9606.ENSP00000452763 | GO Process | GO:0043603 | Cellular amide metabolic process |
| RPL28 | 9606.ENSP00000452763 | GO Process | GO:0043604 | Amide biosynthetic process |
| RPL28 | 9606.ENSP00000452763 | GO Process | GO:0044237 | Cellular metabolic process |
| RPL28 | 9606.ENSP00000452763 | GO Process | GO:0044238 | Primary metabolic process |
| RPL28 | 9606.ENSP00000452763 | GO Process | GO:0044249 | Cellular biosynthetic process |
| RPL28 | 9606.ENSP00000452763 | GO Process | GO:0044260 | Cellular macromolecule metabolic process |
| RPL28 | 9606.ENSP00000452763 | GO Process | GO:0044271 | Cellular nitrogen compound biosynthetic process |
| RPL28 | 9606.ENSP00000452763 | GO Process | GO:0071704 | Organic substance metabolic process |
| RPL28 | 9606.ENSP00000452763 | GO Process | GO:1901564 | Organonitrogen compound metabolic process |
| RPL28 | 9606.ENSP00000452763 | GO Process | GO:1901566 | Organonitrogen compound biosynthetic process |
| RPL28 | 9606.ENSP00000452763 | GO Process | GO:1901576 | Organic substance biosynthetic process |
| RPL28 | 9606.ENSP00000452763 | GO Function | GO:0003676 | Nucleic acid binding |
| RPL28 | 9606.ENSP00000452763 | GO Function | GO:0003723 | RNA binding |

| | | | | |
|---|---|---|---|---|
| RPL28 | 9606.ENSP00000452763 | GO Function | GO:0003735 | Structural constituent of ribosome |
| RPL28 | 9606.ENSP00000452763 | GO Function | GO:0005198 | Structural molecule activity |
| RPL28 | 9606.ENSP00000452763 | GO Function | GO:0005488 | Binding |
| RPL28 | 9606.ENSP00000452763 | GO Function | GO:0097159 | Organic cyclic compound binding |
| RPL28 | 9606.ENSP00000452763 | GO Function | GO:1901363 | Heterocyclic compound binding |
| RPL28 | 9606.ENSP00000452763 | GO Component | GO:0005576 | Extracellular region |
| RPL28 | 9606.ENSP00000452763 | GO Component | GO:0005615 | Extracellular space |
| RPL28 | 9606.ENSP00000452763 | GO Component | GO:0005622 | Intracellular anatomical structure |
| RPL28 | 9606.ENSP00000452763 | GO Component | GO:0005737 | Cytoplasm |
| RPL28 | 9606.ENSP00000452763 | GO Component | GO:0005829 | Cytosol |
| RPL28 | 9606.ENSP00000452763 | GO Component | GO:0005840 | Ribosome |
| RPL28 | 9606.ENSP00000452763 | GO Component | GO:0015934 | Large ribosomal subunit |
| RPL28 | 9606.ENSP00000452763 | GO Component | GO:0016020 | Membrane |
| RPL28 | 9606.ENSP00000452763 | GO Component | GO:0022625 | Cytosolic large ribosomal subunit |
| RPL28 | 9606.ENSP00000452763 | GO Component | GO:0022626 | Cytosolic ribosome |
| RPL28 | 9606.ENSP00000452763 | GO Component | GO:0030425 | Dendrite |
| RPL28 | 9606.ENSP00000452763 | GO Component | GO:0031982 | Vesicle |
| RPL28 | 9606.ENSP00000452763 | GO Component | GO:0032991 | Protein-containing complex |
| RPL28 | 9606.ENSP00000452763 | GO Component | GO:0035770 | Ribonucleoprotein granule |
| RPL28 | 9606.ENSP00000452763 | GO Component | GO:0036464 | Cytoplasmic ribonucleoprotein granule |
| RPL28 | 9606.ENSP00000452763 | GO Component | GO:0036477 | Somatodendritic compartment |
| RPL28 | 9606.ENSP00000452763 | GO Component | GO:0042995 | Cell projection |
| RPL28 | 9606.ENSP00000452763 | GO Component | GO:0043005 | Neuron projection |
| RPL28 | 9606.ENSP00000452763 | GO Component | GO:0043226 | Organelle |
| RPL28 | 9606.ENSP00000452763 | GO Component | GO:0043227 | Membrane-bounded organelle |
| RPL28 | 9606.ENSP00000452763 | GO Component | GO:0043228 | Non-membrane-bounded organelle |
| RPL28 | 9606.ENSP00000452763 | GO Component | GO:0043229 | Intracellular organelle |
| RPL28 | 9606.ENSP00000452763 | GO Component | GO:0043230 | Extracellular organelle |
| RPL28 | 9606.ENSP00000452763 | GO Component | GO:0043232 | Intracellular non-membrane-bounded organelle |
| RPL28 | 9606.ENSP00000452763 | GO Component | GO:0044297 | Cell body |
| RPL28 | 9606.ENSP00000452763 | GO Component | GO:0044391 | Ribosomal subunit |
| RPL28 | 9606.ENSP00000452763 | GO Component | GO:0065010 | Extracellular membrane-bounded organelle |
| RPL28 | 9606.ENSP00000452763 | GO Component | GO:0070062 | Extracellular exosome |
| RPL28 | 9606.ENSP00000452763 | GO Component | GO:0097447 | Dendritic tree |
| RPL28 | 9606.ENSP00000452763 | GO Component | GO:0099080 | Supramolecular complex |

| | | | | |
|---|---|---|---|---|
| RPL28 | 9606.ENSP00000452763 | GO Component | GO:0110165 | Cellular anatomical entity |
| RPL28 | 9606.ENSP00000452763 | GO Component | GO:0120025 | Plasma membrane bounded cell projection |
| RPL28 | 9606.ENSP00000452763 | GO Component | GO:1903561 | Extracellular vesicle |
| RPL28 | 9606.ENSP00000452763 | GO Component | GO:1990904 | Ribonucleoprotein complex |
| RPL28 | 9606.ENSP00000452763 | STRING clusters | CL:112 | Mixed, incl. Eukaryotic Translation Elongation, and This family consists of several GAGE and XAGE proteins which are found exclusively in humans. The function of this family is unknown although they have been implicated in human cancers (PMID:11992404) |
| RPL28 | 9606.ENSP00000452763 | STRING clusters | CL:114 | Mixed, incl. Eukaryotic Translation Elongation, and This family consists of several GAGE and XAGE proteins which are found exclusively in humans. The function of this family is unknown although they have been implicated in human cancers (PMID:11992404) |
| RPL28 | 9606.ENSP00000452763 | STRING clusters | CL:118 | Eukaryotic Translation Elongation, and This family consists of several GAGE and XAGE proteins which are found exclusively in humans. The function of this family is unknown although they have been implicated in human cancers (PMID:11992404) |
| RPL28 | 9606.ENSP00000452763 | STRING clusters | CL:121 | Eukaryotic Translation Elongation, and This family consists of several GAGE and XAGE proteins which are found exclusively in humans. The function of this family is unknown although they have been implicated in human cancers (PMID:11992404) |
| RPL28 | 9606.ENSP00000452763 | STRING clusters | CL:125 | Eukaryotic Translation Elongation, and This family consists of several GAGE and XAGE proteins which are found exclusively in humans. The function of this family is unknown although they have been implicated in human cancers (PMID:11992404) |
| RPL28 | 9606.ENSP00000452763 | STRING clusters | CL:130 | Eukaryotic Translation Elongation, and This family consists of several GAGE and XAGE proteins which are found exclusively in humans. The function of this family is unknown although they have been implicated in human cancers (PMID:11992404) |
| RPL28 | 9606.ENSP00000452763 | STRING clusters | CL:133 | Eukaryotic Translation Elongation, and Positive regulation of cyclin-dependent protein serine/threonine kinase activity |
| RPL28 | 9606.ENSP00000452763 | STRING clusters | CL:135 | Eukaryotic Translation Elongation, and Positive regulation of cyclin-dependent protein serine/threonine kinase activity |
| RPL28 | 9606.ENSP00000452763 | STRING clusters | CL:140 | Eukaryotic Translation Elongation, and Sec61 translocon complex |
| RPL28 | 9606.ENSP00000452763 | STRING clusters | CL:143 | Viral mRNA Translation, and Sec61 translocon complex |
| RPL28 | 9606.ENSP00000452763 | STRING clusters | CL:148 | Viral mRNA Translation |

| | | | | |
|---|---|---|---|---|
| RPL28 | 9606.ENSP00000452763 | STRING clusters | CL:152 | Viral mRNA Translation |
| RPL28 | 9606.ENSP00000452763 | STRING clusters | CL:156 | Viral mRNA Translation |
| RPL28 | 9606.ENSP00000452763 | STRING clusters | CL:159 | Viral mRNA Translation |
| RPL28 | 9606.ENSP00000452763 | STRING clusters | CL:162 | Cytoplasmic ribosomal proteins |
| RPL28 | 9606.ENSP00000452763 | STRING clusters | CL:165 | Cytoplasmic ribosomal proteins |
| RPL28 | 9606.ENSP00000452763 | STRING clusters | CL:166 | Cytoplasmic ribosomal proteins |
| RPL28 | 9606.ENSP00000452763 | STRING clusters | CL:167 | Cytoplasmic ribosomal proteins |
| RPL28 | 9606.ENSP00000452763 | STRING clusters | CL:168 | Cytoplasmic ribosomal proteins |
| RPL28 | 9606.ENSP00000452763 | STRING clusters | CL:169 | Cytoplasmic ribosomal proteins |
| RPL28 | 9606.ENSP00000452763 | STRING clusters | CL:170 | Cytoplasmic ribosomal proteins |
| RPL28 | 9606.ENSP00000452763 | KEGG | hsa03010 | Ribosome |
| RPL28 | 9606.ENSP00000452763 | Reactome | HSA-1266738 | Developmental Biology |
| RPL28 | 9606.ENSP00000452763 | Reactome | HSA-1430728 | Metabolism |
| RPL28 | 9606.ENSP00000452763 | Reactome | HSA-156827 | L13a-mediated translational silencing of Ceruloplasmin expression |
| RPL28 | 9606.ENSP00000452763 | Reactome | HSA-156842 | Eukaryotic Translation Elongation |
| RPL28 | 9606.ENSP00000452763 | Reactome | HSA-156902 | Peptide chain elongation |
| RPL28 | 9606.ENSP00000452763 | Reactome | HSA-1643685 | Disease |
| RPL28 | 9606.ENSP00000452763 | Reactome | HSA-168255 | Influenza Infection |
| RPL28 | 9606.ENSP00000452763 | Reactome | HSA-168273 | Influenza Viral RNA Transcription and Replication |
| RPL28 | 9606.ENSP00000452763 | Reactome | HSA-1799339 | SRP-dependent cotranslational protein targeting to membrane |
| RPL28 | 9606.ENSP00000452763 | Reactome | HSA-192823 | Viral mRNA Translation |
| RPL28 | 9606.ENSP00000452763 | Reactome | HSA-2262752 | Cellular responses to stress |
| RPL28 | 9606.ENSP00000452763 | Reactome | HSA-2408522 | Selenoamino acid metabolism |
| RPL28 | 9606.ENSP00000452763 | Reactome | HSA-2408557 | Selenocysteine synthesis |
| RPL28 | 9606.ENSP00000452763 | Reactome | HSA-376176 | Signaling by ROBO receptors |
| RPL28 | 9606.ENSP00000452763 | Reactome | HSA-392499 | Metabolism of proteins |
| RPL28 | 9606.ENSP00000452763 | Reactome | HSA-422475 | Axon guidance |
| RPL28 | 9606.ENSP00000452763 | Reactome | HSA-5663205 | Infectious disease |
| RPL28 | 9606.ENSP00000452763 | Reactome | HSA-6791226 | Major pathway of rRNA processing in the nucleolus and cytosol |
| RPL28 | 9606.ENSP00000452763 | Reactome | HSA-71291 | Metabolism of amino acids and derivatives |
| RPL28 | 9606.ENSP00000452763 | Reactome | HSA-72312 | rRNA processing |
| RPL28 | 9606.ENSP00000452763 | Reactome | HSA-72613 | Eukaryotic Translation Initiation |
| RPL28 | 9606.ENSP00000452763 | Reactome | HSA-72689 | Formation of a pool of free 40S subunits |
| RPL28 | 9606.ENSP00000452763 | Reactome | HSA-72706 | GTP hydrolysis and joining of the 60S ribosomal subunit |
| RPL28 | 9606.ENSP00000452763 | Reactome | HSA-72737 | Cap-dependent Translation Initiation |

| | | | | |
|---|---|---|---|---|
| RPL28 | 9606.ENSP00000452763 | Reactome | HSA-72764 | Eukaryotic Translation Termination |
| RPL28 | 9606.ENSP00000452763 | Reactome | HSA-72766 | Translation |
| RPL28 | 9606.ENSP00000452763 | Reactome | HSA-8868773 | rRNA processing in the nucleus and cytosol |
| RPL28 | 9606.ENSP00000452763 | Reactome | HSA-8953854 | Metabolism of RNA |
| RPL28 | 9606.ENSP00000452763 | Reactome | HSA-8953897 | Cellular responses to stimuli |
| RPL28 | 9606.ENSP00000452763 | Reactome | HSA-9010553 | Regulation of expression of SLITs and ROBOs |
| RPL28 | 9606.ENSP00000452763 | Reactome | HSA-927802 | Nonsense-Mediated Decay (NMD) |
| RPL28 | 9606.ENSP00000452763 | Reactome | HSA-9633012 | Response of EIF2AK4 (GCN2) to amino acid deficiency |
| RPL28 | 9606.ENSP00000452763 | Reactome | HSA-9675108 | Nervous system development |
| RPL28 | 9606.ENSP00000452763 | Reactome | HSA-9711097 | Cellular response to starvation |
| RPL28 | 9606.ENSP00000452763 | Reactome | HSA-975956 | Nonsense Mediated Decay (NMD) independent of the Exon Junction Complex (EJC) |
| RPL28 | 9606.ENSP00000452763 | Reactome | HSA-975957 | Nonsense Mediated Decay (NMD) enhanced by the Exon Junction Complex (EJC) |
| RPL28 | 9606.ENSP00000452763 | WikiPathways | WP477 | Cytoplasmic ribosomal proteins |
| RPL28 | 9606.ENSP00000452763 | TISSUES | BTO:0000000 | Tissues, cell types and enzyme sources |
| RPL28 | 9606.ENSP00000452763 | TISSUES | BTO:0000042 | Animal |
| RPL28 | 9606.ENSP00000452763 | TISSUES | BTO:0000058 | Alimentary canal |
| RPL28 | 9606.ENSP00000452763 | TISSUES | BTO:0000080 | Male reproductive gland |
| RPL28 | 9606.ENSP00000452763 | TISSUES | BTO:0000081 | Reproductive system |
| RPL28 | 9606.ENSP00000452763 | TISSUES | BTO:0000082 | Male reproductive system |
| RPL28 | 9606.ENSP00000452763 | TISSUES | BTO:0000083 | Female reproductive system |
| RPL28 | 9606.ENSP00000452763 | TISSUES | BTO:0000142 | Brain |
| RPL28 | 9606.ENSP00000452763 | TISSUES | BTO:0000146 | Brain stem |
| RPL28 | 9606.ENSP00000452763 | TISSUES | BTO:0000176 | Carcinoma cell |
| RPL28 | 9606.ENSP00000452763 | TISSUES | BTO:0000227 | Central nervous system |
| RPL28 | 9606.ENSP00000452763 | TISSUES | BTO:0000231 | Cerebral hemisphere |
| RPL28 | 9606.ENSP00000452763 | TISSUES | BTO:0000232 | Cerebellum |
| RPL28 | 9606.ENSP00000452763 | TISSUES | BTO:0000233 | Cerebral cortex |
| RPL28 | 9606.ENSP00000452763 | TISSUES | BTO:0000239 | Telencephalon |
| RPL28 | 9606.ENSP00000452763 | TISSUES | BTO:0000254 | Female reproductive gland |
| RPL28 | 9606.ENSP00000452763 | TISSUES | BTO:0000269 | Colon |
| RPL28 | 9606.ENSP00000452763 | TISSUES | BTO:0000282 | Head |
| RPL28 | 9606.ENSP00000452763 | TISSUES | BTO:0000345 | Digestive gland |
| RPL28 | 9606.ENSP00000452763 | TISSUES | BTO:0000415 | Epithelioma cell |

| | | | | |
|---|---|---|---|---|
| RPL28 | 9606.ENSP00000452763 | TISSUES | BTO:0000431 | Excretory gland |
| RPL28 | 9606.ENSP00000452763 | TISSUES | BTO:0000445 | Cerebral lobe |
| RPL28 | 9606.ENSP00000452763 | TISSUES | BTO:0000478 | Forebrain |
| RPL28 | 9606.ENSP00000452763 | TISSUES | BTO:0000511 | Gastrointestinal tract |
| RPL28 | 9606.ENSP00000452763 | TISSUES | BTO:0000522 | Gland |
| RPL28 | 9606.ENSP00000452763 | TISSUES | BTO:0000594 | Liver cancer cell |
| RPL28 | 9606.ENSP00000452763 | TISSUES | BTO:0000601 | Hippocampus |
| RPL28 | 9606.ENSP00000452763 | TISSUES | BTO:0000604 | Adenocarcinoma cell |
| RPL28 | 9606.ENSP00000452763 | TISSUES | BTO:0000608 | Hepatoma cell |
| RPL28 | 9606.ENSP00000452763 | TISSUES | BTO:0000634 | Integument |
| RPL28 | 9606.ENSP00000452763 | TISSUES | BTO:0000648 | Intestine |
| RPL28 | 9606.ENSP00000452763 | TISSUES | BTO:0000671 | Kidney |
| RPL28 | 9606.ENSP00000452763 | TISSUES | BTO:0000672 | Hindbrain |
| RPL28 | 9606.ENSP00000452763 | TISSUES | BTO:0000673 | Metencephalon |
| RPL28 | 9606.ENSP00000452763 | TISSUES | BTO:0000706 | Large intestine |
| RPL28 | 9606.ENSP00000452763 | TISSUES | BTO:0000928 | Limbic system |
| RPL28 | 9606.ENSP00000452763 | TISSUES | BTO:0000975 | Ovary |
| RPL28 | 9606.ENSP00000452763 | TISSUES | BTO:0000988 | Pancreas |
| RPL28 | 9606.ENSP00000452763 | TISSUES | BTO:0001244 | Urinary tract |
| RPL28 | 9606.ENSP00000452763 | TISSUES | BTO:0001253 | Skin |
| RPL28 | 9606.ENSP00000452763 | TISSUES | BTO:0001355 | Temporal lobe |
| RPL28 | 9606.ENSP00000452763 | TISSUES | BTO:0001424 | Uterus |
| RPL28 | 9606.ENSP00000452763 | TISSUES | BTO:0001484 | Nervous system |
| RPL28 | 9606.ENSP00000452763 | TISSUES | BTO:0001488 | Endocrine gland |
| RPL28 | 9606.ENSP00000452763 | TISSUES | BTO:0001489 | Whole body |
| RPL28 | 9606.ENSP00000452763 | TISSUES | BTO:0001491 | Viscus |
| RPL28 | 9606.ENSP00000452763 | TISSUES | BTO:0001613 | Colorectum |
| RPL28 | 9606.ENSP00000452763 | TISSUES | BTO:0002021 | Prostate adenocarcinoma cell |
| RPL28 | 9606.ENSP00000452763 | TISSUES | BTO:0003091 | Urogenital system |
| RPL28 | 9606.ENSP00000452763 | TISSUES | BTO:0003092 | Urinary system |
| RPL28 | 9606.ENSP00000452763 | TISSUES | BTO:0003099 | Internal female genital organ |
| RPL28 | 9606.ENSP00000452763 | COMPARTMENTS | GOCC:0005622 | Intracellular |
| RPL28 | 9606.ENSP00000452763 | COMPARTMENTS | GOCC:0005737 | Cytoplasm |
| RPL28 | 9606.ENSP00000452763 | COMPARTMENTS | GOCC:0005829 | Cytosol |
| RPL28 | 9606.ENSP00000452763 | COMPARTMENTS | GOCC:0005840 | Ribosome |

| | | | | |
|---|---|---|---|---|
| RPL28 | 9606.ENSP00000452763 | COMPARTMENTS | GOCC:0022626 | Cytosolic ribosome |
| RPL28 | 9606.ENSP00000452763 | COMPARTMENTS | GOCC:0035770 | Ribonucleoprotein granule |
| RPL28 | 9606.ENSP00000452763 | COMPARTMENTS | GOCC:0036464 | Cytoplasmic ribonucleoprotein granule |
| RPL28 | 9606.ENSP00000452763 | COMPARTMENTS | GOCC:0043226 | Organelle |
| RPL28 | 9606.ENSP00000452763 | COMPARTMENTS | GOCC:0043228 | Non-membrane-bounded organelle |
| RPL28 | 9606.ENSP00000452763 | COMPARTMENTS | GOCC:0043229 | Intracellular organelle |
| RPL28 | 9606.ENSP00000452763 | COMPARTMENTS | GOCC:0043232 | Intracellular non-membrane-bounded organelle |
| RPL28 | 9606.ENSP00000452763 | COMPARTMENTS | GOCC:0099080 | Supramolecular complex |
| RPL28 | 9606.ENSP00000452763 | COMPARTMENTS | GOCC:0110165 | Cellular anatomical entity |
| RPL28 | 9606.ENSP00000452763 | Monarch | EFO:0004464 | Brain measurement |
| RPL28 | 9606.ENSP00000452763 | Monarch | EFO:0005674 | White matter microstructure measurement |
| RPL28 | 9606.ENSP00000452763 | UniProt Keywords | KW-0007 | Acetylation |
| RPL28 | 9606.ENSP00000452763 | UniProt Keywords | KW-0025 | Alternative splicing |
| RPL28 | 9606.ENSP00000452763 | UniProt Keywords | KW-0597 | Phosphoprotein |
| RPL28 | 9606.ENSP00000452763 | UniProt Keywords | KW-0687 | Ribonucleoprotein |
| RPL28 | 9606.ENSP00000452763 | UniProt Keywords | KW-0689 | Ribosomal protein |
| RPL28 | 9606.ENSP00000452763 | UniProt Keywords | KW-0832 | Ubl conjugation |
| RPL28 | 9606.ENSP00000452763 | UniProt Keywords | KW-1017 | Isopeptide bond |
| RPL28 | 9606.ENSP00000452763 | Pfam | PF01778 | Ribosomal L28e protein family |
| RPL28 | 9606.ENSP00000452763 | InterPro | IPR002672 | Ribosomal protein L28e |
| RPL28 | 9606.ENSP00000452763 | InterPro | IPR029004 | Ribosomal L28e/Mak16 |
| RPP30 | 9606.ENSP00000389182 | GO Process | GO:0000966 | RNA 5-end processing |
| RPP30 | 9606.ENSP00000389182 | GO Process | GO:0001682 | tRNA 5-leader removal |
| RPP30 | 9606.ENSP00000389182 | GO Process | GO:0006139 | Nucleobase-containing compound metabolic process |
| RPP30 | 9606.ENSP00000389182 | GO Process | GO:0006364 | rRNA processing |
| RPP30 | 9606.ENSP00000389182 | GO Process | GO:0006396 | RNA processing |
| RPP30 | 9606.ENSP00000389182 | GO Process | GO:0006399 | tRNA metabolic process |
| RPP30 | 9606.ENSP00000389182 | GO Process | GO:0006725 | Cellular aromatic compound metabolic process |
| RPP30 | 9606.ENSP00000389182 | GO Process | GO:0006807 | Nitrogen compound metabolic process |
| RPP30 | 9606.ENSP00000389182 | GO Process | GO:0008033 | tRNA processing |
| RPP30 | 9606.ENSP00000389182 | GO Process | GO:0008152 | Metabolic process |
| RPP30 | 9606.ENSP00000389182 | GO Process | GO:0009987 | Cellular process |
| RPP30 | 9606.ENSP00000389182 | GO Process | GO:0010467 | Gene expression |
| RPP30 | 9606.ENSP00000389182 | GO Process | GO:0016070 | RNA metabolic process |
| RPP30 | 9606.ENSP00000389182 | GO Process | GO:0016072 | rRNA metabolic process |

| | | | | |
|---|---|---|---|---|
| RPP30 | 9606.ENSP00000389182 | GO Process | GO:0022613 | Ribonucleoprotein complex biogenesis |
| RPP30 | 9606.ENSP00000389182 | GO Process | GO:0034470 | ncRNA processing |
| RPP30 | 9606.ENSP00000389182 | GO Process | GO:0034641 | Cellular nitrogen compound metabolic process |
| RPP30 | 9606.ENSP00000389182 | GO Process | GO:0034660 | ncRNA metabolic process |
| RPP30 | 9606.ENSP00000389182 | GO Process | GO:0042254 | Ribosome biogenesis |
| RPP30 | 9606.ENSP00000389182 | GO Process | GO:0043170 | Macromolecule metabolic process |
| RPP30 | 9606.ENSP00000389182 | GO Process | GO:0044085 | Cellular component biogenesis |
| RPP30 | 9606.ENSP00000389182 | GO Process | GO:0044237 | Cellular metabolic process |
| RPP30 | 9606.ENSP00000389182 | GO Process | GO:0044238 | Primary metabolic process |
| RPP30 | 9606.ENSP00000389182 | GO Process | GO:0046483 | Heterocycle metabolic process |
| RPP30 | 9606.ENSP00000389182 | GO Process | GO:0071704 | Organic substance metabolic process |
| RPP30 | 9606.ENSP00000389182 | GO Process | GO:0071840 | Cellular component organization or biogenesis |
| RPP30 | 9606.ENSP00000389182 | GO Process | GO:0090304 | Nucleic acid metabolic process |
| RPP30 | 9606.ENSP00000389182 | GO Process | GO:0090305 | Nucleic acid phosphodiester bond hydrolysis |
| RPP30 | 9606.ENSP00000389182 | GO Process | GO:0090501 | RNA phosphodiester bond hydrolysis |
| RPP30 | 9606.ENSP00000389182 | GO Process | GO:0090502 | RNA phosphodiester bond hydrolysis, endonucleolytic |
| RPP30 | 9606.ENSP00000389182 | GO Process | GO:0099116 | tRNA 5-end processing |
| RPP30 | 9606.ENSP00000389182 | GO Process | GO:1901360 | Organic cyclic compound metabolic process |
| RPP30 | 9606.ENSP00000389182 | GO Function | GO:0003676 | Nucleic acid binding |
| RPP30 | 9606.ENSP00000389182 | GO Function | GO:0003723 | RNA binding |
| RPP30 | 9606.ENSP00000389182 | GO Function | GO:0003824 | Catalytic activity |
| RPP30 | 9606.ENSP00000389182 | GO Function | GO:0004518 | Nuclease activity |
| RPP30 | 9606.ENSP00000389182 | GO Function | GO:0004519 | Endonuclease activity |
| RPP30 | 9606.ENSP00000389182 | GO Function | GO:0004521 | Endoribonuclease activity |
| RPP30 | 9606.ENSP00000389182 | GO Function | GO:0004526 | Ribonuclease P activity |
| RPP30 | 9606.ENSP00000389182 | GO Function | GO:0004540 | Ribonuclease activity |
| RPP30 | 9606.ENSP00000389182 | GO Function | GO:0004549 | tRNA-specific ribonuclease activity |
| RPP30 | 9606.ENSP00000389182 | GO Function | GO:0005488 | Binding |
| RPP30 | 9606.ENSP00000389182 | GO Function | GO:0016787 | Hydrolase activity |
| RPP30 | 9606.ENSP00000389182 | GO Function | GO:0016788 | Hydrolase activity, acting on ester bonds |
| RPP30 | 9606.ENSP00000389182 | GO Function | GO:0016891 | Endoribonuclease activity, producing 5-phosphomonoesters |
| RPP30 | 9606.ENSP00000389182 | GO Function | GO:0016893 | Endonuclease activity, active with either ribo- or deoxyribonucleic acids and producing 5-phosphomonoesters |
| RPP30 | 9606.ENSP00000389182 | GO Function | GO:0033204 | Ribonuclease P RNA binding |
| RPP30 | 9606.ENSP00000389182 | GO Function | GO:0097159 | Organic cyclic compound binding |

| | | | | |
|---|---|---|---|---|
| RPP30 | 9606.ENSP00000389182 | GO Function | GO:0140098 | Catalytic activity, acting on RNA |
| RPP30 | 9606.ENSP00000389182 | GO Function | GO:0140101 | Catalytic activity, acting on a tRNA |
| RPP30 | 9606.ENSP00000389182 | GO Function | GO:0140640 | Catalytic activity, acting on a nucleic acid |
| RPP30 | 9606.ENSP00000389182 | GO Function | GO:1901363 | Heterocyclic compound binding |
| RPP30 | 9606.ENSP00000389182 | GO Component | GO:0000172 | Ribonuclease MRP complex |
| RPP30 | 9606.ENSP00000389182 | GO Component | GO:0005622 | Intracellular anatomical structure |
| RPP30 | 9606.ENSP00000389182 | GO Component | GO:0005634 | Nucleus |
| RPP30 | 9606.ENSP00000389182 | GO Component | GO:0005654 | Nucleoplasm |
| RPP30 | 9606.ENSP00000389182 | GO Component | GO:0005655 | Nucleolar ribonuclease P complex |
| RPP30 | 9606.ENSP00000389182 | GO Component | GO:0005730 | Nucleolus |
| RPP30 | 9606.ENSP00000389182 | GO Component | GO:0005732 | sno(s)RNA-containing ribonucleoprotein complex |
| RPP30 | 9606.ENSP00000389182 | GO Component | GO:0030677 | Ribonuclease P complex |
| RPP30 | 9606.ENSP00000389182 | GO Component | GO:0030681 | Multimeric ribonuclease P complex |
| RPP30 | 9606.ENSP00000389182 | GO Component | GO:0031974 | Membrane-enclosed lumen |
| RPP30 | 9606.ENSP00000389182 | GO Component | GO:0031981 | Nuclear lumen |
| RPP30 | 9606.ENSP00000389182 | GO Component | GO:0032991 | Protein-containing complex |
| RPP30 | 9606.ENSP00000389182 | GO Component | GO:0043226 | Organelle |
| RPP30 | 9606.ENSP00000389182 | GO Component | GO:0043227 | Membrane-bounded organelle |
| RPP30 | 9606.ENSP00000389182 | GO Component | GO:0043228 | Non-membrane-bounded organelle |
| RPP30 | 9606.ENSP00000389182 | GO Component | GO:0043229 | Intracellular organelle |
| RPP30 | 9606.ENSP00000389182 | GO Component | GO:0043231 | Intracellular membrane-bounded organelle |
| RPP30 | 9606.ENSP00000389182 | GO Component | GO:0043232 | Intracellular non-membrane-bounded organelle |
| RPP30 | 9606.ENSP00000389182 | GO Component | GO:0043233 | Organelle lumen |
| RPP30 | 9606.ENSP00000389182 | GO Component | GO:0070013 | Intracellular organelle lumen |
| RPP30 | 9606.ENSP00000389182 | GO Component | GO:0110165 | Cellular anatomical entity |
| RPP30 | 9606.ENSP00000389182 | GO Component | GO:0140513 | Nuclear protein-containing complex |
| RPP30 | 9606.ENSP00000389182 | GO Component | GO:1902494 | Catalytic complex |
| RPP30 | 9606.ENSP00000389182 | GO Component | GO:1902555 | Endoribonuclease complex |
| RPP30 | 9606.ENSP00000389182 | GO Component | GO:1990904 | Ribonucleoprotein complex |
| RPP30 | 9606.ENSP00000389182 | STRING clusters | CL:2921 | Mixed, incl. Endoribonuclease complex, and Deadenylation-dependent mRNA decay |
| RPP30 | 9606.ENSP00000389182 | STRING clusters | CL:3082 | Endoribonuclease complex, and Argonaute hook |
| RPP30 | 9606.ENSP00000389182 | STRING clusters | CL:3116 | Ribonuclease P complex, and Anauxetic dysplasia 1 |
| RPP30 | 9606.ENSP00000389182 | STRING clusters | CL:3118 | Ribonuclease P RNA binding |
| RPP30 | 9606.ENSP00000389182 | STRING clusters | CL:3120 | Nucleolar ribonuclease P complex |

| | | | | |
|---|---|---|---|---|
| RPP30 | 9606.ENSP00000389182 | KEGG | hsa03008 | Ribosome biogenesis in eukaryotes |
| RPP30 | 9606.ENSP00000389182 | KEGG | hsa03013 | RNA transport |
| RPP30 | 9606.ENSP00000389182 | Reactome | HSA-6784531 | tRNA processing in the nucleus |
| RPP30 | 9606.ENSP00000389182 | Reactome | HSA-6791226 | Major pathway of rRNA processing in the nucleolus and cytosol |
| RPP30 | 9606.ENSP00000389182 | Reactome | HSA-72306 | tRNA processing |
| RPP30 | 9606.ENSP00000389182 | Reactome | HSA-72312 | rRNA processing |
| RPP30 | 9606.ENSP00000389182 | Reactome | HSA-8868773 | rRNA processing in the nucleus and cytosol |
| RPP30 | 9606.ENSP00000389182 | Reactome | HSA-8953854 | Metabolism of RNA |
| RPP30 | 9606.ENSP00000389182 | TISSUES | BTO:0000000 | Tissues, cell types and enzyme sources |
| RPP30 | 9606.ENSP00000389182 | TISSUES | BTO:0000042 | Animal |
| RPP30 | 9606.ENSP00000389182 | TISSUES | BTO:0000372 | Bone cancer cell |
| RPP30 | 9606.ENSP00000389182 | TISSUES | BTO:0000431 | Excretory gland |
| RPP30 | 9606.ENSP00000389182 | TISSUES | BTO:0000522 | Gland |
| RPP30 | 9606.ENSP00000389182 | TISSUES | BTO:0000671 | Kidney |
| RPP30 | 9606.ENSP00000389182 | TISSUES | BTO:0000970 | Osteosarcoma cell |
| RPP30 | 9606.ENSP00000389182 | TISSUES | BTO:0001207 | Sarcoma cell |
| RPP30 | 9606.ENSP00000389182 | TISSUES | BTO:0001244 | Urinary tract |
| RPP30 | 9606.ENSP00000389182 | TISSUES | BTO:0001489 | Whole body |
| RPP30 | 9606.ENSP00000389182 | TISSUES | BTO:0001491 | Viscus |
| RPP30 | 9606.ENSP00000389182 | TISSUES | BTO:0001801 | Skeletal muscle cancer cell |
| RPP30 | 9606.ENSP00000389182 | TISSUES | BTO:0003091 | Urogenital system |
| RPP30 | 9606.ENSP00000389182 | TISSUES | BTO:0003092 | Urinary system |
| RPP30 | 9606.ENSP00000389182 | COMPARTMENTS | GOCC:0000172 | Ribonuclease MRP complex |
| RPP30 | 9606.ENSP00000389182 | COMPARTMENTS | GOCC:0005622 | Intracellular |
| RPP30 | 9606.ENSP00000389182 | COMPARTMENTS | GOCC:0005634 | Nucleus |
| RPP30 | 9606.ENSP00000389182 | COMPARTMENTS | GOCC:0005655 | Nucleolar ribonuclease P complex |
| RPP30 | 9606.ENSP00000389182 | COMPARTMENTS | GOCC:0005730 | Nucleolus |
| RPP30 | 9606.ENSP00000389182 | COMPARTMENTS | GOCC:0005732 | Small nucleolar ribonucleoprotein complex |
| RPP30 | 9606.ENSP00000389182 | COMPARTMENTS | GOCC:0030677 | Ribonuclease P complex |
| RPP30 | 9606.ENSP00000389182 | COMPARTMENTS | GOCC:0030681 | Multimeric ribonuclease P complex |
| RPP30 | 9606.ENSP00000389182 | COMPARTMENTS | GOCC:0031974 | Membrane-enclosed lumen |
| RPP30 | 9606.ENSP00000389182 | COMPARTMENTS | GOCC:0031981 | Nuclear lumen |
| RPP30 | 9606.ENSP00000389182 | COMPARTMENTS | GOCC:0032991 | Protein-containing complex |
| RPP30 | 9606.ENSP00000389182 | COMPARTMENTS | GOCC:0043226 | Organelle |
| RPP30 | 9606.ENSP00000389182 | COMPARTMENTS | GOCC:0043227 | Membrane-bounded organelle |

| | | | | |
|---|---|---|---|---|
| RPP30 | 9606.ENSP00000389182 | COMPARTMENTS | GOCC:0043228 | Non-membrane-bounded organelle |
| RPP30 | 9606.ENSP00000389182 | COMPARTMENTS | GOCC:0043229 | Intracellular organelle |
| RPP30 | 9606.ENSP00000389182 | COMPARTMENTS | GOCC:0043231 | Intracellular membrane-bounded organelle |
| RPP30 | 9606.ENSP00000389182 | COMPARTMENTS | GOCC:0043232 | Intracellular non-membrane-bounded organelle |
| RPP30 | 9606.ENSP00000389182 | COMPARTMENTS | GOCC:0043233 | Organelle lumen |
| RPP30 | 9606.ENSP00000389182 | COMPARTMENTS | GOCC:0070013 | Intracellular organelle lumen |
| RPP30 | 9606.ENSP00000389182 | COMPARTMENTS | GOCC:0110165 | Cellular anatomical entity |
| RPP30 | 9606.ENSP00000389182 | COMPARTMENTS | GOCC:1902494 | Catalytic complex |
| RPP30 | 9606.ENSP00000389182 | COMPARTMENTS | GOCC:1902555 | Endoribonuclease complex |
| RPP30 | 9606.ENSP00000389182 | COMPARTMENTS | GOCC:1905348 | Endonuclease complex |
| RPP30 | 9606.ENSP00000389182 | COMPARTMENTS | GOCC:1990904 | Ribonucleoprotein complex |
| RPP30 | 9606.ENSP00000389182 | Monarch | EFO:0004827 | Economic and social preference |
| RPP30 | 9606.ENSP00000389182 | UniProt Keywords | KW-0007 | Acetylation |
| RPP30 | 9606.ENSP00000389182 | UniProt Keywords | KW-0025 | Alternative splicing |
| RPP30 | 9606.ENSP00000389182 | UniProt Keywords | KW-0378 | Hydrolase |
| RPP30 | 9606.ENSP00000389182 | UniProt Keywords | KW-0539 | Nucleus |
| RPP30 | 9606.ENSP00000389182 | UniProt Keywords | KW-0597 | Phosphoprotein |
| RPP30 | 9606.ENSP00000389182 | UniProt Keywords | KW-0694 | RNA-binding |
| RPP30 | 9606.ENSP00000389182 | UniProt Keywords | KW-0698 | rRNA processing |
| RPP30 | 9606.ENSP00000389182 | UniProt Keywords | KW-0819 | tRNA processing |
| RPP30 | 9606.ENSP00000389182 | Pfam | PF01876 | RNase P subunit p30 |
| RPP30 | 9606.ENSP00000389182 | InterPro | IPR002738 | RNase P subunit p30 |
| RPP30 | 9606.ENSP00000389182 | InterPro | IPR016195 | Polymerase/histidinol phosphatase-like |
| SLC4A10 | 9606.ENSP00000393066 | GO Process | GO:0003008 | System process |
| SLC4A10 | 9606.ENSP00000393066 | GO Process | GO:0006810 | Transport |
| SLC4A10 | 9606.ENSP00000393066 | GO Process | GO:0006811 | Ion transport |
| SLC4A10 | 9606.ENSP00000393066 | GO Process | GO:0006812 | Cation transport |
| SLC4A10 | 9606.ENSP00000393066 | GO Process | GO:0006814 | Sodium ion transport |
| SLC4A10 | 9606.ENSP00000393066 | GO Process | GO:0006820 | Anion transport |
| SLC4A10 | 9606.ENSP00000393066 | GO Process | GO:0006821 | Chloride transport |
| SLC4A10 | 9606.ENSP00000393066 | GO Process | GO:0006873 | Cellular ion homeostasis |
| SLC4A10 | 9606.ENSP00000393066 | GO Process | GO:0006885 | Regulation of pH |
| SLC4A10 | 9606.ENSP00000393066 | GO Process | GO:0007275 | Multicellular organism development |
| SLC4A10 | 9606.ENSP00000393066 | GO Process | GO:0007399 | Nervous system development |
| SLC4A10 | 9606.ENSP00000393066 | GO Process | GO:0007417 | Central nervous system development |

| | | | | |
|---|---|---|---|---|
| SLC4A10 | 9606.ENSP00000393066 | GO Process | GO:0007420 | Brain development |
| SLC4A10 | 9606.ENSP00000393066 | GO Process | GO:0007600 | Sensory perception |
| SLC4A10 | 9606.ENSP00000393066 | GO Process | GO:0007601 | Visual perception |
| SLC4A10 | 9606.ENSP00000393066 | GO Process | GO:0007610 | Behavior |
| SLC4A10 | 9606.ENSP00000393066 | GO Process | GO:0007626 | Locomotory behavior |
| SLC4A10 | 9606.ENSP00000393066 | GO Process | GO:0009314 | Response to radiation |
| SLC4A10 | 9606.ENSP00000393066 | GO Process | GO:0009416 | Response to light stimulus |
| SLC4A10 | 9606.ENSP00000393066 | GO Process | GO:0009628 | Response to abiotic stimulus |
| SLC4A10 | 9606.ENSP00000393066 | GO Process | GO:0009653 | Anatomical structure morphogenesis |
| SLC4A10 | 9606.ENSP00000393066 | GO Process | GO:0009791 | Post-embryonic development |
| SLC4A10 | 9606.ENSP00000393066 | GO Process | GO:0009887 | Animal organ morphogenesis |
| SLC4A10 | 9606.ENSP00000393066 | GO Process | GO:0009987 | Cellular process |
| SLC4A10 | 9606.ENSP00000393066 | GO Process | GO:0010646 | Regulation of cell communication |
| SLC4A10 | 9606.ENSP00000393066 | GO Process | GO:0015698 | Inorganic anion transport |
| SLC4A10 | 9606.ENSP00000393066 | GO Process | GO:0015701 | Bicarbonate transport |
| SLC4A10 | 9606.ENSP00000393066 | GO Process | GO:0015711 | Organic anion transport |
| SLC4A10 | 9606.ENSP00000393066 | GO Process | GO:0019725 | Cellular homeostasis |
| SLC4A10 | 9606.ENSP00000393066 | GO Process | GO:0021859 | Pyramidal neuron differentiation |
| SLC4A10 | 9606.ENSP00000393066 | GO Process | GO:0021860 | Pyramidal neuron development |
| SLC4A10 | 9606.ENSP00000393066 | GO Process | GO:0021872 | Forebrain generation of neurons |
| SLC4A10 | 9606.ENSP00000393066 | GO Process | GO:0021879 | Forebrain neuron differentiation |
| SLC4A10 | 9606.ENSP00000393066 | GO Process | GO:0021884 | Forebrain neuron development |
| SLC4A10 | 9606.ENSP00000393066 | GO Process | GO:0021953 | Central nervous system neuron differentiation |
| SLC4A10 | 9606.ENSP00000393066 | GO Process | GO:0021954 | Central nervous system neuron development |
| SLC4A10 | 9606.ENSP00000393066 | GO Process | GO:0022008 | Neurogenesis |
| SLC4A10 | 9606.ENSP00000393066 | GO Process | GO:0023051 | Regulation of signaling |
| SLC4A10 | 9606.ENSP00000393066 | GO Process | GO:0030001 | Metal ion transport |
| SLC4A10 | 9606.ENSP00000393066 | GO Process | GO:0030003 | Cellular cation homeostasis |
| SLC4A10 | 9606.ENSP00000393066 | GO Process | GO:0030004 | Cellular monovalent inorganic cation homeostasis |
| SLC4A10 | 9606.ENSP00000393066 | GO Process | GO:0030154 | Cell differentiation |
| SLC4A10 | 9606.ENSP00000393066 | GO Process | GO:0030182 | Neuron differentiation |
| SLC4A10 | 9606.ENSP00000393066 | GO Process | GO:0030641 | Regulation of cellular pH |
| SLC4A10 | 9606.ENSP00000393066 | GO Process | GO:0030900 | Forebrain development |
| SLC4A10 | 9606.ENSP00000393066 | GO Process | GO:0032501 | Multicellular organismal process |
| SLC4A10 | 9606.ENSP00000393066 | GO Process | GO:0032502 | Developmental process |

| | | | | |
|---|---|---|---|---|
| SLC4A10 | 9606.ENSP00000393066 | GO Process | GO:0034220 | Ion transmembrane transport |
| SLC4A10 | 9606.ENSP00000393066 | GO Process | GO:0035264 | Multicellular organism growth |
| SLC4A10 | 9606.ENSP00000393066 | GO Process | GO:0035640 | Exploration behavior |
| SLC4A10 | 9606.ENSP00000393066 | GO Process | GO:0035641 | Locomotory exploration behavior |
| SLC4A10 | 9606.ENSP00000393066 | GO Process | GO:0035725 | Sodium ion transmembrane transport |
| SLC4A10 | 9606.ENSP00000393066 | GO Process | GO:0040007 | Growth |
| SLC4A10 | 9606.ENSP00000393066 | GO Process | GO:0042592 | Homeostatic process |
| SLC4A10 | 9606.ENSP00000393066 | GO Process | GO:0048167 | Regulation of synaptic plasticity |
| SLC4A10 | 9606.ENSP00000393066 | GO Process | GO:0048168 | Regulation of neuronal synaptic plasticity |
| SLC4A10 | 9606.ENSP00000393066 | GO Process | GO:0048172 | Regulation of short-term neuronal synaptic plasticity |
| SLC4A10 | 9606.ENSP00000393066 | GO Process | GO:0048468 | Cell development |
| SLC4A10 | 9606.ENSP00000393066 | GO Process | GO:0048513 | Animal organ development |
| SLC4A10 | 9606.ENSP00000393066 | GO Process | GO:0048589 | Developmental growth |
| SLC4A10 | 9606.ENSP00000393066 | GO Process | GO:0048666 | Neuron development |
| SLC4A10 | 9606.ENSP00000393066 | GO Process | GO:0048699 | Generation of neurons |
| SLC4A10 | 9606.ENSP00000393066 | GO Process | GO:0048731 | System development |
| SLC4A10 | 9606.ENSP00000393066 | GO Process | GO:0048854 | Brain morphogenesis |
| SLC4A10 | 9606.ENSP00000393066 | GO Process | GO:0048856 | Anatomical structure development |
| SLC4A10 | 9606.ENSP00000393066 | GO Process | GO:0048869 | Cellular developmental process |
| SLC4A10 | 9606.ENSP00000393066 | GO Process | GO:0048878 | Chemical homeostasis |
| SLC4A10 | 9606.ENSP00000393066 | GO Process | GO:0050789 | Regulation of biological process |
| SLC4A10 | 9606.ENSP00000393066 | GO Process | GO:0050794 | Regulation of cellular process |
| SLC4A10 | 9606.ENSP00000393066 | GO Process | GO:0050801 | Ion homeostasis |
| SLC4A10 | 9606.ENSP00000393066 | GO Process | GO:0050804 | Modulation of chemical synaptic transmission |
| SLC4A10 | 9606.ENSP00000393066 | GO Process | GO:0050877 | Nervous system process |
| SLC4A10 | 9606.ENSP00000393066 | GO Process | GO:0050896 | Response to stimulus |
| SLC4A10 | 9606.ENSP00000393066 | GO Process | GO:0050953 | Sensory perception of light stimulus |
| SLC4A10 | 9606.ENSP00000393066 | GO Process | GO:0051179 | Localization |
| SLC4A10 | 9606.ENSP00000393066 | GO Process | GO:0051234 | Establishment of localization |
| SLC4A10 | 9606.ENSP00000393066 | GO Process | GO:0051453 | Regulation of intracellular pH |
| SLC4A10 | 9606.ENSP00000393066 | GO Process | GO:0055067 | Monovalent inorganic cation homeostasis |
| SLC4A10 | 9606.ENSP00000393066 | GO Process | GO:0055080 | Cation homeostasis |
| SLC4A10 | 9606.ENSP00000393066 | GO Process | GO:0055082 | Cellular chemical homeostasis |
| SLC4A10 | 9606.ENSP00000393066 | GO Process | GO:0055085 | Transmembrane transport |
| SLC4A10 | 9606.ENSP00000393066 | GO Process | GO:0060322 | Head development |

| | | | | |
|---|---|---|---|---|
| SLC4A10 | 9606.ENSP00000393066 | GO Process | GO:0065007 | Biological regulation |
| SLC4A10 | 9606.ENSP00000393066 | GO Process | GO:0065008 | Regulation of biological quality |
| SLC4A10 | 9606.ENSP00000393066 | GO Process | GO:0071702 | Organic substance transport |
| SLC4A10 | 9606.ENSP00000393066 | GO Process | GO:0098655 | Cation transmembrane transport |
| SLC4A10 | 9606.ENSP00000393066 | GO Process | GO:0098656 | Anion transmembrane transport |
| SLC4A10 | 9606.ENSP00000393066 | GO Process | GO:0098660 | Inorganic ion transmembrane transport |
| SLC4A10 | 9606.ENSP00000393066 | GO Process | GO:0098662 | Inorganic cation transmembrane transport |
| SLC4A10 | 9606.ENSP00000393066 | GO Process | GO:0098771 | Inorganic ion homeostasis |
| SLC4A10 | 9606.ENSP00000393066 | GO Process | GO:0099177 | Regulation of trans-synaptic signaling |
| SLC4A10 | 9606.ENSP00000393066 | GO Process | GO:1902600 | Proton transmembrane transport |
| SLC4A10 | 9606.ENSP00000393066 | GO Function | GO:0005215 | Transporter activity |
| SLC4A10 | 9606.ENSP00000393066 | GO Function | GO:0005452 | Inorganic anion exchanger activity |
| SLC4A10 | 9606.ENSP00000393066 | GO Function | GO:0008324 | Cation transmembrane transporter activity |
| SLC4A10 | 9606.ENSP00000393066 | GO Function | GO:0008509 | Anion transmembrane transporter activity |
| SLC4A10 | 9606.ENSP00000393066 | GO Function | GO:0008510 | Sodium:bicarbonate symporter activity |
| SLC4A10 | 9606.ENSP00000393066 | GO Function | GO:0008514 | Organic anion transmembrane transporter activity |
| SLC4A10 | 9606.ENSP00000393066 | GO Function | GO:0015075 | Ion transmembrane transporter activity |
| SLC4A10 | 9606.ENSP00000393066 | GO Function | GO:0015081 | Sodium ion transmembrane transporter activity |
| SLC4A10 | 9606.ENSP00000393066 | GO Function | GO:0015103 | Inorganic anion transmembrane transporter activity |
| SLC4A10 | 9606.ENSP00000393066 | GO Function | GO:0015106 | Bicarbonate transmembrane transporter activity |
| SLC4A10 | 9606.ENSP00000393066 | GO Function | GO:0015291 | Secondary active transmembrane transporter activity |
| SLC4A10 | 9606.ENSP00000393066 | GO Function | GO:0015293 | Symporter activity |
| SLC4A10 | 9606.ENSP00000393066 | GO Function | GO:0015294 | Solute:cation symporter activity |
| SLC4A10 | 9606.ENSP00000393066 | GO Function | GO:0015297 | Antiporter activity |
| SLC4A10 | 9606.ENSP00000393066 | GO Function | GO:0015301 | Anion:anion antiporter activity |
| SLC4A10 | 9606.ENSP00000393066 | GO Function | GO:0015318 | Inorganic molecular entity transmembrane transporter activity |
| SLC4A10 | 9606.ENSP00000393066 | GO Function | GO:0015370 | Solute:sodium symporter activity |
| SLC4A10 | 9606.ENSP00000393066 | GO Function | GO:0022804 | Active transmembrane transporter activity |
| SLC4A10 | 9606.ENSP00000393066 | GO Function | GO:0022853 | Active ion transmembrane transporter activity |
| SLC4A10 | 9606.ENSP00000393066 | GO Function | GO:0022857 | Transmembrane transporter activity |
| SLC4A10 | 9606.ENSP00000393066 | GO Function | GO:0022890 | Inorganic cation transmembrane transporter activity |
| SLC4A10 | 9606.ENSP00000393066 | GO Function | GO:0046873 | Metal ion transmembrane transporter activity |
| SLC4A10 | 9606.ENSP00000393066 | GO Function | GO:0140323 | Solute:anion antiporter activity |
| SLC4A10 | 9606.ENSP00000393066 | GO Function | GO:0140410 | Solute:bicarbonate symporter activity |
| SLC4A10 | 9606.ENSP00000393066 | GO Component | GO:0005886 | Plasma membrane |

| | | | | |
|---|---|---|---|---|
| SLC4A10 | 9606.ENSP00000393066 | GO Component | GO:0009925 | Basal plasma membrane |
| SLC4A10 | 9606.ENSP00000393066 | GO Component | GO:0016020 | Membrane |
| SLC4A10 | 9606.ENSP00000393066 | GO Component | GO:0016021 | Integral component of membrane |
| SLC4A10 | 9606.ENSP00000393066 | GO Component | GO:0016323 | Basolateral plasma membrane |
| SLC4A10 | 9606.ENSP00000393066 | GO Component | GO:0016324 | Apical plasma membrane |
| SLC4A10 | 9606.ENSP00000393066 | GO Component | GO:0030054 | Cell junction |
| SLC4A10 | 9606.ENSP00000393066 | GO Component | GO:0030424 | Axon |
| SLC4A10 | 9606.ENSP00000393066 | GO Component | GO:0030425 | Dendrite |
| SLC4A10 | 9606.ENSP00000393066 | GO Component | GO:0031224 | Intrinsic component of membrane |
| SLC4A10 | 9606.ENSP00000393066 | GO Component | GO:0036477 | Somatodendritic compartment |
| SLC4A10 | 9606.ENSP00000393066 | GO Component | GO:0042995 | Cell projection |
| SLC4A10 | 9606.ENSP00000393066 | GO Component | GO:0043005 | Neuron projection |
| SLC4A10 | 9606.ENSP00000393066 | GO Component | GO:0043025 | Neuronal cell body |
| SLC4A10 | 9606.ENSP00000393066 | GO Component | GO:0043204 | Perikaryon |
| SLC4A10 | 9606.ENSP00000393066 | GO Component | GO:0043679 | Axon terminus |
| SLC4A10 | 9606.ENSP00000393066 | GO Component | GO:0044297 | Cell body |
| SLC4A10 | 9606.ENSP00000393066 | GO Component | GO:0044306 | Neuron projection terminus |
| SLC4A10 | 9606.ENSP00000393066 | GO Component | GO:0045177 | Apical part of cell |
| SLC4A10 | 9606.ENSP00000393066 | GO Component | GO:0045178 | Basal part of cell |
| SLC4A10 | 9606.ENSP00000393066 | GO Component | GO:0045202 | Synapse |
| SLC4A10 | 9606.ENSP00000393066 | GO Component | GO:0070161 | Anchoring junction |
| SLC4A10 | 9606.ENSP00000393066 | GO Component | GO:0071944 | Cell periphery |
| SLC4A10 | 9606.ENSP00000393066 | GO Component | GO:0097440 | Apical dendrite |
| SLC4A10 | 9606.ENSP00000393066 | GO Component | GO:0097441 | Basal dendrite |
| SLC4A10 | 9606.ENSP00000393066 | GO Component | GO:0097442 | CA3 pyramidal cell dendrite |
| SLC4A10 | 9606.ENSP00000393066 | GO Component | GO:0097447 | Dendritic tree |
| SLC4A10 | 9606.ENSP00000393066 | GO Component | GO:0098590 | Plasma membrane region |
| SLC4A10 | 9606.ENSP00000393066 | GO Component | GO:0098793 | Presynapse |
| SLC4A10 | 9606.ENSP00000393066 | GO Component | GO:0098794 | Postsynapse |
| SLC4A10 | 9606.ENSP00000393066 | GO Component | GO:0110165 | Cellular anatomical entity |
| SLC4A10 | 9606.ENSP00000393066 | GO Component | GO:0120025 | Plasma membrane bounded cell projection |
| SLC4A10 | 9606.ENSP00000393066 | GO Component | GO:0150034 | Distal axon |
| SLC4A10 | 9606.ENSP00000393066 | STRING clusters | CL:12531 | Mixed, incl. Transport of inorganic cations/anions and amino acids/oligopeptides, and Carbonate dehydratase activity |

| | | | | |
|---|---|---|---|---|
| SLC4A10 | 9606.ENSP00000393066 | STRING clusters | CL:12532 | Mixed, incl. Transport of inorganic cations/anions and amino acids/oligopeptides, and Renal tubular transport disease |
| SLC4A10 | 9606.ENSP00000393066 | STRING clusters | CL:12533 | Mixed, incl. Antiport, and Chloride ion homeostasis |
| SLC4A10 | 9606.ENSP00000393066 | STRING clusters | CL:12534 | Mixed, incl. Antiport, and Chloride ion homeostasis |
| SLC4A10 | 9606.ENSP00000393066 | STRING clusters | CL:12605 | Chloride ion homeostasis, and Bartter syndrome |
| SLC4A10 | 9606.ENSP00000393066 | Reactome | HSA-382551 | Transport of small molecules |
| SLC4A10 | 9606.ENSP00000393066 | Reactome | HSA-425381 | Bicarbonate transporters |
| SLC4A10 | 9606.ENSP00000393066 | Reactome | HSA-425393 | Transport of inorganic cations/anions and amino acids/oligopeptides |
| SLC4A10 | 9606.ENSP00000393066 | Reactome | HSA-425407 | SLC-mediated transmembrane transport |
| SLC4A10 | 9606.ENSP00000393066 | DISEASES | DOID:12382 | Complex partial epilepsy |
| SLC4A10 | 9606.ENSP00000393066 | DISEASES | DOID:1826 | Epilepsy |
| SLC4A10 | 9606.ENSP00000393066 | DISEASES | DOID:2234 | Focal epilepsy |
| SLC4A10 | 9606.ENSP00000393066 | DISEASES | DOID:331 | Central nervous system disease |
| SLC4A10 | 9606.ENSP00000393066 | DISEASES | DOID:4 | Disease |
| SLC4A10 | 9606.ENSP00000393066 | DISEASES | DOID:7 | Disease of anatomical entity |
| SLC4A10 | 9606.ENSP00000393066 | DISEASES | DOID:863 | Nervous system disease |
| SLC4A10 | 9606.ENSP00000393066 | DISEASES | DOID:936 | Brain disease |
| SLC4A10 | 9606.ENSP00000393066 | TISSUES | BTO:0000000 | Tissues, cell types and enzyme sources |
| SLC4A10 | 9606.ENSP00000393066 | TISSUES | BTO:0000042 | Animal |
| SLC4A10 | 9606.ENSP00000393066 | TISSUES | BTO:0000142 | Brain |
| SLC4A10 | 9606.ENSP00000393066 | TISSUES | BTO:0000146 | Brain stem |
| SLC4A10 | 9606.ENSP00000393066 | TISSUES | BTO:0000227 | Central nervous system |
| SLC4A10 | 9606.ENSP00000393066 | TISSUES | BTO:0000231 | Cerebral hemisphere |
| SLC4A10 | 9606.ENSP00000393066 | TISSUES | BTO:0000232 | Cerebellum |
| SLC4A10 | 9606.ENSP00000393066 | TISSUES | BTO:0000233 | Cerebral cortex |
| SLC4A10 | 9606.ENSP00000393066 | TISSUES | BTO:0000239 | Telencephalon |
| SLC4A10 | 9606.ENSP00000393066 | TISSUES | BTO:0000282 | Head |
| SLC4A10 | 9606.ENSP00000393066 | TISSUES | BTO:0000445 | Cerebral lobe |
| SLC4A10 | 9606.ENSP00000393066 | TISSUES | BTO:0000478 | Forebrain |
| SLC4A10 | 9606.ENSP00000393066 | TISSUES | BTO:0000672 | Hindbrain |
| SLC4A10 | 9606.ENSP00000393066 | TISSUES | BTO:0000673 | Metencephalon |
| SLC4A10 | 9606.ENSP00000393066 | TISSUES | BTO:0000928 | Limbic system |
| SLC4A10 | 9606.ENSP00000393066 | TISSUES | BTO:0001355 | Temporal lobe |
| SLC4A10 | 9606.ENSP00000393066 | TISSUES | BTO:0001484 | Nervous system |
| SLC4A10 | 9606.ENSP00000393066 | TISSUES | BTO:0001489 | Whole body |

| | | | | |
|---|---|---|---|---|
| SLC4A10 | 9606.ENSP00000393066 | COMPARTMENTS | GOCC:0005886 | Plasma membrane |
| SLC4A10 | 9606.ENSP00000393066 | COMPARTMENTS | GOCC:0009925 | Basal plasma membrane |
| SLC4A10 | 9606.ENSP00000393066 | COMPARTMENTS | GOCC:0016020 | Membrane |
| SLC4A10 | 9606.ENSP00000393066 | COMPARTMENTS | GOCC:0016021 | Integral component of membrane |
| SLC4A10 | 9606.ENSP00000393066 | COMPARTMENTS | GOCC:0016323 | Basolateral plasma membrane |
| SLC4A10 | 9606.ENSP00000393066 | COMPARTMENTS | GOCC:0016324 | Apical plasma membrane |
| SLC4A10 | 9606.ENSP00000393066 | COMPARTMENTS | GOCC:0030054 | Cell junction |
| SLC4A10 | 9606.ENSP00000393066 | COMPARTMENTS | GOCC:0030424 | Axon |
| SLC4A10 | 9606.ENSP00000393066 | COMPARTMENTS | GOCC:0030425 | Dendrite |
| SLC4A10 | 9606.ENSP00000393066 | COMPARTMENTS | GOCC:0031224 | Intrinsic component of membrane |
| SLC4A10 | 9606.ENSP00000393066 | COMPARTMENTS | GOCC:0033774 | Basal labyrinth |
| SLC4A10 | 9606.ENSP00000393066 | COMPARTMENTS | GOCC:0036477 | Somatodendritic compartment |
| SLC4A10 | 9606.ENSP00000393066 | COMPARTMENTS | GOCC:0042995 | Cell projection |
| SLC4A10 | 9606.ENSP00000393066 | COMPARTMENTS | GOCC:0043005 | Neuron projection |
| SLC4A10 | 9606.ENSP00000393066 | COMPARTMENTS | GOCC:0043025 | Neuronal cell body |
| SLC4A10 | 9606.ENSP00000393066 | COMPARTMENTS | GOCC:0043679 | Axon terminus |
| SLC4A10 | 9606.ENSP00000393066 | COMPARTMENTS | GOCC:0044297 | Cell body |
| SLC4A10 | 9606.ENSP00000393066 | COMPARTMENTS | GOCC:0044306 | Neuron projection terminus |
| SLC4A10 | 9606.ENSP00000393066 | COMPARTMENTS | GOCC:0045177 | Apical part of cell |
| SLC4A10 | 9606.ENSP00000393066 | COMPARTMENTS | GOCC:0045178 | Basal part of cell |
| SLC4A10 | 9606.ENSP00000393066 | COMPARTMENTS | GOCC:0045202 | Synapse |
| SLC4A10 | 9606.ENSP00000393066 | COMPARTMENTS | GOCC:0071944 | Cell periphery |
| SLC4A10 | 9606.ENSP00000393066 | COMPARTMENTS | GOCC:0097447 | Dendritic tree |
| SLC4A10 | 9606.ENSP00000393066 | COMPARTMENTS | GOCC:0098590 | Plasma membrane region |
| SLC4A10 | 9606.ENSP00000393066 | COMPARTMENTS | GOCC:0098793 | Presynapse |
| SLC4A10 | 9606.ENSP00000393066 | COMPARTMENTS | GOCC:0110165 | Cellular anatomical entity |
| SLC4A10 | 9606.ENSP00000393066 | COMPARTMENTS | GOCC:0120025 | Plasma membrane bounded cell projection |
| SLC4A10 | 9606.ENSP00000393066 | COMPARTMENTS | GOCC:0150034 | Distal axon |
| SLC4A10 | 9606.ENSP00000393066 | Monarch | EFO:0000719 | Temporal measurement |
| SLC4A10 | 9606.ENSP00000393066 | Monarch | EFO:0000727 | Treatment |
| SLC4A10 | 9606.ENSP00000393066 | Monarch | EFO:0003925 | Cognition |
| SLC4A10 | 9606.ENSP00000393066 | Monarch | EFO:0004302 | Anthropometric measurement |
| SLC4A10 | 9606.ENSP00000393066 | Monarch | EFO:0004318 | Smoking behavior |
| SLC4A10 | 9606.ENSP00000393066 | Monarch | EFO:0004323 | Mental process |
| SLC4A10 | 9606.ENSP00000393066 | Monarch | EFO:0004324 | Body weights and measures |

| | | | | |
|---|---|---|---|---|
| SLC4A10 | 9606.ENSP00000393066 | Monarch | EFO:0004337 | Intelligence |
| SLC4A10 | 9606.ENSP00000393066 | Monarch | EFO:0004339 | Body height |
| SLC4A10 | 9606.ENSP00000393066 | Monarch | EFO:0004340 | Body mass index |
| SLC4A10 | 9606.ENSP00000393066 | Monarch | EFO:0004346 | Neuroimaging measurement |
| SLC4A10 | 9606.ENSP00000393066 | Monarch | EFO:0004464 | Brain measurement |
| SLC4A10 | 9606.ENSP00000393066 | Monarch | EFO:0004542 | Planned process |
| SLC4A10 | 9606.ENSP00000393066 | Monarch | EFO:0004784 | Self reported educational attainment |
| SLC4A10 | 9606.ENSP00000393066 | Monarch | EFO:0004840 | Cortical thickness |
| SLC4A10 | 9606.ENSP00000393066 | Monarch | EFO:0004870 | Sleep measurement |
| SLC4A10 | 9606.ENSP00000393066 | Monarch | EFO:0004875 | Mathematical ability |
| SLC4A10 | 9606.ENSP00000393066 | Monarch | EFO:0004918 | Age at diagnosis |
| SLC4A10 | 9606.ENSP00000393066 | Monarch | EFO:0004949 | Clinical temporal measurement |
| SLC4A10 | 9606.ENSP00000393066 | Monarch | EFO:0005670 | Smoking initiation |
| SLC4A10 | 9606.ENSP00000393066 | Monarch | EFO:0005671 | Smoking behaviour measurement |
| SLC4A10 | 9606.ENSP00000393066 | Monarch | EFO:0006527 | Smoking status measurement |
| SLC4A10 | 9606.ENSP00000393066 | Monarch | EFO:0006848 | Mental or behavioural disorder biomarker |
| SLC4A10 | 9606.ENSP00000393066 | Monarch | EFO:0006918 | Female fertility |
| SLC4A10 | 9606.ENSP00000393066 | Monarch | EFO:0006923 | Fertility measurement |
| SLC4A10 | 9606.ENSP00000393066 | Monarch | EFO:0007660 | Neuroticism measurement |
| SLC4A10 | 9606.ENSP00000393066 | Monarch | EFO:0007803 | Emotional symptom measurement |
| SLC4A10 | 9606.ENSP00000393066 | Monarch | EFO:0007820 | Cognitive behavioural therapy |
| SLC4A10 | 9606.ENSP00000393066 | Monarch | EFO:0007828 | Daytime rest measurement |
| SLC4A10 | 9606.ENSP00000393066 | Monarch | EFO:0007878 | Alcohol consumption measurement |
| SLC4A10 | 9606.ENSP00000393066 | Monarch | EFO:0007911 | Personality trait measurement |
| SLC4A10 | 9606.ENSP00000393066 | Monarch | EFO:0007967 | Blood osmolality measurement |
| SLC4A10 | 9606.ENSP00000393066 | Monarch | EFO:0008354 | Cognitive function measurement |
| SLC4A10 | 9606.ENSP00000393066 | Monarch | EFO:0008394 | Verbal-numerical reasoning measurement |
| SLC4A10 | 9606.ENSP00000393066 | Monarch | EFO:0009282 | Sodium measurement |
| SLC4A10 | 9606.ENSP00000393066 | Monarch | EFO:0009592 | Social interaction measurement |
| SLC4A10 | 9606.ENSP00000393066 | Monarch | EFO:0010724 | Lifestyle measurement |
| SLC4A10 | 9606.ENSP00000393066 | Monarch | EFO:0010736 | Cortical surface area measurement |
| SLC4A10 | 9606.ENSP00000393066 | Monarch | EFO:0011015 | Educational attainment |
| SLC4A10 | 9606.ENSP00000393066 | UniProt Keywords | KW-0025 | Alternative splicing |
| SLC4A10 | 9606.ENSP00000393066 | UniProt Keywords | KW-0050 | Antiport |
| SLC4A10 | 9606.ENSP00000393066 | UniProt Keywords | KW-0325 | Glycoprotein |

| | | | | |
|---|---|---|---|---|
| SLC4A10 | 9606.ENSP00000393066 | UniProt Keywords | KW-0406 | Ion transport |
| SLC4A10 | 9606.ENSP00000393066 | UniProt Keywords | KW-0472 | Membrane |
| SLC4A10 | 9606.ENSP00000393066 | UniProt Keywords | KW-0597 | Phosphoprotein |
| SLC4A10 | 9606.ENSP00000393066 | UniProt Keywords | KW-0739 | Sodium transport |
| SLC4A10 | 9606.ENSP00000393066 | UniProt Keywords | KW-0769 | Symport |
| SLC4A10 | 9606.ENSP00000393066 | UniProt Keywords | KW-0770 | Synapse |
| SLC4A10 | 9606.ENSP00000393066 | UniProt Keywords | KW-0812 | Transmembrane |
| SLC4A10 | 9606.ENSP00000393066 | UniProt Keywords | KW-0813 | Transport |
| SLC4A10 | 9606.ENSP00000393066 | UniProt Keywords | KW-0915 | Sodium |
| SLC4A10 | 9606.ENSP00000393066 | UniProt Keywords | KW-0965 | Cell junction |
| SLC4A10 | 9606.ENSP00000393066 | UniProt Keywords | KW-0966 | Cell projection |
| SLC4A10 | 9606.ENSP00000393066 | UniProt Keywords | KW-1003 | Cell membrane |
| SLC4A10 | 9606.ENSP00000393066 | UniProt Keywords | KW-1133 | Transmembrane helix |
| SLC4A10 | 9606.ENSP00000393066 | Pfam | PF00955 | HCO3- transporter family |
| SLC4A10 | 9606.ENSP00000393066 | Pfam | PF07565 | Band 3 cytoplasmic domain |
| SLC4A10 | 9606.ENSP00000393066 | InterPro | IPR003020 | Bicarbonate transporter, eukaryotic |
| SLC4A10 | 9606.ENSP00000393066 | InterPro | IPR003024 | Sodium bicarbonate cotransporter |
| SLC4A10 | 9606.ENSP00000393066 | InterPro | IPR011531 | Bicarbonate transporter-like, transmembrane domain |
| SLC4A10 | 9606.ENSP00000393066 | InterPro | IPR013769 | Band 3 cytoplasmic domain |
| SLC4A10 | 9606.ENSP00000393066 | InterPro | IPR016152 | Phosphotransferase/anion transporter |
| SMAP1 | 9606.ENSP00000359484 | GO Process | GO:0002682 | Regulation of immune system process |
| SMAP1 | 9606.ENSP00000359484 | GO Process | GO:0030100 | Regulation of endocytosis |
| SMAP1 | 9606.ENSP00000359484 | GO Process | GO:0032879 | Regulation of localization |
| SMAP1 | 9606.ENSP00000359484 | GO Process | GO:0045595 | Regulation of cell differentiation |
| SMAP1 | 9606.ENSP00000359484 | GO Process | GO:0045597 | Positive regulation of cell differentiation |
| SMAP1 | 9606.ENSP00000359484 | GO Process | GO:0045637 | Regulation of myeloid cell differentiation |
| SMAP1 | 9606.ENSP00000359484 | GO Process | GO:0045639 | Positive regulation of myeloid cell differentiation |
| SMAP1 | 9606.ENSP00000359484 | GO Process | GO:0045646 | Regulation of erythrocyte differentiation |
| SMAP1 | 9606.ENSP00000359484 | GO Process | GO:0045648 | Positive regulation of erythrocyte differentiation |
| SMAP1 | 9606.ENSP00000359484 | GO Process | GO:0048259 | Regulation of receptor-mediated endocytosis |
| SMAP1 | 9606.ENSP00000359484 | GO Process | GO:0048518 | Positive regulation of biological process |
| SMAP1 | 9606.ENSP00000359484 | GO Process | GO:0048522 | Positive regulation of cellular process |
| SMAP1 | 9606.ENSP00000359484 | GO Process | GO:0050789 | Regulation of biological process |
| SMAP1 | 9606.ENSP00000359484 | GO Process | GO:0050790 | Regulation of catalytic activity |
| SMAP1 | 9606.ENSP00000359484 | GO Process | GO:0050793 | Regulation of developmental process |

| | | | | |
|---|---|---|---|---|
| SMAP1 | 9606.ENSP00000359484 | GO Process | GO:0050794 | Regulation of cellular process |
| SMAP1 | 9606.ENSP00000359484 | GO Process | GO:0051049 | Regulation of transport |
| SMAP1 | 9606.ENSP00000359484 | GO Process | GO:0051094 | Positive regulation of developmental process |
| SMAP1 | 9606.ENSP00000359484 | GO Process | GO:0051128 | Regulation of cellular component organization |
| SMAP1 | 9606.ENSP00000359484 | GO Process | GO:0051239 | Regulation of multicellular organismal process |
| SMAP1 | 9606.ENSP00000359484 | GO Process | GO:0060627 | Regulation of vesicle-mediated transport |
| SMAP1 | 9606.ENSP00000359484 | GO Process | GO:0065007 | Biological regulation |
| SMAP1 | 9606.ENSP00000359484 | GO Process | GO:0065009 | Regulation of molecular function |
| SMAP1 | 9606.ENSP00000359484 | GO Process | GO:1903706 | Regulation of hemopoiesis |
| SMAP1 | 9606.ENSP00000359484 | GO Process | GO:2000026 | Regulation of multicellular organismal development |
| SMAP1 | 9606.ENSP00000359484 | GO Process | GO:2000369 | Regulation of clathrin-dependent endocytosis |
| SMAP1 | 9606.ENSP00000359484 | GO Function | GO:0005096 | GTPase activator activity |
| SMAP1 | 9606.ENSP00000359484 | GO Function | GO:0005488 | Binding |
| SMAP1 | 9606.ENSP00000359484 | GO Function | GO:0005515 | Protein binding |
| SMAP1 | 9606.ENSP00000359484 | GO Function | GO:0008047 | Enzyme activator activity |
| SMAP1 | 9606.ENSP00000359484 | GO Function | GO:0030234 | Enzyme regulator activity |
| SMAP1 | 9606.ENSP00000359484 | GO Function | GO:0030276 | Clathrin binding |
| SMAP1 | 9606.ENSP00000359484 | GO Function | GO:0030695 | GTPase regulator activity |
| SMAP1 | 9606.ENSP00000359484 | GO Function | GO:0043167 | Ion binding |
| SMAP1 | 9606.ENSP00000359484 | GO Function | GO:0043169 | Cation binding |
| SMAP1 | 9606.ENSP00000359484 | GO Function | GO:0046872 | Metal ion binding |
| SMAP1 | 9606.ENSP00000359484 | GO Function | GO:0060589 | Nucleoside-triphosphatase regulator activity |
| SMAP1 | 9606.ENSP00000359484 | GO Function | GO:0098772 | Molecular function regulator activity |
| SMAP1 | 9606.ENSP00000359484 | GO Component | GO:0005622 | Intracellular anatomical structure |
| SMAP1 | 9606.ENSP00000359484 | GO Component | GO:0005737 | Cytoplasm |
| SMAP1 | 9606.ENSP00000359484 | GO Component | GO:0005886 | Plasma membrane |
| SMAP1 | 9606.ENSP00000359484 | GO Component | GO:0016020 | Membrane |
| SMAP1 | 9606.ENSP00000359484 | GO Component | GO:0071944 | Cell periphery |
| SMAP1 | 9606.ENSP00000359484 | GO Component | GO:0110165 | Cellular anatomical entity |
| SMAP1 | 9606.ENSP00000359484 | STRING clusters | CL:13766 | Mixed, incl. Membrane coat, and Sec7, C-terminal domain superfamily |
| SMAP1 | 9606.ENSP00000359484 | STRING clusters | CL:13767 | Clathrin coat, and Presynaptic endocytosis |
| SMAP1 | 9606.ENSP00000359484 | KEGG | hsa04144 | Endocytosis |
| SMAP1 | 9606.ENSP00000359484 | TISSUES | BTO:0000000 | Tissues, cell types and enzyme sources |
| SMAP1 | 9606.ENSP00000359484 | TISSUES | BTO:0000042 | Animal |
| SMAP1 | 9606.ENSP00000359484 | TISSUES | BTO:0000142 | Brain |

| | | | | |
|---|---|---|---|---|
| SMAP1 | 9606.ENSP00000359484 | TISSUES | BTO:0000227 | Central nervous system |
| SMAP1 | 9606.ENSP00000359484 | TISSUES | BTO:0000282 | Head |
| SMAP1 | 9606.ENSP00000359484 | TISSUES | BTO:0001484 | Nervous system |
| SMAP1 | 9606.ENSP00000359484 | TISSUES | BTO:0001489 | Whole body |
| SMAP1 | 9606.ENSP00000359484 | Monarch | EFO:0004747 | Protein measurement |
| SMAP1 | 9606.ENSP00000359484 | Monarch | EFO:0007937 | Blood protein measurement |
| SMAP1 | 9606.ENSP00000359484 | UniProt Keywords | KW-0025 | Alternative splicing |
| SMAP1 | 9606.ENSP00000359484 | UniProt Keywords | KW-0343 | GTPase activation |
| SMAP1 | 9606.ENSP00000359484 | UniProt Keywords | KW-0472 | Membrane |
| SMAP1 | 9606.ENSP00000359484 | UniProt Keywords | KW-0479 | Metal-binding |
| SMAP1 | 9606.ENSP00000359484 | UniProt Keywords | KW-0862 | Zinc |
| SMAP1 | 9606.ENSP00000359484 | UniProt Keywords | KW-0863 | Zinc-finger |
| SMAP1 | 9606.ENSP00000359484 | UniProt Keywords | KW-1003 | Cell membrane |
| SMAP1 | 9606.ENSP00000359484 | InterPro | IPR001164 | Arf GTPase activating protein |
| SMAP1 | 9606.ENSP00000359484 | InterPro | IPR037278 | ARFGAP/RecO-like zinc finger |
| SMAP1 | 9606.ENSP00000359484 | InterPro | IPR038508 | ArfGAP domain superfamily |
| SMAP1 | 9606.ENSP00000359484 | InterPro | IPR044732 | SMAP1-like, ArfGAP domain |
| SMAP1 | 9606.ENSP00000359484 | SMART | SM00105 | Putative GTP-ase activating proteins for the small GTPase, ARF |
| TFCP2L1 | 9606.ENSP00000263707 | GO Process | GO:0000122 | Negative regulation of transcription by RNA polymerase II |
| TFCP2L1 | 9606.ENSP00000263707 | GO Process | GO:0000902 | Cell morphogenesis |
| TFCP2L1 | 9606.ENSP00000263707 | GO Process | GO:0002064 | Epithelial cell development |
| TFCP2L1 | 9606.ENSP00000263707 | GO Process | GO:0002070 | Epithelial cell maturation |
| TFCP2L1 | 9606.ENSP00000263707 | GO Process | GO:0006355 | Regulation of transcription, DNA-templated |
| TFCP2L1 | 9606.ENSP00000263707 | GO Process | GO:0006357 | Regulation of transcription by RNA polymerase II |
| TFCP2L1 | 9606.ENSP00000263707 | GO Process | GO:0007028 | Cytoplasm organization |
| TFCP2L1 | 9606.ENSP00000263707 | GO Process | GO:0007275 | Multicellular organism development |
| TFCP2L1 | 9606.ENSP00000263707 | GO Process | GO:0007431 | Salivary gland development |
| TFCP2L1 | 9606.ENSP00000263707 | GO Process | GO:0008340 | Determination of adult lifespan |
| TFCP2L1 | 9606.ENSP00000263707 | GO Process | GO:0009653 | Anatomical structure morphogenesis |
| TFCP2L1 | 9606.ENSP00000263707 | GO Process | GO:0009888 | Tissue development |
| TFCP2L1 | 9606.ENSP00000263707 | GO Process | GO:0009889 | Regulation of biosynthetic process |
| TFCP2L1 | 9606.ENSP00000263707 | GO Process | GO:0009890 | Negative regulation of biosynthetic process |
| TFCP2L1 | 9606.ENSP00000263707 | GO Process | GO:0009891 | Positive regulation of biosynthetic process |
| TFCP2L1 | 9606.ENSP00000263707 | GO Process | GO:0009892 | Negative regulation of metabolic process |
| TFCP2L1 | 9606.ENSP00000263707 | GO Process | GO:0009893 | Positive regulation of metabolic process |

| | | | | |
|---|---|---|---|---|
| TFCP2L1 | 9606.ENSP00000263707 | GO Process | GO:0009987 | Cellular process |
| TFCP2L1 | 9606.ENSP00000263707 | GO Process | GO:0010468 | Regulation of gene expression |
| TFCP2L1 | 9606.ENSP00000263707 | GO Process | GO:0010556 | Regulation of macromolecule biosynthetic process |
| TFCP2L1 | 9606.ENSP00000263707 | GO Process | GO:0010557 | Positive regulation of macromolecule biosynthetic process |
| TFCP2L1 | 9606.ENSP00000263707 | GO Process | GO:0010558 | Negative regulation of macromolecule biosynthetic process |
| TFCP2L1 | 9606.ENSP00000263707 | GO Process | GO:0010604 | Positive regulation of macromolecule metabolic process |
| TFCP2L1 | 9606.ENSP00000263707 | GO Process | GO:0010605 | Negative regulation of macromolecule metabolic process |
| TFCP2L1 | 9606.ENSP00000263707 | GO Process | GO:0016043 | Cellular component organization |
| TFCP2L1 | 9606.ENSP00000263707 | GO Process | GO:0019219 | Regulation of nucleobase-containing compound metabolic process |
| TFCP2L1 | 9606.ENSP00000263707 | GO Process | GO:0019222 | Regulation of metabolic process |
| TFCP2L1 | 9606.ENSP00000263707 | GO Process | GO:0021700 | Developmental maturation |
| TFCP2L1 | 9606.ENSP00000263707 | GO Process | GO:0030154 | Cell differentiation |
| TFCP2L1 | 9606.ENSP00000263707 | GO Process | GO:0030855 | Epithelial cell differentiation |
| TFCP2L1 | 9606.ENSP00000263707 | GO Process | GO:0031323 | Regulation of cellular metabolic process |
| TFCP2L1 | 9606.ENSP00000263707 | GO Process | GO:0031324 | Negative regulation of cellular metabolic process |
| TFCP2L1 | 9606.ENSP00000263707 | GO Process | GO:0031325 | Positive regulation of cellular metabolic process |
| TFCP2L1 | 9606.ENSP00000263707 | GO Process | GO:0031326 | Regulation of cellular biosynthetic process |
| TFCP2L1 | 9606.ENSP00000263707 | GO Process | GO:0031327 | Negative regulation of cellular biosynthetic process |
| TFCP2L1 | 9606.ENSP00000263707 | GO Process | GO:0031328 | Positive regulation of cellular biosynthetic process |
| TFCP2L1 | 9606.ENSP00000263707 | GO Process | GO:0032501 | Multicellular organismal process |
| TFCP2L1 | 9606.ENSP00000263707 | GO Process | GO:0032502 | Developmental process |
| TFCP2L1 | 9606.ENSP00000263707 | GO Process | GO:0035272 | Exocrine system development |
| TFCP2L1 | 9606.ENSP00000263707 | GO Process | GO:0040008 | Regulation of growth |
| TFCP2L1 | 9606.ENSP00000263707 | GO Process | GO:0045892 | Negative regulation of transcription, DNA-templated |
| TFCP2L1 | 9606.ENSP00000263707 | GO Process | GO:0045893 | Positive regulation of transcription, DNA-templated |
| TFCP2L1 | 9606.ENSP00000263707 | GO Process | GO:0045927 | Positive regulation of growth |
| TFCP2L1 | 9606.ENSP00000263707 | GO Process | GO:0045934 | Negative regulation of nucleobase-containing compound metabolic process |
| TFCP2L1 | 9606.ENSP00000263707 | GO Process | GO:0045935 | Positive regulation of nucleobase-containing compound metabolic process |
| TFCP2L1 | 9606.ENSP00000263707 | GO Process | GO:0045944 | Positive regulation of transcription by RNA polymerase II |
| TFCP2L1 | 9606.ENSP00000263707 | GO Process | GO:0048468 | Cell development |
| TFCP2L1 | 9606.ENSP00000263707 | GO Process | GO:0048469 | Cell maturation |
| TFCP2L1 | 9606.ENSP00000263707 | GO Process | GO:0048513 | Animal organ development |
| TFCP2L1 | 9606.ENSP00000263707 | GO Process | GO:0048518 | Positive regulation of biological process |
| TFCP2L1 | 9606.ENSP00000263707 | GO Process | GO:0048519 | Negative regulation of biological process |

| | | | | |
|---|---|---|---|---|
| TFCP2L1 | 9606.ENSP00000263707 | GO Process | GO:0048522 | Positive regulation of cellular process |
| TFCP2L1 | 9606.ENSP00000263707 | GO Process | GO:0048523 | Negative regulation of cellular process |
| TFCP2L1 | 9606.ENSP00000263707 | GO Process | GO:0048731 | System development |
| TFCP2L1 | 9606.ENSP00000263707 | GO Process | GO:0048732 | Gland development |
| TFCP2L1 | 9606.ENSP00000263707 | GO Process | GO:0048856 | Anatomical structure development |
| TFCP2L1 | 9606.ENSP00000263707 | GO Process | GO:0048869 | Cellular developmental process |
| TFCP2L1 | 9606.ENSP00000263707 | GO Process | GO:0050789 | Regulation of biological process |
| TFCP2L1 | 9606.ENSP00000263707 | GO Process | GO:0050794 | Regulation of cellular process |
| TFCP2L1 | 9606.ENSP00000263707 | GO Process | GO:0051171 | Regulation of nitrogen compound metabolic process |
| TFCP2L1 | 9606.ENSP00000263707 | GO Process | GO:0051172 | Negative regulation of nitrogen compound metabolic process |
| TFCP2L1 | 9606.ENSP00000263707 | GO Process | GO:0051173 | Positive regulation of nitrogen compound metabolic process |
| TFCP2L1 | 9606.ENSP00000263707 | GO Process | GO:0051252 | Regulation of RNA metabolic process |
| TFCP2L1 | 9606.ENSP00000263707 | GO Process | GO:0051253 | Negative regulation of RNA metabolic process |
| TFCP2L1 | 9606.ENSP00000263707 | GO Process | GO:0051254 | Positive regulation of RNA metabolic process |
| TFCP2L1 | 9606.ENSP00000263707 | GO Process | GO:0060255 | Regulation of macromolecule metabolic process |
| TFCP2L1 | 9606.ENSP00000263707 | GO Process | GO:0060429 | Epithelium development |
| TFCP2L1 | 9606.ENSP00000263707 | GO Process | GO:0065007 | Biological regulation |
| TFCP2L1 | 9606.ENSP00000263707 | GO Process | GO:0071695 | Anatomical structure maturation |
| TFCP2L1 | 9606.ENSP00000263707 | GO Process | GO:0071840 | Cellular component organization or biogenesis |
| TFCP2L1 | 9606.ENSP00000263707 | GO Process | GO:0080090 | Regulation of primary metabolic process |
| TFCP2L1 | 9606.ENSP00000263707 | GO Process | GO:1902679 | Negative regulation of RNA biosynthetic process |
| TFCP2L1 | 9606.ENSP00000263707 | GO Process | GO:1902680 | Positive regulation of RNA biosynthetic process |
| TFCP2L1 | 9606.ENSP00000263707 | GO Process | GO:1903506 | Regulation of nucleic acid-templated transcription |
| TFCP2L1 | 9606.ENSP00000263707 | GO Process | GO:1903507 | Negative regulation of nucleic acid-templated transcription |
| TFCP2L1 | 9606.ENSP00000263707 | GO Process | GO:1903508 | Positive regulation of nucleic acid-templated transcription |
| TFCP2L1 | 9606.ENSP00000263707 | GO Process | GO:2001141 | Regulation of RNA biosynthetic process |
| TFCP2L1 | 9606.ENSP00000263707 | GO Function | GO:0000976 | Transcription cis-regulatory region binding |
| TFCP2L1 | 9606.ENSP00000263707 | GO Function | GO:0000977 | RNA polymerase II transcription regulatory region sequence-specific DNA binding |
| TFCP2L1 | 9606.ENSP00000263707 | GO Function | GO:0000978 | RNA polymerase II cis-regulatory region sequence-specific DNA binding |
| TFCP2L1 | 9606.ENSP00000263707 | GO Function | GO:0000981 | DNA-binding transcription factor activity, RNA polymerase II-specific |
| TFCP2L1 | 9606.ENSP00000263707 | GO Function | GO:0000987 | Cis-regulatory region sequence-specific DNA binding |
| TFCP2L1 | 9606.ENSP00000263707 | GO Function | GO:0001067 | Transcription regulatory region nucleic acid binding |
| TFCP2L1 | 9606.ENSP00000263707 | GO Function | GO:0001216 | DNA-binding transcription activator activity |
| TFCP2L1 | 9606.ENSP00000263707 | GO Function | GO:0001228 | DNA-binding transcription activator activity, RNA polymerase II-specific |

| | | | | |
|---|---|---|---|---|
| TFCP2L1 | 9606.ENSP00000263707 | GO Function | GO:0003676 | Nucleic acid binding |
| TFCP2L1 | 9606.ENSP00000263707 | GO Function | GO:0003677 | DNA binding |
| TFCP2L1 | 9606.ENSP00000263707 | GO Function | GO:0003690 | Double-stranded DNA binding |
| TFCP2L1 | 9606.ENSP00000263707 | GO Function | GO:0003700 | DNA-binding transcription factor activity |
| TFCP2L1 | 9606.ENSP00000263707 | GO Function | GO:0005488 | Binding |
| TFCP2L1 | 9606.ENSP00000263707 | GO Function | GO:0043565 | Sequence-specific DNA binding |
| TFCP2L1 | 9606.ENSP00000263707 | GO Function | GO:0097159 | Organic cyclic compound binding |
| TFCP2L1 | 9606.ENSP00000263707 | GO Function | GO:0140110 | Transcription regulator activity |
| TFCP2L1 | 9606.ENSP00000263707 | GO Function | GO:1901363 | Heterocyclic compound binding |
| TFCP2L1 | 9606.ENSP00000263707 | GO Function | GO:1990837 | Sequence-specific double-stranded DNA binding |
| TFCP2L1 | 9606.ENSP00000263707 | GO Component | GO:0000785 | Chromatin |
| TFCP2L1 | 9606.ENSP00000263707 | GO Component | GO:0005622 | Intracellular anatomical structure |
| TFCP2L1 | 9606.ENSP00000263707 | GO Component | GO:0005634 | Nucleus |
| TFCP2L1 | 9606.ENSP00000263707 | GO Component | GO:0005694 | Chromosome |
| TFCP2L1 | 9606.ENSP00000263707 | GO Component | GO:0005737 | Cytoplasm |
| TFCP2L1 | 9606.ENSP00000263707 | GO Component | GO:0016020 | Membrane |
| TFCP2L1 | 9606.ENSP00000263707 | GO Component | GO:0043226 | Organelle |
| TFCP2L1 | 9606.ENSP00000263707 | GO Component | GO:0043227 | Membrane-bounded organelle |
| TFCP2L1 | 9606.ENSP00000263707 | GO Component | GO:0043228 | Non-membrane-bounded organelle |
| TFCP2L1 | 9606.ENSP00000263707 | GO Component | GO:0043229 | Intracellular organelle |
| TFCP2L1 | 9606.ENSP00000263707 | GO Component | GO:0043231 | Intracellular membrane-bounded organelle |
| TFCP2L1 | 9606.ENSP00000263707 | GO Component | GO:0043232 | Intracellular non-membrane-bounded organelle |
| TFCP2L1 | 9606.ENSP00000263707 | GO Component | GO:0110165 | Cellular anatomical entity |
| TFCP2L1 | 9606.ENSP00000263707 | STRING clusters | CL:20713 | Mixed, incl. Transcription coregulator binding, and DNA-binding transcription repressor activity |
| TFCP2L1 | 9606.ENSP00000263707 | STRING clusters | CL:20967 | Mixed, incl. Transcriptional regulation of pluripotent stem cells, and Zinc finger, CXXC-type |
| TFCP2L1 | 9606.ENSP00000263707 | STRING clusters | CL:20969 | Mixed, incl. Transcriptional regulation of pluripotent stem cells, and 5-methylcytosine catabolic process |
| TFCP2L1 | 9606.ENSP00000263707 | STRING clusters | CL:20971 | Mixed, incl. POU5F1 (OCT4), SOX2, NANOG activate genes related to proliferation, and Ureteric peristalsis |
| TFCP2L1 | 9606.ENSP00000263707 | TISSUES | BTO:0000000 | Tissues, cell types and enzyme sources |
| TFCP2L1 | 9606.ENSP00000263707 | TISSUES | BTO:0000042 | Animal |
| TFCP2L1 | 9606.ENSP00000263707 | TISSUES | BTO:0000174 | Embryonic structure |
| TFCP2L1 | 9606.ENSP00000263707 | TISSUES | BTO:0000284 | Organism form |

| | | | | |
|---|---|---|---|---|
| TFCP2L1 | 9606.ENSP00000263707 | TISSUES | BTO:0001489 | Whole body |
| TFCP2L1 | 9606.ENSP00000263707 | Monarch | EFO:0004303 | Vital signs |
| TFCP2L1 | 9606.ENSP00000263707 | Monarch | EFO:0004305 | Erythrocyte count |
| TFCP2L1 | 9606.ENSP00000263707 | Monarch | EFO:0004306 | Erythrocyte indices |
| TFCP2L1 | 9606.ENSP00000263707 | Monarch | EFO:0004325 | Blood pressure |
| TFCP2L1 | 9606.ENSP00000263707 | Monarch | EFO:0004348 | Hematocrit |
| TFCP2L1 | 9606.ENSP00000263707 | Monarch | EFO:0004503 | Hematological measurement |
| TFCP2L1 | 9606.ENSP00000263707 | Monarch | EFO:0004509 | Hemoglobin measurement |
| TFCP2L1 | 9606.ENSP00000263707 | Monarch | EFO:0004518 | Creatinine measurement |
| TFCP2L1 | 9606.ENSP00000263707 | Monarch | EFO:0004531 | Urate measurement |
| TFCP2L1 | 9606.ENSP00000263707 | Monarch | EFO:0004586 | Complete blood cell count |
| TFCP2L1 | 9606.ENSP00000263707 | Monarch | EFO:0004617 | Cystatin C measurement |
| TFCP2L1 | 9606.ENSP00000263707 | Monarch | EFO:0004725 | Metabolite measurement |
| TFCP2L1 | 9606.ENSP00000263707 | Monarch | EFO:0004741 | Blood urea nitrogen measurement |
| TFCP2L1 | 9606.ENSP00000263707 | Monarch | EFO:0004742 | Renal system measurement |
| TFCP2L1 | 9606.ENSP00000263707 | Monarch | EFO:0004747 | Protein measurement |
| TFCP2L1 | 9606.ENSP00000263707 | Monarch | EFO:0005047 | Erythrocyte measurement |
| TFCP2L1 | 9606.ENSP00000263707 | Monarch | EFO:0005208 | Glomerular filtration rate |
| TFCP2L1 | 9606.ENSP00000263707 | Monarch | EFO:0006335 | Systolic blood pressure |
| TFCP2L1 | 9606.ENSP00000263707 | Monarch | EFO:0007978 | Red blood cell density measurement |
| TFCP2L1 | 9606.ENSP00000263707 | Monarch | EFO:0009283 | Potassium measurement |
| TFCP2L1 | 9606.ENSP00000263707 | Monarch | EFO:0009795 | Serum urea measurement |
| TFCP2L1 | 9606.ENSP00000263707 | UniProt Keywords | KW-0238 | DNA-binding |
| TFCP2L1 | 9606.ENSP00000263707 | UniProt Keywords | KW-0539 | Nucleus |
| TFCP2L1 | 9606.ENSP00000263707 | UniProt Keywords | KW-0804 | Transcription |
| TFCP2L1 | 9606.ENSP00000263707 | UniProt Keywords | KW-0805 | Transcription regulation |
| TFCP2L1 | 9606.ENSP00000263707 | Pfam | PF04516 | CP2 transcription factor |
| TFCP2L1 | 9606.ENSP00000263707 | InterPro | IPR007604 | CP2 transcription factor |
| TFCP2L1 | 9606.ENSP00000263707 | InterPro | IPR013761 | Sterile alpha motif/pointed domain superfamily |
| TFCP2L1 | 9606.ENSP00000263707 | InterPro | IPR037598 | TFCP2L1, SAM domain |
| TFCP2L1 | 9606.ENSP00000263707 | InterPro | IPR040167 | Transcription factor CP2-like |
| TFCP2L1 | 9606.ENSP00000263707 | InterPro | IPR041418 | SAM domain |
| TRABD2A | 9606.ENSP00000387075 | GO Process | GO:0006508 | Proteolysis |
| TRABD2A | 9606.ENSP00000387075 | GO Process | GO:0006807 | Nitrogen compound metabolic process |
| TRABD2A | 9606.ENSP00000387075 | GO Process | GO:0007154 | Cell communication |

| | | | | |
|---|---|---|---|---|
| TRABD2A | 9606.ENSP00000387075 | GO Process | GO:0007165 | Signal transduction |
| TRABD2A | 9606.ENSP00000387075 | GO Process | GO:0007166 | Cell surface receptor signaling pathway |
| TRABD2A | 9606.ENSP00000387075 | GO Process | GO:0007267 | Cell-cell signaling |
| TRABD2A | 9606.ENSP00000387075 | GO Process | GO:0008152 | Metabolic process |
| TRABD2A | 9606.ENSP00000387075 | GO Process | GO:0009893 | Positive regulation of metabolic process |
| TRABD2A | 9606.ENSP00000387075 | GO Process | GO:0009966 | Regulation of signal transduction |
| TRABD2A | 9606.ENSP00000387075 | GO Process | GO:0009968 | Negative regulation of signal transduction |
| TRABD2A | 9606.ENSP00000387075 | GO Process | GO:0009987 | Cellular process |
| TRABD2A | 9606.ENSP00000387075 | GO Process | GO:0010604 | Positive regulation of macromolecule metabolic process |
| TRABD2A | 9606.ENSP00000387075 | GO Process | GO:0010646 | Regulation of cell communication |
| TRABD2A | 9606.ENSP00000387075 | GO Process | GO:0010648 | Negative regulation of cell communication |
| TRABD2A | 9606.ENSP00000387075 | GO Process | GO:0016055 | Wnt signaling pathway |
| TRABD2A | 9606.ENSP00000387075 | GO Process | GO:0019222 | Regulation of metabolic process |
| TRABD2A | 9606.ENSP00000387075 | GO Process | GO:0019538 | Protein metabolic process |
| TRABD2A | 9606.ENSP00000387075 | GO Process | GO:0023051 | Regulation of signaling |
| TRABD2A | 9606.ENSP00000387075 | GO Process | GO:0023052 | Signaling |
| TRABD2A | 9606.ENSP00000387075 | GO Process | GO:0023057 | Negative regulation of signaling |
| TRABD2A | 9606.ENSP00000387075 | GO Process | GO:0030111 | Regulation of Wnt signaling pathway |
| TRABD2A | 9606.ENSP00000387075 | GO Process | GO:0030178 | Negative regulation of Wnt signaling pathway |
| TRABD2A | 9606.ENSP00000387075 | GO Process | GO:0031334 | Positive regulation of protein-containing complex assembly |
| TRABD2A | 9606.ENSP00000387075 | GO Process | GO:0031399 | Regulation of protein modification process |
| TRABD2A | 9606.ENSP00000387075 | GO Process | GO:0031401 | Positive regulation of protein modification process |
| TRABD2A | 9606.ENSP00000387075 | GO Process | GO:0032502 | Developmental process |
| TRABD2A | 9606.ENSP00000387075 | GO Process | GO:0043170 | Macromolecule metabolic process |
| TRABD2A | 9606.ENSP00000387075 | GO Process | GO:0043254 | Regulation of protein-containing complex assembly |
| TRABD2A | 9606.ENSP00000387075 | GO Process | GO:0044087 | Regulation of cellular component biogenesis |
| TRABD2A | 9606.ENSP00000387075 | GO Process | GO:0044089 | Positive regulation of cellular component biogenesis |
| TRABD2A | 9606.ENSP00000387075 | GO Process | GO:0044238 | Primary metabolic process |
| TRABD2A | 9606.ENSP00000387075 | GO Process | GO:0048518 | Positive regulation of biological process |
| TRABD2A | 9606.ENSP00000387075 | GO Process | GO:0048519 | Negative regulation of biological process |
| TRABD2A | 9606.ENSP00000387075 | GO Process | GO:0048522 | Positive regulation of cellular process |
| TRABD2A | 9606.ENSP00000387075 | GO Process | GO:0048523 | Negative regulation of cellular process |
| TRABD2A | 9606.ENSP00000387075 | GO Process | GO:0048583 | Regulation of response to stimulus |
| TRABD2A | 9606.ENSP00000387075 | GO Process | GO:0048585 | Negative regulation of response to stimulus |
| TRABD2A | 9606.ENSP00000387075 | GO Process | GO:0048856 | Anatomical structure development |

| | | | | |
|---|---|---|---|---|
| TRABD2A | 9606.ENSP00000387075 | GO Process | GO:0050789 | Regulation of biological process |
| TRABD2A | 9606.ENSP00000387075 | GO Process | GO:0050794 | Regulation of cellular process |
| TRABD2A | 9606.ENSP00000387075 | GO Process | GO:0050896 | Response to stimulus |
| TRABD2A | 9606.ENSP00000387075 | GO Process | GO:0051128 | Regulation of cellular component organization |
| TRABD2A | 9606.ENSP00000387075 | GO Process | GO:0051130 | Positive regulation of cellular component organization |
| TRABD2A | 9606.ENSP00000387075 | GO Process | GO:0051171 | Regulation of nitrogen compound metabolic process |
| TRABD2A | 9606.ENSP00000387075 | GO Process | GO:0051173 | Positive regulation of nitrogen compound metabolic process |
| TRABD2A | 9606.ENSP00000387075 | GO Process | GO:0051246 | Regulation of protein metabolic process |
| TRABD2A | 9606.ENSP00000387075 | GO Process | GO:0051247 | Positive regulation of protein metabolic process |
| TRABD2A | 9606.ENSP00000387075 | GO Process | GO:0051716 | Cellular response to stimulus |
| TRABD2A | 9606.ENSP00000387075 | GO Process | GO:0060255 | Regulation of macromolecule metabolic process |
| TRABD2A | 9606.ENSP00000387075 | GO Process | GO:0060322 | Head development |
| TRABD2A | 9606.ENSP00000387075 | GO Process | GO:0065007 | Biological regulation |
| TRABD2A | 9606.ENSP00000387075 | GO Process | GO:0071704 | Organic substance metabolic process |
| TRABD2A | 9606.ENSP00000387075 | GO Process | GO:0080090 | Regulation of primary metabolic process |
| TRABD2A | 9606.ENSP00000387075 | GO Process | GO:0198738 | Cell-cell signaling by wnt |
| TRABD2A | 9606.ENSP00000387075 | GO Process | GO:1901564 | Organonitrogen compound metabolic process |
| TRABD2A | 9606.ENSP00000387075 | GO Process | GO:1904806 | Regulation of protein oxidation |
| TRABD2A | 9606.ENSP00000387075 | GO Process | GO:1904808 | Positive regulation of protein oxidation |
| TRABD2A | 9606.ENSP00000387075 | GO Process | GO:1905114 | Cell surface receptor signaling pathway involved in cell-cell signaling |
| TRABD2A | 9606.ENSP00000387075 | GO Function | GO:0003824 | Catalytic activity |
| TRABD2A | 9606.ENSP00000387075 | GO Function | GO:0004175 | Endopeptidase activity |
| TRABD2A | 9606.ENSP00000387075 | GO Function | GO:0004222 | Metalloendopeptidase activity |
| TRABD2A | 9606.ENSP00000387075 | GO Function | GO:0005488 | Binding |
| TRABD2A | 9606.ENSP00000387075 | GO Function | GO:0005515 | Protein binding |
| TRABD2A | 9606.ENSP00000387075 | GO Function | GO:0008233 | Peptidase activity |
| TRABD2A | 9606.ENSP00000387075 | GO Function | GO:0008237 | Metallopeptidase activity |
| TRABD2A | 9606.ENSP00000387075 | GO Function | GO:0016787 | Hydrolase activity |
| TRABD2A | 9606.ENSP00000387075 | GO Function | GO:0017147 | Wnt-protein binding |
| TRABD2A | 9606.ENSP00000387075 | GO Function | GO:0043167 | Ion binding |
| TRABD2A | 9606.ENSP00000387075 | GO Function | GO:0043169 | Cation binding |
| TRABD2A | 9606.ENSP00000387075 | GO Function | GO:0046872 | Metal ion binding |
| TRABD2A | 9606.ENSP00000387075 | GO Function | GO:0140096 | Catalytic activity, acting on a protein |
| TRABD2A | 9606.ENSP00000387075 | GO Component | GO:0005886 | Plasma membrane |
| TRABD2A | 9606.ENSP00000387075 | GO Component | GO:0005887 | Integral component of plasma membrane |

| | | | | |
|---|---|---|---|---|
| TRABD2A | 9606.ENSP00000387075 | GO Component | GO:0016020 | Membrane |
| TRABD2A | 9606.ENSP00000387075 | GO Component | GO:0016021 | Integral component of membrane |
| TRABD2A | 9606.ENSP00000387075 | GO Component | GO:0031090 | Organelle membrane |
| TRABD2A | 9606.ENSP00000387075 | GO Component | GO:0031224 | Intrinsic component of membrane |
| TRABD2A | 9606.ENSP00000387075 | GO Component | GO:0031226 | Intrinsic component of plasma membrane |
| TRABD2A | 9606.ENSP00000387075 | GO Component | GO:0031300 | Intrinsic component of organelle membrane |
| TRABD2A | 9606.ENSP00000387075 | GO Component | GO:0031301 | Integral component of organelle membrane |
| TRABD2A | 9606.ENSP00000387075 | GO Component | GO:0043226 | Organelle |
| TRABD2A | 9606.ENSP00000387075 | GO Component | GO:0043227 | Membrane-bounded organelle |
| TRABD2A | 9606.ENSP00000387075 | GO Component | GO:0071944 | Cell periphery |
| TRABD2A | 9606.ENSP00000387075 | GO Component | GO:0110165 | Cellular anatomical entity |
| TRABD2A | 9606.ENSP00000387075 | STRING clusters | CL:22079 | Mixed, incl. APCDD1 domain, and Olfactory learning |
| TRABD2A | 9606.ENSP00000387075 | TISSUES | BTO:0000000 | Tissues, cell types and enzyme sources |
| TRABD2A | 9606.ENSP00000387075 | TISSUES | BTO:0000042 | Animal |
| TRABD2A | 9606.ENSP00000387075 | TISSUES | BTO:0000058 | Alimentary canal |
| TRABD2A | 9606.ENSP00000387075 | TISSUES | BTO:0000269 | Colon |
| TRABD2A | 9606.ENSP00000387075 | TISSUES | BTO:0000511 | Gastrointestinal tract |
| TRABD2A | 9606.ENSP00000387075 | TISSUES | BTO:0000648 | Intestine |
| TRABD2A | 9606.ENSP00000387075 | TISSUES | BTO:0000706 | Large intestine |
| TRABD2A | 9606.ENSP00000387075 | TISSUES | BTO:0001489 | Whole body |
| TRABD2A | 9606.ENSP00000387075 | TISSUES | BTO:0001491 | Viscus |
| TRABD2A | 9606.ENSP00000387075 | TISSUES | BTO:0001613 | Colorectum |
| TRABD2A | 9606.ENSP00000387075 | COMPARTMENTS | GOCC:0005886 | Plasma membrane |
| TRABD2A | 9606.ENSP00000387075 | COMPARTMENTS | GOCC:0005887 | Integral component of plasma membrane |
| TRABD2A | 9606.ENSP00000387075 | COMPARTMENTS | GOCC:0016020 | Membrane |
| TRABD2A | 9606.ENSP00000387075 | COMPARTMENTS | GOCC:0016021 | Integral component of membrane |
| TRABD2A | 9606.ENSP00000387075 | COMPARTMENTS | GOCC:0031090 | Organelle membrane |
| TRABD2A | 9606.ENSP00000387075 | COMPARTMENTS | GOCC:0031224 | Intrinsic component of membrane |
| TRABD2A | 9606.ENSP00000387075 | COMPARTMENTS | GOCC:0031226 | Intrinsic component of plasma membrane |
| TRABD2A | 9606.ENSP00000387075 | COMPARTMENTS | GOCC:0031300 | Intrinsic component of organelle membrane |
| TRABD2A | 9606.ENSP00000387075 | COMPARTMENTS | GOCC:0031301 | Integral component of organelle membrane |
| TRABD2A | 9606.ENSP00000387075 | COMPARTMENTS | GOCC:0043226 | Organelle |
| TRABD2A | 9606.ENSP00000387075 | COMPARTMENTS | GOCC:0043227 | Membrane-bounded organelle |
| TRABD2A | 9606.ENSP00000387075 | COMPARTMENTS | GOCC:0071944 | Cell periphery |
| TRABD2A | 9606.ENSP00000387075 | COMPARTMENTS | GOCC:0110165 | Cellular anatomical entity |

| | | | | |
|---|---|---|---|---|
| TRABD2A | 9606.ENSP00000387075 | Monarch | EFO:0004298 | Cardiovascular measurement |
| TRABD2A | 9606.ENSP00000387075 | Monarch | EFO:0004311 | Heart function measurement |
| TRABD2A | 9606.ENSP00000387075 | Monarch | EFO:0004747 | Protein measurement |
| TRABD2A | 9606.ENSP00000387075 | Monarch | EFO:0005043 | Cardiac troponin T measurement |
| TRABD2A | 9606.ENSP00000387075 | Monarch | EFO:0005278 | Cardiovascular disease biomarker measurement |
| TRABD2A | 9606.ENSP00000387075 | UniProt Keywords | KW-0025 | Alternative splicing |
| TRABD2A | 9606.ENSP00000387075 | UniProt Keywords | KW-0325 | Glycoprotein |
| TRABD2A | 9606.ENSP00000387075 | UniProt Keywords | KW-0378 | Hydrolase |
| TRABD2A | 9606.ENSP00000387075 | UniProt Keywords | KW-0472 | Membrane |
| TRABD2A | 9606.ENSP00000387075 | UniProt Keywords | KW-0479 | Metal-binding |
| TRABD2A | 9606.ENSP00000387075 | UniProt Keywords | KW-0482 | Metalloprotease |
| TRABD2A | 9606.ENSP00000387075 | UniProt Keywords | KW-0645 | Protease |
| TRABD2A | 9606.ENSP00000387075 | UniProt Keywords | KW-0732 | Signal |
| TRABD2A | 9606.ENSP00000387075 | UniProt Keywords | KW-0812 | Transmembrane |
| TRABD2A | 9606.ENSP00000387075 | UniProt Keywords | KW-0879 | Wnt signaling pathway |
| TRABD2A | 9606.ENSP00000387075 | UniProt Keywords | KW-1003 | Cell membrane |
| TRABD2A | 9606.ENSP00000387075 | UniProt Keywords | KW-1133 | Transmembrane helix |
| TRABD2A | 9606.ENSP00000387075 | InterPro | IPR002816 | TraB/PrgY/GumN family |
| TRABD2A | 9606.ENSP00000387075 | InterPro | IPR040230 | Protease TIKI1/2-like |
